# α-Synuclein aggregates inhibit ESCRT-III through sequestration and collateral degradation

**DOI:** 10.1101/2025.01.13.632710

**Authors:** Cole S Sitron, Victoria A Trinkaus, Ana Galesic, Maximilian Garhammer, Patricia Yuste Checa, Ulrich Dransfeld, Dennis Feigenbutz, Jiuchun Zhang, Irina Dudanova, J Wade Harper, F Ulrich Hartl

**Affiliations:** Department of Cellular Biochemistry, Max Planck Institute of Biochemistry, Martinsried, Germany; Aligning Science Across Parkinson’s (ASAP) Collaborative Research Network, Chevy Chase, Maryland, USA; Munich Cluster for Systems Neurology (SyNergy), Munich, Germany; Department of Cell Biology, Harvard Medical School, Boston, MA, USA; Molecular Neurodegeneration Group, Max Planck Institute for Biological Intelligence, Martinsried, Germany; Department of Molecules – Signaling – Development, Max Planck Institute for Biological Intelligence, Martinsried, Germany; Center for Anatomy, Faculty of Medicine and University Hospital Cologne, University of Cologne, Cologne, Germany

**Keywords:** α-Synuclein, CHMP2B, ESCRT-III, ESCRT, protein aggregate spreading, aggregation, sequestration, lysosome, proteostasis, Parkinson’s disease

## Abstract

α-Synuclein aggregation is a hallmark of Parkinson’s disease and related synucleinopathies. Extracellular α-synuclein fibrils enter naïve cells via endocytosis, followed by transit into the cytoplasm to seed endogenous α-synuclein aggregation. Intracellular aggregates sequester numerous proteins, including subunits of the ESCRT-III system for endolysosome membrane repair, but the toxic effects of these events remain poorly understood. Using cellular models and in vitro reconstitution, we found that α-synuclein fibrils interact with an α-helix common to ESCRT-III proteins. This interaction results in sequestration of ESCRT-III subunits and triggers their proteasomal destruction in a process of “collateral degradation.” These twin mechanisms deplete the available ESCRT-III pool, initiating a toxic feedback loop. The ensuing loss of ESCRT function compromises endolysosome membranes, thereby facilitating escape of aggregate seeds into the cytoplasm, which in turn increases aggregation and ESCRT-III sequestration. We suggest that collateral degradation and triggering of self-perpetuating systems could be general mechanisms of sequestration-induced proteotoxicity.

**HIGHLIGHTS:**

- α-Synuclein fibrils bind and sequester ESCRT-III endolysosome repair proteins
- An α-helical segment common to ESCRT-III mediates fibril-selective interaction
- Fibril-bound ESCRT-III subunits undergo “collateral degradation” via the proteasome
- ESCRT-III depletion damages endolysosomes and worsens α-synuclein aggregation

## INTRODUCTION

Progression of many neurodegenerative diseases, including Alzheimer’s (AD) and Parkinson’s disease (PD), is associated with the spreading of aggregate pathology throughout interconnected brain regions^1–5^. Aggregation initiates in an area of the nervous system that is characteristic to each disease. For instance, α-synuclein (α-syn) aggregates in PD arise in the brainstem or peripheral nervous system before spreading to higher brain regions, correlating with progression from a prodromal stage to motor deficits and eventually to dementia^6–10^.

The molecular mechanism underlying spreading is thought to be prion-like aggregate “seeding.” In this process, a fibrillar protein seed transfers from one cell to another, where it templates misfolding and aggregation of its monomeric counterparts^11–14^. Seeds leave donor cells via several mechanisms, including release into the extracellular milieu^15–19^. Highlighting the importance of this exocytic route, diagnostics detecting α-syn aggregate seeds in cerebrospinal fluid show promise in identifying PD in its early stages^20–23^. Aggregate seeds released as naked proteins can enter acceptor cells by endocytosis, arriving in the membrane-bound organelles of the endolysosomal system^12,24–30^. Confinement inside endolysosomes would restrict seeds from accessing the cytoplasmic pool of monomeric protein. However, endolysosomal membrane ruptures permit seeds to escape and trigger aggregation^26,31–35^.

The ESCRT (endosomal sorting complexes required for transport) machinery, a series of cytosolic protein complexes (ESCRT-0, -I, -II and -III) involved in membrane fission events^36–46^, plays a critical role in repairing endolysosomal membrane damage, thereby preventing leakage of aggregate seeds into the cytoplasm. The ESCRT-III complex consists of 12 proteins in humans, including three groups of paralogs with at least partially overlapping function. ESCRT-III proteins assemble into spiral spring-shaped hetero-oligomers and function to invaginate the membrane they bind^31,47–55^. The AAA ATPase VPS4 then interacts with ESCRT-III and extracts subunits to disassemble the oligomer, severing the invaginated membrane to form an intralumenal vesicle^56–64^. In the context of membrane repair, the membrane constriction driven by ESCRT-III oligomerization at the wound site brings the areas of membrane at the edge of the lesion into contact, thereby resealing the damaged membrane^40,42,65,66^. Underscoring the importance of ESCRT’s role in maintaining nervous system health, mutations in the ESCRT-III protein CHMP2B cause diseases of the Amyotrophic Lateral Sclerosis– Frontotemporal Dementia spectrum (FTD-ALS) ^67–69^ and have additionally been detected in patients suffering from Lewy Body Dementia, a synucleinopathy related to PD^70^. Although CHMP2B has a paralog, CHMP2A, it is interesting to note that only CHMP2B mutation has thus far been implicated as a genetic risk factor for neurodegenerative disease.

When neurodegeneration-associated aggregation proceeds, the resultant aggregates contain numerous macromolecules in addition to the primary disease protein. Lewy Bodies in PD, a well characterized form of aggregate deposit, contain α-syn as well as organelles, membranes, and over 100 other proteins^8,71,72^. Researchers have long posited that sequestration of bystander proteins by aggregates depletes the available pool of these proteins, resulting in widespread loss-of-function and subsequent proteostasis collapse^73–75^. However, while this “*trans*-acting loss-of-function” is a commonly-invoked mechanism of aggregate toxicity, demonstrations of pathomechanistic relevance of such processes are rare^76–78^ and mostly concern chaperones or ubiquitin-binding proteins^79–84^.

Interestingly, the ESCRT-III protein CHMP2B co-localizes with aggregates of Amyloid beta (Aβ), tau, and α-syn in patient brains and cellular models^85–91^. Despite the clear association, whether this co-localization reflects *trans*-acting loss-of-function by sequestration remains unclear. Here we analyzed the α-syn-CHMP2B interaction to determine whether and how it leads to collapse of proteostasis. To this end, we developed tools to control α-syn aggregation in a mammalian cell culture model and to measure three processes underlying α-syn cellular effects – disruption of ESCRT-III function, endolysosome breakage, and fibril leakage into the cytoplasm. We find that seeded α-syn aggregates sequester CHMP2B and several other ESCRT-III proteins. Notably, a central α-helix of CHMP2B binds preferentially to the fibrillar aggregated state rather than monomeric α-syn. This aberrant interaction confines ESCRT-III proteins within α-syn aggregates and, importantly, targets some ESCRT-III subunits for “collateral degradation” via the proteasome. In combination, these two effects lead to a critical depletion of available ESCRT-III subunits and loss of ESCRT function. As a result, α-syn aggregation compromises endolysosomal membrane integrity. This in turn causes leakage of exogenous α-syn fibrils into the cytoplasm, forming a positive feedback loop between α-syn aggregation and ESCRT-III loss-of-function. Proteopathic aggregates thus exert *trans*-acting loss-of-function by both sequestering other cellular proteins and targeting these proteins for collateral degradation. In the case of transmissible intracellular aggregates, this *trans*-acting loss-of-function can collapse endolysosomal homeostasis by sequestering membrane repair proteins that otherwise limit seeded aggregation and combat aggregate spreading.

## RESULTS

### α-Syn aggregates sequester ESCRT-III proteins

To study the effect of α-syn aggregation on ESCRT-III function, we developed a HEK cellular model that permitted control of the α-syn aggregation state. HEK cells express only low levels of endogenous α-syn^92^, so we expressed the highly aggregation-prone α-syn A53T mutant^30,93^ with a doxycycline (Dox)-repressible promoter as a source of soluble α-syn that could be templated to aggregate (Figure 1A). The protein was not retained on a cellulose acetate filter, suggesting that it was soluble (Figure 1B). We then exposed these cells to exogenous α-syn A53T pre-formed fibrils (PFFs)^8,13,14,72,94,95^ as well as lipofectamine (“Lpf”), a commonly-used transfection reagent that enables fibrils to access the cytoplasm^72,96–101^. The combination of PFFs and Lpf led to the accumulation of high molecular weight α-syn species on SDS-PAGE and retention of Ser129-phosphorylated α-syn (pS129 α-syn) on a cellulose acetate filter after two days (Figure 1B), indicative of insoluble α-syn aggregates^102^. Blocking intracellular α-syn expression by culturing with Dox abolished production of both high molecular weight and filter-retained α-syn (Figure 1B), indicating that aggregates contained endogenous α-syn. For simplicity, the combination of conditions resulting in insoluble α-syn production (PFF and Lpf co-treatment in the absence of Dox) will be referred to as “induction of aggregation.” When visualized by microscopy, induction of aggregation produced inclusions that were positive for pS129 α-syn, Lys48-linked ubiquitin, and p62 (also called SQSTM1) (Figure S1A), markers of pathologic α-syn aggregation^72,102–105^. Thus, the HEK system permits the formation of α-syn aggregates that bear pathologic hallmarks under conditions where intracellular α-syn is expressed and PFFs reach the cytoplasm.

**Figure 1.**
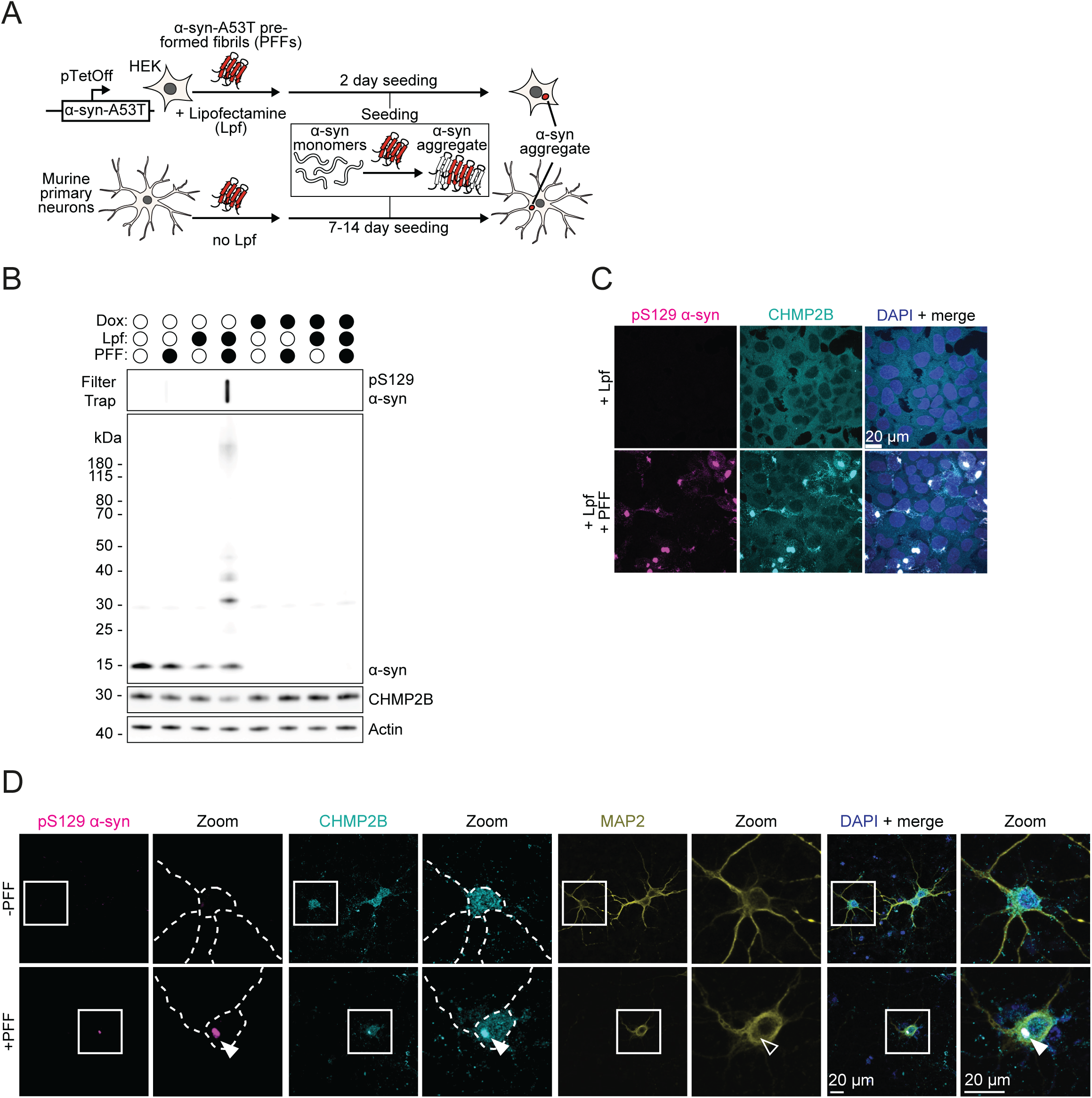
α-Syn aggregates sequester ESCRT-III proteins. (A) Experimental workflow used to induce α-syn aggregation in the cellular systems used in this study. α-syn, α-synuclein A53T mutant. PFFs, α-syn-A53T pre-formed fibrils. Lpf, lipofectamine. (B) Representative immunoblot and filter trap of lysates from HEK cells expressing α-syn under a tetracycline-repressible promoter. These cells were exposed to doxycycline (Dox) for 10 days to turn off the promoter driving α-syn expression prior to 2-day treatment with PFFs to template intracellular α-syn aggregation and/or lipofectamine (Lpf) to allow the PFFs to access the cytoplasm. The filter was stained with an antibody against Ser129-phosphorylated (pS129) α-syn and the immunoblot was stained with anti-α-syn, anti-CHMP2B, and anti-β-actin antibodies. β-actin served as a loading control (n=3). (C) Representative immunofluorescence microscopy of HEK cells expressing α-syn, before and after induction of α-syn aggregation as described in (A). Cells are stained for pS129 α-syn (a marker for pathologic α-syn aggregation) and the ESCRT-III protein CHMP2B (Scale bar, 20 μm). (D) Representative immunofluorescence microscopy of primary mouse neurons after inducing α-syn aggregation as described in (A). Neurons are stained for pS129 α-syn, CHMP2B, and the neuronal cytoskeletal marker MAP2 (Scale bar, 20 μm). Higher magnification images on the right show the areas marked by the white boxes. Dashed lines outline the contours of the neurons. Arrowheads mark pS129 α-syn inclusions. See also Figure S1.

Induction of aggregation markedly altered the localization of the ESCRT-III protein CHMP2B in HEK cells. While control cells featured a diffuse CHMP2B distribution, CHMP2B mainly localized to pS129 α-syn inclusions in cells with aggregates (Figure 1C). α-Syn inclusions with CHMP2B sequestration also formed without Lpf, provided the cells were exposed to PFFs for 6 days (Figure S1B). Similarly, exposure of primary mouse neurons to PFFs for 14 days (without Lpf) resulted in the co-localization of CHMP2B with pS129 α-syn inclusions (Figures 1D and S1C). The clinical finding of CHMP2B co-localization with α-syn aggregates^86,87,89^ can therefore be reproduced in cell culture upon aggregate seeding with or without Lpf.

In addition to co-localizing with α-syn inclusions, we observed that CHMP2B decreased in abundance after induction of aggregation (Figures 1B and S1D). CHMP2B levels also declined in primary neurons treated with PFFs (Figure S1E). These results are consistent with observations in the brain of Lewy Body Dementia patients^86^ and will be addressed in depth below.

We next tested whether α-syn associated with other ESCRT-III subunits in addition to CHMP2B. Immunofluorescent staining revealed that CHMP2A (paralog of CHMP2B), CHMP3, and CHMP4B re-localized to α-syn aggregates as well, while co-localization with CHMP6 was not detectable (Figure S1F). This result implies that α-syn aggregation may sequester much of the ESCRT-III system and that the association of CHMP2B with α-syn aggregates can serve as a proxy for this effect. Given the strength of the clinical data on CHMP2B and its close connection to neurodegenerative disease^67–70,85–88,106,107^, this study will focus on CHMP2B and its paralog CHMP2A, though what we find for these proteins may also apply for other ESCRT-III subunits.

### An α-helical segment of CHMP2B mediates binding to α-syn fibrils

To determine whether the microscopic co-localization of α-syn aggregates and CHMP2B reflects a physical interaction, we immunoprecipitated CHMP2B from cells with and without α-syn aggregates. Without aggregation induction (Lpf only), the pulldown did not recover α-syn (Figure 2A). Immunoprecipitation from lysates of aggregation-induced cells, however, specifically co-purified high molecular weight detergent-resistant α-syn species along with CHMP2B (Figure 2A). We obtained similar results with α-syn aggregates generated without Lpf (Figure S2A).

**Figure 2.**
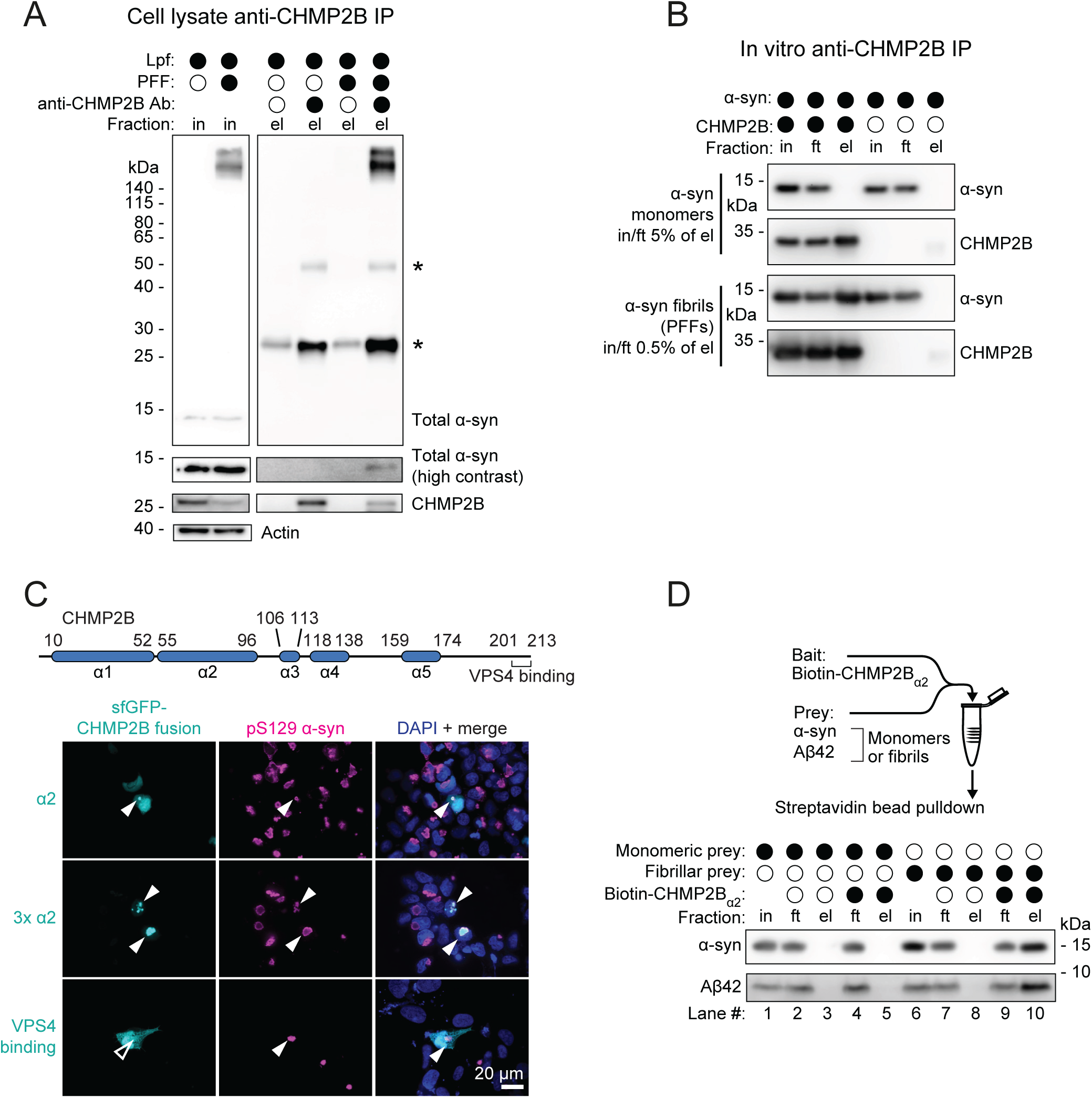
An α-helical segment of CHMP2B mediates binding to α-syn fibrils. (A) Representative immunoblot of an anti-CHMP2B immunoprecipitation from cells with and without α-syn aggregation induction. Proteins were detected using anti-CHMP2B, anti-α-syn, and anti-β-actin antibodies, with β-actin serving as a loading control (n=2). Asterisks mark non-specific bands that originate from the Protein G-IgG antibody complex used in the immunoprecipitation. IP, immunoprecipitation. In, input. El, eluate. (B) Representative anti-CHMP2B and anti-α-syn immunoblots of an in vitro co-immunoprecipitation of CHMP2B with either monomeric α-syn or PFFs, performed using an anti-CHMP2B antibody (n=3). Ft, flow-through. MW, molecular weight. (C) Determination of whether superfolder GFP (sfGFP) fusions with indicated CHMP2B regions (see secondary structure representation above) co-localize with α-syn aggregates. Representative immunofluorescence micrographs were stained for pS129 α-syn. Arrowheads denote the location of α-syn inclusions (Scale bar, 20 μm). (D) Representative immunoblots from α-syn and Aβ42 pulldowns in either their monomeric or fibrillar states, carried out in vitro with a synthetic biotinylated peptide spanning CHMP2B residues 55-96 (α2) conjugated to streptavidin resin (n=3). See also Figure S2.

Given that both α-syn and ESCRT-III proteins bind membranes^108–118^, the interaction between these species could be indirect and bridged by membranous organelles found in α-syn aggregates^71,72,119–122^. To address this possibility, we first assessed the ability of a membrane-binding deficient CHMP2B mutant (CHMP2B L4D F5D^123^) to associate with α-syn aggregates. Both wild-type (wt) and mutant CHMP2B co-localized with pS129 α-syn inclusions (Figure S2B), suggesting that the interaction is membrane independent. We further evaluated the interaction by in vitro co-immunoprecipitation of recombinant components using an anti-CHMP2B antibody (Figure 2B). Purified CHMP2B interacted with α-syn PFFs but not with monomers (Figure 2B). The CHMP2B-α-syn interaction is therefore direct and specific for fibrillar α-syn over soluble α-syn.

To map the interaction interface between α-syn and ESCRT-III proteins, we evaluated the ability of a series of CHMP2B mutants to associate with α-syn aggregates in cells. This experiment utilized membrane-binding deficient CHMP2B to eliminate possible confounding effects of the mutations on membrane binding. CHMP2B comprises five main structural helices as well as a small C-terminal helix that interacts with the ATPase VPS4^36–39,41,43^ (Figures S2C and 2C diagrams). We deleted these helices one at a time. While most mutants retained at least some ability to co-localize with α-syn aggregates, the mutant lacking the second helix (α2), spanning residues 55-96, no longer co-localized (“Δα2;” Figure S2C). Ablation of the region spanning α2 to α4 (residues 55-138) prevented co-localization with α-syn aggregates similarly to loss of α2 (“Δα2-4;” Figure S2C). To test whether α2 was sufficient for binding, we fused it to superfolder GFP (sfGFP). The sfGFP-CHMP2B_α2_ fusion co-localized with α-syn inclusions (Figure 2C). This co-localization was less pronounced than for endogenous CHMP2B (Figure 1C), perhaps because α2 represents a sufficient but partially truncated, and therefore lower affinity, α-syn binder than full-length CHMP2B. Indeed, expressing a triple tandem CHMP2B_α2_ repeat to increase avidity led to qualitatively more robust co-localization than sfGFP-CHMP2B_α2_ (Figure 2C). As a negative control, we fused sfGFP to the C-terminal VPS4-binding helix (residues 201-213) and observed no co-localization (Figure 2C). Thus, CHMP2B_α2_ is necessary and sufficient for the co-localization between CHMP2B and α-syn aggregates.

We then assessed whether CHMP2B_α2_ possesses the same fibril-discriminating properties as the full-length protein. We conjugated a biotinylated synthetic CHMP2B_α2_ peptide to streptavidin resin and evaluated its ability to pull down different fibrillar or monomeric proteins. α-Syn monomers failed to associate with both peptide-conjugated resin as well as an unconjugated control (Figure 2D; lanes 5 and 3). However, α-syn PFFs specifically bound CHMP2B_α2_ resin but not unconjugated control resin (Figure 2D; lanes 10 and 8). CHMP2B_α2_ therefore binds fibrillar α-syn but not monomeric α-syn, and this interaction is not an artifact of α-syn PFFs nonspecifically binding any protein (e.g. streptavidin). Given the presence of CHMP2B in several different neurodegeneration-associated aggregates^85–88,91^, we tested whether its α-syn-interacting helix could bind another amyloid protein, Aβ42. As for α-syn, CHMP2B_α2_ specifically bound fibrillar but not monomeric Aβ42 (Figure 2D; lanes 10 and 5).

All ESCRT-III proteins share this α2 motif, including the aggregate-associated CHMP2A, CHMP3 and CHMP4B subunits (Figure S1F), as evidenced by an alignment of human CHMP protein α2 regions (Figure S2D). These sequences display similar physico-chemical properties: they are enriched in basic and aliphatic amino acids, notably methionine (Figure S2E). The basic amino acids mostly lie at the start of the helix, giving the N-terminus a positive charge (Figure S2D). The C-terminus contains clusters of strongly hydrophobic amino acids, spaced with a 7-residue periodicity, with additional hydrophobic residues located at the midpoints between these clusters (Figure S2F). CHMP2B_α2_ also possesses this pattern, albeit shifted by 3 amino acids C-terminally relative to the other sequences (Figure S2D). The spacing of these hydrophobic residues suggests the presence of a hydrophobic face on the α-helix that could presumably facilitate binding to amyloid fibrils (Figure S2G).

In summary, α-syn aggregates in cell culture specifically sequester CHMP2B and other ESCRT-III components, based on a direct interaction with aggregated α-syn. CHMP2B interacts with fibrillar α-syn using its α2. This same region also specifically binds Aβ42 fibrils, suggesting that α2 of ESCRT-III proteins might recognize amyloid folds more broadly.

### α-Syn aggregation compromises ESCRT-III function

Does binding of CHMP2B and other ESCRT-III proteins to α-syn aggregates affect ESCRT function? To measure ESCRT activity, we took advantage of a recent finding that several mammalian lysosome membrane proteins undergo rapid ESCRT-III-dependent turnover. One such protein is the E3 ubiquitin ligase RNF152, which autoubiquitinates to trigger recruitment of ESCRT-III proteins and subsequent degradation after sorting into the lysosomal lumen^44^. We adapted this process into an assay for ESCRT activity in the α-syn expressing HEK cell line by introducing a bicistronic construct encoding an EGFP-RNF152 reporter as well as mScarlet linked via an IRES (Figure 3A), the latter serving as an internal control for translation of the reporter. When monitored by flow cytometry (Figure S3A), the EGFP/mScarlet ratio thereby reflects the stability of the reporter, increasing with inhibition of EGFP-RNF152 degradation. Small molecule inhibitors targeting E1 ubiquitin ligases, lysosome acidification, and lysosomal proteases stabilized the reporter (Figure S3B). As expected, the EGFP signal from the reporter co-localized with the lysosomal marker TMEM192 (Figure S3C).

**Figure 3.**
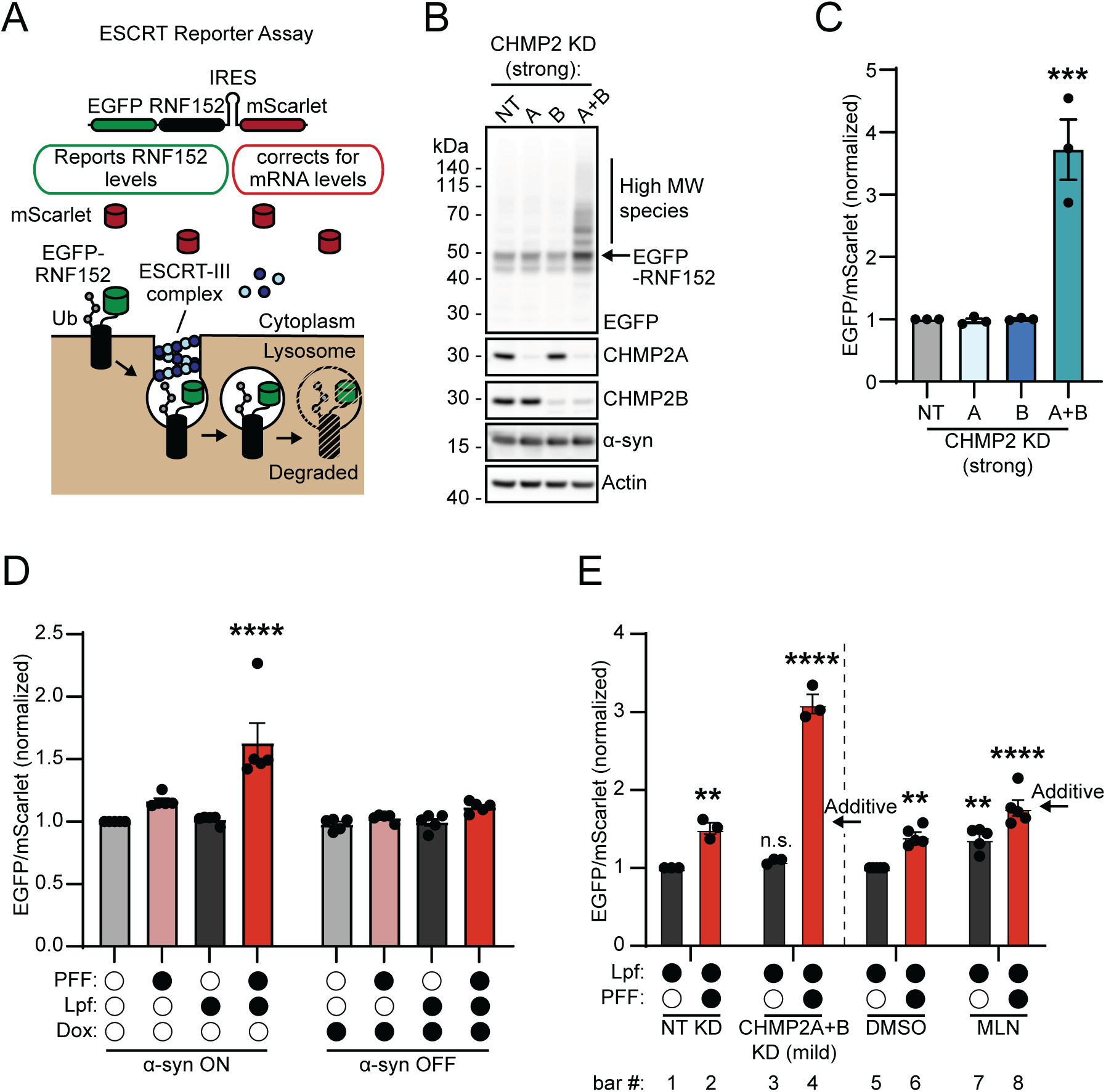
α-Syn aggregation inhibits ESCRT. (A) Schematic describing the ESCRT reporter assay. An EGFP-RNF152 fusion is expressed alongside an IRES-driven mScarlet in the HEK α-syn expressing cell line. The mScarlet serves as an internal control for reporter expression, while the EGFP level tracks degradation of the RNF152 reporter, which is internalized into the lysosomal lumen and degraded in an ESCRT-III-dependent manner. EGFP, enhanced green fluorescent protein. IRES, internal ribosome entry site. (B) Representative immunoblot of lysates from cells expressing the ESCRT reporter with strong knockdown of CHMP2A and CHMP2B. Membranes were stained with anti-GFP, anti-CHMP2A, anti-CHMP2B, anti-α-syn, and anti-β-actin. β-actin served as a loading control. Quantification appears in Figure S3D (n=3). KD, knockdown. NT, non-targeting control. (C) Flow cytometry to measure the stability (EGFP/mScarlet) of the ESCRT reporter after strong CHMP2A and CHMP2B knockdown, normalized to the value in NT. Error bars indicate mean ± SEM (n=3). ***p<0.001 relative to NT control by one-way ANOVA. (D) Flow cytometry of the ESCRT reporter cell line after exposure to PFFs, Lpf, and/or Dox to control the α-syn aggregation state. Measurements are normalized to the untreated condition. Error bars signify mean ± SEM (n=5). ****p<0.0001 relative to untreated control by two-way ANOVA. (E) Flow cytometric ESCRT reporter stability measurements of the effect of combining aggregation induction (Lpf/PFF vs Lpf) with either mild CHMP2A/B knockdown or 5 hr exposure to 100 nM of the E1 ubiquitin ligase inhibitor MLN7243 (MLN) (n=3 for the knockdown conditions and n = 5 for MLN conditions). EGFP/mScarlet values are normalized to appropriate control condition, either Lpf/NT or Lpf/DMSO. Error bars denote mean ± SEM. Arrows mark the EGFP/mScarlet ratios that would be expected if the combined perturbations were additive. Numbering for each bar (referred to in the text) appears below. The supporting immunoblot of the mild knockdown and a quantification thereof appear in Figures S3I and S3J. **p<0.01;****p<0.0001 relative to NT or DMSO control by two-way ANOVA. See also Figure S3.

To establish that the reporter is sensitive to disruption of ESCRT-III activity, we took the existence of paralogous CHMP2 proteins (CHMP2A and 2B) with overlapping functions into consideration^31,54^, both of which associate with α-syn aggregates (Figure S1F). We targeted CHMP2A using siRNA and CHMP2B using a Cas9-sgRNA pair, achieving a 90% and 78% reduction in protein levels, respectively, without affecting α-syn levels (Figure 3B and Figure S3D). While loss of each individual CHMP2 protein did not affect the reporter, CHMP2A/B double knockdown led to nearly four-fold stabilization (Figure 3C), effecting accumulation of the full-length reporter as well as high molecular weight material likely representing ubiquitinated protein (Figure 3B). Given that reporter stabilization required knockdown of both CHMP2 paralogs, we hereafter use double knockdown as an ESCRT perturbation in the HEK system. Stabilization of the reporter was also observed upon overexpressing a dominant negative CHMP2B mutant, Q165X, that causes Frontotemporal Dementia^69^, but not upon overexpression of wt CHMP2B (Figure S3E). Turning off α-syn expression with Dox had no effect on reporter levels, nor did it alter the stabilization induced by CHMP2B Q165X (Figure S3E). The reporter construct thus responds to loss of ESCRT-III protein availability.

We next analyzed whether α-syn aggregation affects ESCRT function. Inducing aggregation with Lpf/PFF treatment for two days stabilized the ESCRT reporter 1.6-fold, while Lpf or PFFs alone had no effect (Figure 3D). This stabilization disappeared when α-syn expression was switched off with Dox (Figure 3D). Inducing aggregation without the use of Lpf, by exposing cells to PFFs for 6 days, also stabilized the reporter (Figure S3F). Thus, reporter stabilization occurred only under conditions that allow for substantial production of α-syn aggregates from intracellular α-syn (as in Figures 1B and S1B). These results indicate that production of α-syn aggregates, which sequester ESCRT-III proteins, leads to impairment of ESCRT function.

Assuming that α-syn aggregation perturbs ESCRT function by sequestering ESCRT-III subunits, the reporter should be both ubiquitinated and lysosome-associated. We next analyzed the reporter’s ubiquitination status by anti-ubiquitin pulldown. Induction of aggregation led to an increase in global ubiquitinated species in the input (PFF/Lpf vs Lpf control; Figure S3G). Aggregate induction caused the accumulation of ubiquitinated reporter (Figure S3G), migrating as a smear higher in molecular weight than full-length GFP-RNF152 (Figure S3G). We note that aggregation induction also stabilized a cleaved monomeric GFP band (Figure S3G), a common stable degradation product after lysosomal digestion of fluorescent protein fusions^124^. Nonetheless, stabilization of the full-length band matched the degree of stabilization assessed by flow cytometry (Figures S3G and 3D). To assess the lysosome association of the reporter, we performed lysosome immunoprecipitation (Lyso-IP), which employs expression of an HA-tagged lysosomal membrane protein, TMEM192^125^. As expected, HA pulldown enriched the lysosomal marker LAMP1 specifically in cells expressing TMEM192-HA, without enriching markers of other subcellular compartments (Figure S3H). This pulldown also recovered a greater amount of the reporter after α-syn aggregation was induced, reflecting the increased abundance observed in the input fractions (Figure S3H). These data argue for α-syn aggregation stabilizing the ESCRT reporter in a ubiquitinated and lysosome-associated state, consistent with a loss of ESCRT-III availability.

When two perturbations converge on a common target, the combination of perturbations often produces synergistic, non-additive phenotypes^126,127^. This property can be used to discover the molecular target of a perturbation. We leveraged the quantitative nature of the ESCRT reporter to determine whether ESCRT-III knockdown and α-syn aggregation stabilize the reporter in the same way. To avoid saturating the reporter, we used a mild double knockdown of CHMP2A/B (51% for CHMP2A and 59% for CHMP2B; Figures S3I and S3J) that did not result in significant stabilization (Figure 3E; bars 1 and 3). Induction of aggregation resulted in 1.5-fold stabilization (Figure 3E; bars 1 and 2). Notably, the combination of the two perturbations synergized to cause a 3.1-fold increase in reporter stability (Figure 3E; bars 1 and 4). As a negative control, we combined α-syn aggregation with E1 ubiquitin ligase inhibition as a perturbation not mechanistically related to impaired ESCRT function. E1 inhibition did not synergize with induction of α-syn aggregation (Figure 3E; bars 5 and 8). The combination of a partial CHMP2 knockdown and α-syn aggregation therefore synergistically disrupts ESCRT activity. This synergy is consistent with both CHMP2 knockdown and α-syn aggregation limiting the availability of CHMP2 subunits (and other ESCRT-III factors), thereby reducing ESCRT function.

### Interaction with aggregated α-syn targets ESCRT-III for collateral degradation

We next considered that α-syn aggregation could deplete the available pool of ESCRT-III subunits by two mechanisms. First, interaction with aggregates could lead to spatial confinement, restricting ESCRT-III subunits from reaching their sites of action. Second, we previously observed that production of aggregates reduced CHMP2B levels by roughly half in HEK cells and primary neurons (Figures 1B, S1D, and S1E). Induction of α-syn aggregation additionally reduced CHMP2A and CHMP3 abundance in HEK cells, without significantly altering levels of CHMP4B or CHMP6 (Figures S4A and S4B). We therefore sought to better understand this mechanism of aggregation-dependent reduction of ESCRT-III protein levels.

To explore the interplay between α-syn aggregation and ESCRT-III protein depletion, we measured CHMP2B levels after PFF treatment in α-syn expressing HEK cells. We omitted Lpf usage, reasoning that the slower pace of aggregation in the absence of Lpf would allow for tighter temporal control than Lpf-facilitated aggregation. By immunoblot, CHMP2B levels declined only after appearance of α-syn aggregates (high molecular weight SDS-resistant smears), then continued decreasing until the experiment ended at 6 days post-PFF treatment (Figure 4A). Additionally, treating with Dox to limit α-syn expression after PFF exposure diminished the amount of α-syn aggregates and ameliorated the decline in CHMP2B levels (Figure 4B). These results establish α-syn aggregate production as a precursor to the reduction in ESCRT-III subunit abundance.

**Figure 4.**
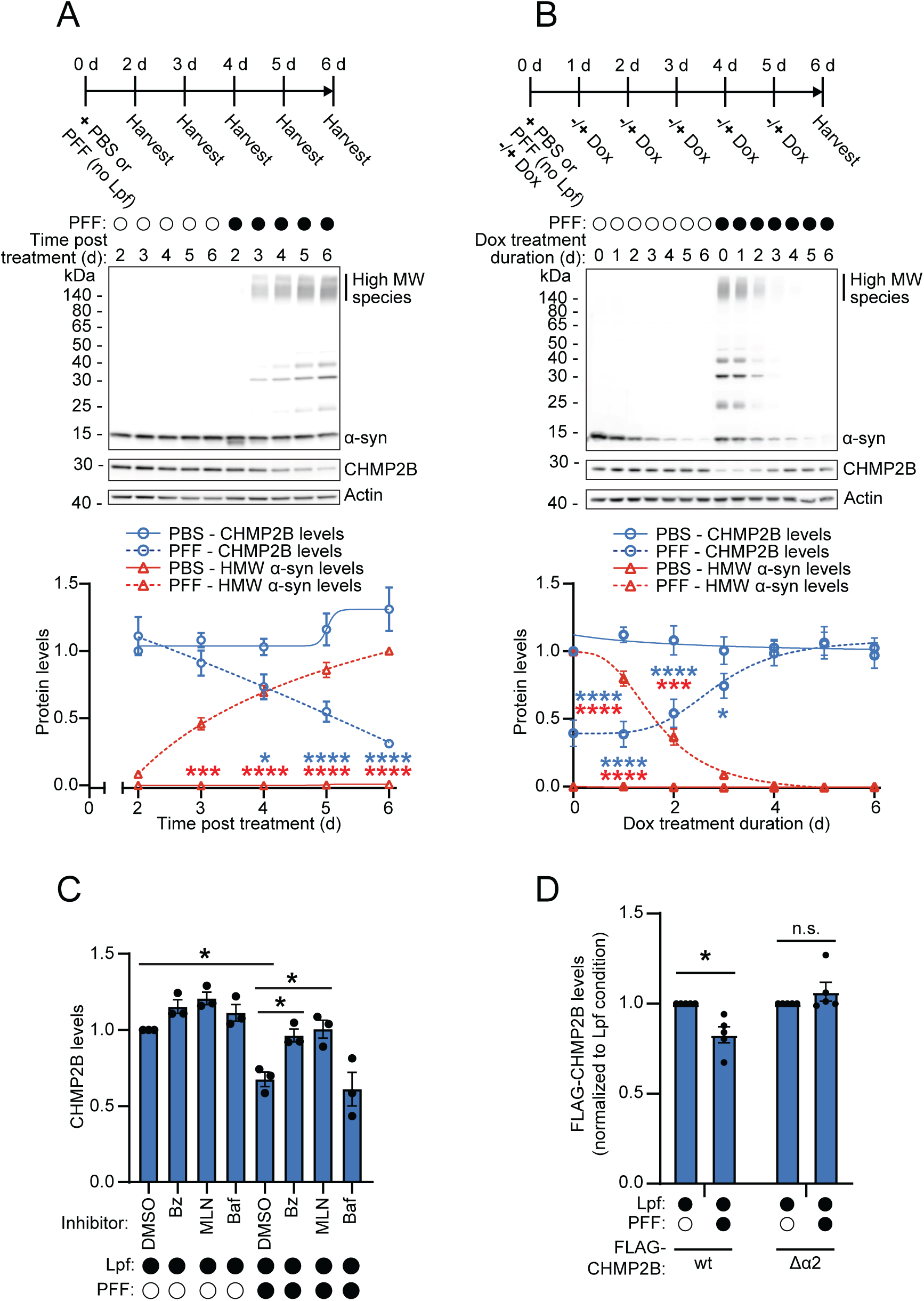
α-Syn aggregation triggers collateral degradation of ESCRT-III proteins. (A and B) Representative immunoblots of HEK α-syn expressing cell lysates after treatment with either PBS or PFFs (without Lpf) for increasing amounts of time (A) or for 6 days with Dox treatment at different times (B), according to the timeline above. Note in (B) that 6 days of Dox treatment indicates Dox addition on day 0, with 5 days of treatment indicating addition on day 1, etc. Immunoblots were stained with anti-α-syn, anti-CHMP2B, and anti-β-actin antibodies. β-Actin was the loading control. Densiometric quantifications of the immunoblots appear below. CHMP2B values are normalized to the value obtained at the first timepoint and α-syn values are normalized to the maximum value – 6 d in (A) and 0 d in (B). Error bars signify mean ± SEM (n=3). *p<0.05;***p<0.001;****p<0.0001 by two-way ANOVA, pairwise comparisons shown between PBS/PFF-treated CHMP2B (blue) or high MW α-syn (red) measurements at matched timepoints. (C) Densiometric quantification of immunoblots in Figure S4C, showing the effect of 8 hr inhibitor treatment on CHMP2B levels with and without aggregation induction. Bz, 500 nM bortezomib (proteasome inhibitor). MLN, 500 nM MLN7243 (E1 ubiquitin ligase inhibitor). Baf, 250 nM bafilomycin A1 (lysosomal V-ATPase inhibitor). Data are presented after β-actin normalization and subsequent normalization to the Lpf/DMSO control condition. Error bars indicate mean ± SEM (n=3). *p<0.05 by two-way ANOVA. Only significant comparisons are shown. (D) Densiometric quantification of immunoblots in Figure S4E, testing how aggregation induction affects transfected FLAG-tagged CHMP2B variant levels. CHMP2B levels were normalized to β-actin, then to the value obtained for each variant in the Lpf control condition. Error bars represent mean ± SEM (n=5). n.s. p>0.05;*p<0.05 by two-way ANOVA. See also Figure S4.

Considering that α-syn inclusions contain strong ubiquitin and CHMP2B signal (Figures 1C and S1A), we hypothesized that aggregation leads to ubiquitin-dependent CHMP2B degradation, perhaps resulting from an α-syn-targeting E3 ligase nonspecifically ubiquitinating α-syn-bound CHMP2B. We tested this hypothesis by inducing aggregation for two days and then treating cells with inhibitors of various degradative pathways. Inhibition of E1 ubiquitin ligases or the proteasome rescued CHMP2B levels, while inhibition of lysosome acidification with bafilomycin had no effect (Figures 4C and S4C). Furthermore, anti-ubiquitin immunoprecipitation recovered increased levels of ubiquitinated CHMP2B after aggregation induction (Figure S4D). This immunoprecipitation additionally co-purified unmodified CHMP2B, suggesting that non-ubiquitinated CHMP2B may be complexed with ubiquitinated cellular material (Figure S4D). Thus, α-syn aggregation leads to ubiquitination and proteasomal degradation of CHMP2B.

To test directly whether ESCRT-III subunit degradation relies on interaction with α-syn aggregates, we induced aggregation after overexpressing either FLAG-tagged wt CHMP2B or the α2 deletion mutant with diminished ability to associate with α-syn aggregates (Figure S2C). While induction of aggregation reduced wt FLAG-CHMP2B levels, the α2 mutant was unaffected (Figures 4D and S4E), indicating that aggregate binding is required for aggregation-induced CHMP2B degradation. Interestingly, overexpression of wt FLAG-CHMP2B attenuated endogenous CHMP2B degradation (Figures S4E and S4F). In contrast, the non-degraded α2 mutant did not impact endogenous CHMP2B degradation, suggesting that the collateral degradation pathway is saturable (Figures S4E and S4F). Saturation of collateral degradation would also explain why cells degrade overexpressed FLAG-CHMP2B less readily than endogenous CHMP2B (17% vs 38%, Figures 4D, S4E, and S4F).

These data suggest that interaction with α-syn aggregates triggers degradation of CHMP2B and, by extension, other ESCRT-III subunits. We hereafter refer to this phenomenon as “collateral degradation,” as degradation appears to be a collateral consequence of aggregate binding. This mechanism contributes to the functional depletion of ESCRT-III factors, in addition to their sequestration.

### α-Syn aggregation impairs endolysosomal homeostasis

The findings described above suggested that α-syn aggregation, by compromising ESCRT-III function, may impair endolysosome homeostasis and lead to loss of endolysosome membrane integrity. Perforation of these membranes exposes β-galactoside sugars on the inner leaflet to the contents of the cytoplasm, which results in microscopically observable clustering of Galectin proteins on the membrane^128–131 132^. To assess endolysosome membrane integrity, we expressed a FRET pair of Gal3 (tagged with mRuby3 and mClover3) in the α-syn expressing HEK cells (Figure 5A). As expected, exposing these cells to the lysosome-damaging drug L-Leucyl-L-Leucine methyl ester (LLOMe) elicited a dose-dependent increase in the number of Gal3 FRET-positive cells (Figures S5A and S5B). Furthermore, LLOMe treatment triggered the formation of mRuby3- and mClover3-positive Gal3 puncta that co-localized with markers of the endolysosomal system (Figure S5C).

**Figure 5.**
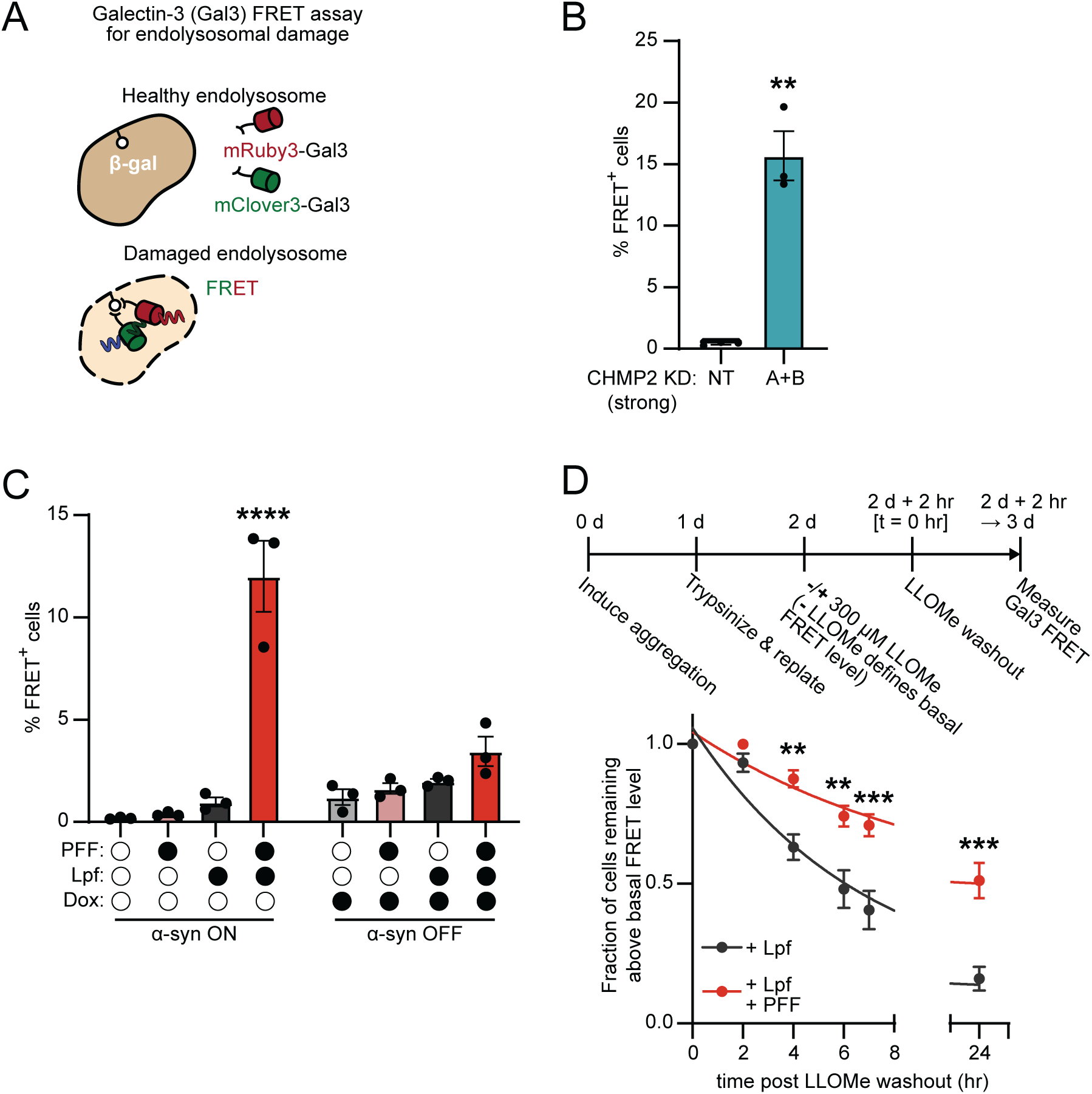
α-Syn aggregation causes endolysosomal damage. (A) Schematic describing the Gal3 FRET assay to measure endolysosomal damage. A FRET pair of mRuby3- and mClover3-tagged Gal3 is expressed in HEK α-syn expressing cells. When endolysosomal damage occurs, clusters of the normally cytoplasmic Gal3 fusions bind newly-exposed β-galactoside sugars (β-gal) on the inner leaflet of the endolysosomal membrane, producing FRET. (B) Gal3 FRET flow cytometry of cells after strong CHMP2A/B knockdown (as in Figures 3B and 3C). Error bars signify mean ± SEM (n=3). **p<0.01 relative to NT control by t-test. (C) Flow cytometry of Gal3 FRET cells after treatment with indicated α-syn aggregation-inducing conditions. Error bars signify mean ± SEM (n=3). ****p<0.0001 relative to untreated control by two-way ANOVA. (D) Flow cytometry timecourse of the disappearance of Gal3 FRET signal after cells with and without aggregates (Lpf/PFF and Lpf) were briefly exposed to 300 µM LLOMe. Within the two conditions, the fraction of FRET-positive cells was normalized to the value obtained at t=0. Error bars signify mean ± SEM (n=4, except for t=24 hr where n=3). Statistical comparisons were made within each timepoint. **p<0.01;***p<0.001 by two-way ANOVA. See also Figure S5.

Consistent with the role of ESCRT-III proteins in preserving endolysosomal membrane integrity^40,42^, strong CHMP2A/B double knockdown increased the amount of Gal3 FRET-positive cells even in the absence of additional lysosome stress (Figure 5B). Furthermore, overexpression of dominant negative CHMP2B Q165X also increased the proportion of FRET-positive cells, while wt CHMP2B had no effect (Figure S5D). These data support the notion that loss of ESCRT-III availability raises the steady-state level of endolysosomal damage, indicating a necessity for constant repair. This finding is consistent with a recent observation of perforated endolysosomes in unstressed neurons^35^.

To test our hypothesis that α-syn aggregation increases endolysosomal damage, we treated the Gal3 reporter cells with Lpf and PFFs for two days. Aggregate formation markedly increased the fraction of Gal3 FRET-positive cells, while treatment with Lpf or PFFs alone had no effect (Figure 5C). Halting intracellular α-syn expression with Dox prevented the increase in FRET-positive cells (Figure 5C). Consistently, cells with aggregates contained more prominent Gal3 puncta than cells without aggregates; these puncta appeared both within the bounds of the aggregates as well as in locations distal to the aggregates (Figure S5E). Endolysosome membrane damage, reflected by Gal3 FRET, was also observed when α-syn aggregation was induced by treatment with PFFs for 6 days in the absence of Lpf (Figure S5F). Thus, α-syn aggregate formation triggers endolysosomal damage, as we observed with direct perturbation of ESCRT-III proteins (Figure 5B).

α-Syn aggregation could either increase steady-state endolysosomal damage by acting as the damaging agent, by inhibiting repair, or by a combination of both. To measure endolysosomal membrane repair, we briefly exposed cells carrying α-syn aggregates (Lpf/PFF treatment) and control cells (Lpf only) to LLOMe, then washed the drug out and followed the reduction of Gal3 FRET signal over time. Although the amount of Gal3 FRET-positive cells decreased after LLOMe washout in both cases, this decrease was substantially slower in aggregate-induced cells (Figure 5D), indicating diminished repair of endolysosomal damage upon α-syn aggregation.

In summary, these results support a model wherein endolysosomes exist in a balance between damage and repair, with repair at least partially mediated by ESCRT-III. α-Syn aggregates sequester ESCRT-III subunits and target them for degradation, thereby inhibiting their function and impairing the ability of cells to repair damaged endolysosomes.

### Disruption of ESCRT function exacerbates seeded α-syn aggregation

α-Syn fibrils can enter cells by the traditional endocytic route, where they are internalized after binding cell surface receptors, arriving in endosomes that mature into late endosomes and eventually into lysosomes^25,28–30,133^. Accordingly, in the cellular model, PFFs labeled with the fluorescent dye Alexa 647 co-localized with the early endosome marker EEA1 after 15 min of incubation and with the late endosome/lysosome marker LAMP2 after 4 hr (Figure S6A). We wondered whether the endolysosomal damage triggered by ESCRT-III disruption could liberate such fibrils from the endolysosomal compartment and allow them to seed intracellular α-syn aggregation.

To measure seeded α-syn aggregation, we constructed a HEK cell line expressing two A53T mutant α-syn variants, tagged C-terminally with either FRET donor mClover3 or FRET acceptor mRuby3 (Figure 6A). Inducing aggregation with PFFs and Lpf produced 68% FRET-positive cells within 2 days (Figure S6B), consistent with our results inducing aggregation in cells expressing untagged α-syn (Figure 1B). This increase in FRET coincided with the appearance of filter-retained aggregate material that reacted with anti-GFP (detecting mClover) and anti-pS129 α-syn antibodies (Figure S6C). Induction of aggregation also caused formation of mClover3- and mRuby3-positive inclusions that stained for classical markers of α-syn aggregation (Figure S6D). Thus, the α-syn FRET cell line reports on the formation of *bona fide* α-syn aggregates.

**Figure 6.**
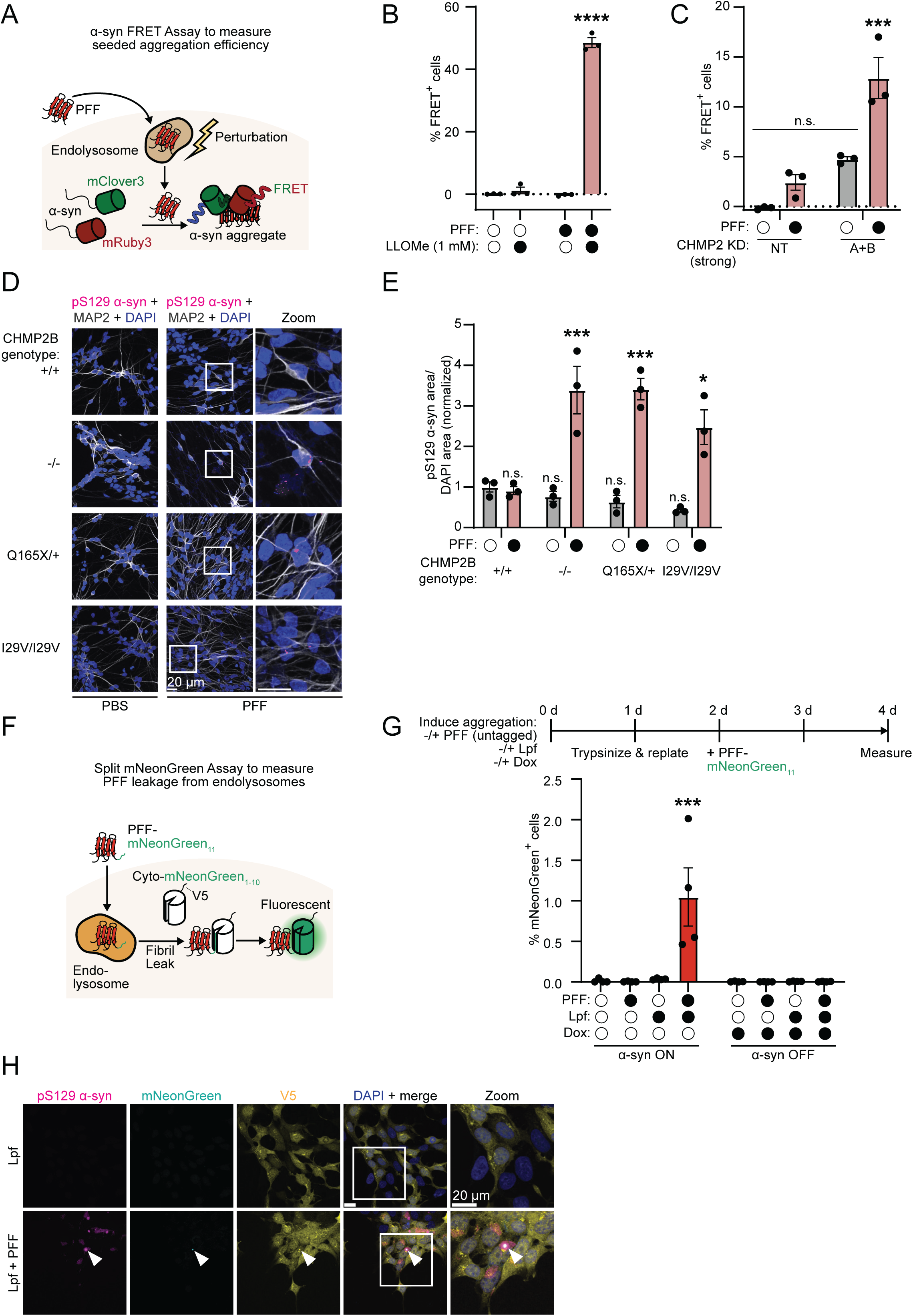
ESCRT-III disruption and α-syn aggregation cause fibril leakage from endolysosomes. (A) Schematic of FRET assay to measure seeded α-syn aggregation. HEK cells expressing a FRET pair of α-syn-mClover3 and α-syn-mRuby3 are treated with PFFs. Additional perturbations to the endolysosomal system are combined with PFF treatment to affect PFF leakage into the cytoplasm. When α-syn aggregates, mClover3 and mRuby3 produce a FRET signal that can be detected by flow cytometry. (B) Flow cytometry of α-syn FRET cells exposed to PFFs, then exposed 1 hr later to the lysosomotropic drug LLOMe. Error bars signify mean ± SEM (n=3). ****p<0.0001 relative to untreated control by two-way ANOVA. (C) Flow cytometry of the effect of strong CHMP2A/B knockdown (as in Figures 3B and 3C) on seeded α-syn aggregation, as assessed by α-syn FRET. Error bars represent mean ± SEM (n=3). n.s. p>0.05;***p<0.001 relative to untreated NT control by two-way ANOVA. (D) Representative immunofluorescence micrographs of seeded α-syn aggregation in iNeurons with indicated CHMP2B mutations. iNeurons were treated with PFFs for 7 days, starting on DIV16. Samples were stained with anti-pS129 α-syn and anti-MAP2 antibodies (Scale bar, 20 μm). Quantification of α-syn aggregation from micrographs of PFF-seeded iNeurons with indicated CHMP2B mutations. Error bars signify mean ± SEM (n=3). ***p<0.001;*p<0.05; n.s. p>0.05 relative to untreated CHMP2B^+/+^ control by two-way ANOVA. (E) Schematic describing the split mNeonGreen PFF leakage assay. HEK α-syn cells expressing a cytosolically-targeted split mNeonGreen (cyto-mNeonGreen_1-10_) are exposed to PFFs tagged with the complementary mNeonGreen fragment (PFF-mNeonGreen_11_. Leakage of endocytosed tagged fibrils into the cytoplasm would result in mNeonGreen complementation and fluorescence. Cyto-mNeonGreen1-10 is also V5-tagged to enable its visualization independently from fluorescence complementation. (F) Flow cytometry of the split mNeonGreen PFF leakage assay in cells pre-treated to induce α-syn aggregation (as in Figure 1B) prior to exposure to PFF-mNeonGreen_11_. Error bars represent mean ± SEM (n=4). ***p<0.001 relative to no pre-treatment control by two-way ANOVA. (G) Representative immunofluorescence microscopy demonstrating the localization of the complemented mNeonGreen signal in selected conditions from the experiment in (G). Cells were stained with anti-pS129 α-syn to mark α-syn inclusions (denoted by arrowheads) and anti-V5 to mark the total population of Cyto-mNeonGreen_1-10_ (Scale bar, 20 μm). See also Figures S6 and S7.

We then employed the α-syn FRET cells to test whether endolysosomal damage facilitates seeded aggregation. In these assays we omitted the use of Lpf to ensure that PFFs enter cells through the physiologic endocytic route (Figure 6A). We first tested whether direct lysosome damage with LLOMe would facilitate seeding. While PFFs alone were without measurable effect within 2 days of treatment, the combination of PFFs and LLOMe resulted in a massive increase in FRET-positive cells, reflecting intracellular α-syn aggregation (Figure 6B). Perforations in the endolysosomal system therefore allow for PFFs to efficiently seed intracellular α-syn aggregation. We suggest that without induced perforation, seeding is inefficient because only small amounts of seeds reach the cytoplasm. Inefficient cytoplasmic translocation would explain why seeding takes longer without Lpf, with initial aggregates appearing after 3 days of PFF treatment (Figure 4A).

Next, we investigated whether ESCRT-III perturbation facilitates α-syn seeding similarly to LLOMe. CHMP2A/B double knockdown significantly increased the fraction of FRET-positive cells after PFF treatment without Lpf (Figure 6C). This knockdown also generated a small FRET-positive cell population in the absence of PFFs, although this effect was not significant (Figure 6C). Furthermore, expression of dominant negative CHMP2B Q165X also increased the proportion of FRET-positive cells upon PFF treatment, while wt CHMP2B had no effect (Figure S6E). Thus, perturbation of ESCRT-III, which we showed to cause endolysosomal damage (Figure 5B), also exacerbates templated α-syn aggregation.

Notably, CHMP2/B double knockdown had no effect on PFF-induced aggregation when Lpf was present (Figure S6F), suggesting that seed transit from endolysosomes to the cytosol is not limiting for seeding in this condition. This observation validates Lpf as a tool to study the effect of intracellular α-syn aggregation on the ESCRT system without confounding effects on cytoplasmic entry.

To determine whether loss of ESCRT-III also enhances α-syn seeding in neuronal cells, we disrupted ESCRT-III in iPSCs before differentiating these cells into iNeurons by NGN2-driven reprogramming^134^. We performed the experiment at endogenous α-syn levels to keep aggregation signal low and avoid a saturating aggregation scheme that would mask the effect of ESCRT-III disruption. Preliminary experiments suggested that deleting CHMP2B alone was sufficient to produce a clear phenotype in iNeurons, in contrast to HEK cells. We therefore perturbed ESCRT-III by deleting CHMP2B in the parental line with CRISPR-Cas9 (or Cpf1) as well as editing two dominant disease-associated point-mutations into CHMP2B – Q165X and I29V (Figure S6G), yielding CHMP2B^-/-^, CHMP2B^Q165X/+^, and CHMP2B^I29V/I29V^ cell lines. iNeurons were seeding-resistant, as PFF treatment in the parental line (without Lpf) did not reliably permit formation of pS129 α-syn inclusions (Figures 6D and 6E). However, all CHMP2B mutations significantly increased the number of iNeurons with inclusions upon PFF treatment (Figures 6D and 6E), indicating that loss of CHMP2B alone facilitates α-syn seeding in neurons.

In summary, PFFs can escape from endolysosomes and template misfolding and aggregation of soluble α-syn. ESCRT-III proteins, which help maintain endolysosomal homeostasis, normally limit seeding. As a result, loss of ESCRT-III function increases α-syn seeding efficiency.

### α-Syn aggregation enhances leakage of exogenous fibrils into the cytoplasm

Our results so far suggested that α-syn aggregates sequester ESCRT-III members and trigger their degradation, inhibiting their ability to maintain endolysosomal integrity. We hypothesized that the collapse of endolysosome homeostasis triggered by α-syn aggregation would lead to increased leakage of endocytosed fibrils into the cytoplasm. These secondary fibril leakage events could accelerate conversion of monomeric α-syn into aggregates by providing more sites for templated aggregation. As a critical prediction of this mechanism, cells with aggregates should be more prone to leaking endocytosed fibrils into the cytoplasm compared to cells without aggregates.

To test this hypothesis, we developed a flow cytometry-based fluorescence complementation assay using a split mNeonGreen system^135^. We generated exogenous PFFs from α-syn tagged C-terminally with the final beta strand of mNeonGreen (PFF-mNeonGreen_11_) and stably expressed beta strands 1-10 in the cytoplasm (cyto-mNeonGreen_1-10_) of the α-syn expressing acceptor cells (see STAR methods; Figure 6F). Introducing the tagged PFFs into the cytoplasm of these cells using Lpf resulted in mNeonGreen fluorescence and mNeonGreen-positive inclusions (Figures S7A and S7B). The fluorescence signal, though robust, was present in fewer cells (18%) than what we previously obtained with the α-syn FRET assay (68%; Figure S6B), potentially due to limited fluorescence complementation in the topological context of α-syn fibrils. The mNeonGreen inclusions were mostly pS129-positive (Figure S7C). However, pS129-positive, mNeonGreen-negative aggregates were also observed (Figure S7C), suggesting seeding events without fluorescence complementation as well as secondary α-syn aggregates resulting from either primary aggregate fragmentation or secondary nucleation^136–139^. Efficient fluorescence complementation also occurred upon PFF-mNeonGreen_11_ treatment in the absence of Lpf when the function of ESCRT-III in maintaining membrane integrity was disrupted by overexpression of the dominant negative CHMP2B Q165X (Figure S7D). Thus, the split mNeonGreen assay detects PFF leakage from endolysosomes after loss of ESCRT-III function.

We next tested the prediction that α-syn aggregation facilitates leakage of exogenous PFFs into the cytoplasm. Cells were first pre-treated using aggregation inducing conditions (untagged PFFs plus Lpf; Figure 1B) to generate α-syn aggregates that were not detectable by the split mNeonGreen assay (Figures 6G, S7A and S7B). After 2 days we exposed the cells to PFF-mNeonGreen_11_ without Lpf. Significantly, PFF-mNeonGreen_11_ exposure caused a fluorescence complementation signal only when the cells contained preexistent intracellular α-syn aggregates induced by PFF/Lpf pretreatment (Figure 6G). Dox treatment, which depletes the intracellular supply of α-syn and thereby prevents aggregation, abolished this increase in mNeonGreen fluorescence (Figure 6G). The mNeonGreen puncta that formed in cells already carrying α-syn aggregates co-localized with pS129 α-syn inclusions (Figure 6H). These data therefore provide evidence that the endolysosome damage caused by α-syn aggregation exacerbates leakage of exogenously-added α-syn fibrils from endolysosomes into the cytoplasm, thereby increasing aggregate load. As expected, the magnitude of leakage, indicated by the fraction of mNeonGreen-positive cells (Figure 6G), is much lower than upon direct disruption of ESCRT-III function by overexpression of dominant negative CHMP2B Q165X (Figure S7D).

## DISCUSSION

In this study, we uncovered the cell biological implications of the clinically-observed co-localization between the ESCRT-III factor CHMP2B and disease protein inclusions, specifically those of α-syn^85–89,91^. We found that CHMP2B binds preferentially to fibrillar α-syn aggregates rather than to monomeric α-syn (Figure 2B), with helix α2 of CHMP2B being necessary and sufficient to mediate this conformation-specific interaction (Figures S2C, 2C, and 2D). As a consequence, α-syn aggregates sequester CHMP2B as well as other ESCRT-III proteins including CHMP2A, CHMP3, and CHMP4B that feature similar helical regions (Figures 1C, 1D, S1F, and 7 Panels 1-3). In addition to sequestration of these proteins, aberrant interaction with α-syn aggregates triggers collateral degradation of some ESCRT-III subunits by the proteasome (Figures 4A-D, S4A-F, and 7 Panel 3). These findings support a model wherein α-syn aggregation depletes the available pool of ESCRT-III subunits, inhibiting ESCRT function and collapsing endolysosomal homeostasis (Figures 7 Panels 3 and 4). As a result, cells carrying inclusions accumulate endolysosome membrane damage and become prone to leaking endocytosed fibrils into the cytoplasm (Figure 7 Panel 4). We posit that this leakage triggers further seeding events, increasing endogenous aggregate burden and completing a feedback loop (Figure 7 Panel 4), wherein α-syn aggregation disrupts ESCRT function, begetting further aggregation and endolysosomal damage.

**Figure 7.**
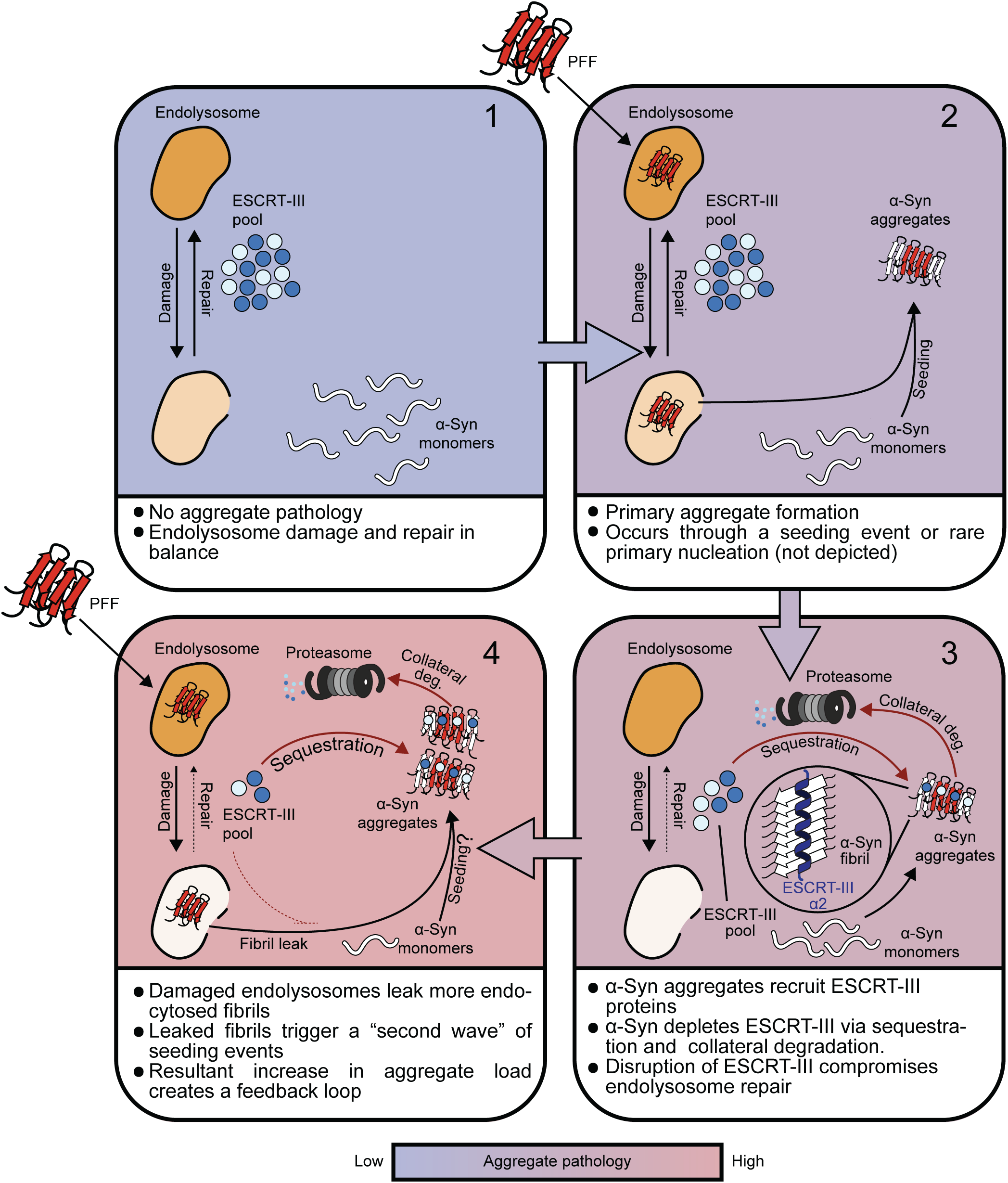
Working model of a feedback loop connecting seeded α-syn aggregation and loss of ESCRT function. (1) In healthy cells, α-syn exists in a monomeric, soluble state and the ESCRT-III pool suffices to repair endolysosomal damage. (2) α-Syn aggregate pathology begins as the primary aggregate forms either through seeding or primary nucleation. (3) α-Syn aggregates sequester ESCRT-III subunits via interaction with helix α2. Interaction with aggregates additionally targets ESCRT-III proteins for collateral degradation by the proteasome. These dual effects functionally deplete ESCRT-III, impairing endolysosomal integrity. Below, scale bar indicating the degree of aggregate pathology in panels. (4) Endolysosomes disrupted by ESCRT-III depletion leak fibrils, triggering a hypothesized “second wave” of seeding events. This second wave would increase the kinetics of aggregation and further perturb ESCRT-III, forming a feedback loop. Note that question marks denote facets of the model that remain open questions, i.e. the “second wave” of seeding.

### Experimental systems for cell biological analysis of α-syn aggregation

Key to our investigation was a cellular system that controlled α-syn aggregation (Figure 1A). We achieved this with inducible α-syn expression, PFFs to seed intracellular aggregation, and Lpf to introduce PFFs into the cytoplasm. This control allowed precise analysis of how α-syn aggregation impacts specific phenotypes. Notably, ESCRT-III-sequestering inclusions of intracellular α-syn had the strongest effect, while α-syn expression or brief PFF treatment alone caused minimal disruption (Figures 1B, S1D, 3D, S3F, 5C, S5F, and 6G). Silencing α-syn expression significantly reduced toxicity, highlighting its therapeutic potential^140^.

Because feedback loops amplify inputs, isolating each segment experimentally requires severing the loop. Lpf permitted α-syn aggregate seeding at saturation, unaffected by further ESCRT-III perturbation (Figure S6F), which was crucial for untangling the feedback mechanism. This approach enabled us to measure the combined effect of aggregation and CHMP2 knockdown on ESCRT function (Figure 3E), without impaired ESCRT function leading to increased aggregation and amplifying the effect. We nonetheless controlled for Lpf treatment throughout and reproduced key results under Lpf-independent aggregation schemes.

### Sequestration and collateral degradation functionally deplete ESCRT-III

Association of ESCRT-III proteins with α-syn aggregates can lead to functional depletion in two ways. First, spatial confinement within aggregates may prevent ESCRT-III subunits from reaching their site of action. Second, sequestration could cause incidental degradation of ESCRT-III proteins along with aggregated α-syn, which we call “collateral degradation” (Figure 7 Panel 3).

Analysis of patient brains offers a snapshot of pathological aggregates, wherein ESCRT-III proteins co-localize with α-syn^86,87,89^. How this interaction first originates remains an open question. Recent evidence indicates that tau fibrils upon uptake cause nanoscale tears in endolysosomes, allowing them to protrude into the cytoplasm and template the misfolding of monomeric tau on the other side of the membrane. These sites of membrane damage also recruit the ESCRT-III repair machinery^34^. If α-syn aggregates originate similarly, the first instances of α-syn-ESCRT-III interaction may occur at these hotspots of fibril growth and membrane damage. However, we found that CHMP2B does not need to bind to membranes to interact with α-syn aggregates (Figures 2B and S2B). The cytoplasmic pool of ESCRT-III proteins may therefore also bind to aggregates distal from endolysosomes, which originate either from fragmentation of the primary aggregate or through secondary nucleation of new α-syn on the surface of fibrils^136–139^.

Our results indicate that ESCRT-III proteins bind specifically to the aggregated state of α-syn (Figures 1C, 1D, S1C, S1F, 2A, 2B, 2D, and S2A). Contrary to a recent report^90^, we detected no interaction between CHMP2B with α-syn monomers or α-syn from lysates lacking aggregates (Figures 2A, 2B, and S2A). We mapped the biochemical basis of this aggregation-specific interaction to CHMP2B helix 2 (α2), spanning residues 55 to 96 (Figures S2C, 2C, and 2D). In solved structures of ESCRT-III oligomers, α2 forms an extended helix with α3. This extended helical region forms an elongated hydrophobic interaction interface between monomers^47,51,53,141^. We speculate that this same region recognizes α-syn fibrils, potentially using its hydrophobic face to bind a hydrophobic groove on the fibril surface (Figures S2D, S2G, and 7 Panel 3). The apparent broad specificity of α2, and by extension CHMP2B^85–89,91^, for multiple amyloid-like proteins (Figure 2D) suggests that the motif recognized on the fibril surface might be a general feature of amyloid folds. This broad recognition capability and its compact size may make α2 an attractive vehicle for delivering therapeutic molecules to amyloid fibrils across diseases.

All ESCRT-III complex structures feature this extended α2/3 helical binding motif^47,51,53,141^, potentially explaining why α-syn sequesters several different ESCRT-III proteins in addition to CHMP2B. Indeed, while this study focused on CHMP2B and its paralog CHMP2A, we also detected recruitment of the core ESCRT-III subunits CHMP3 and CHMP4B to α-syn aggregates (Figure S1F). The effect of α-syn aggregation on ESCRT function may therefore result from a broad effect on multiple ESCRT-III proteins, not just CHMP2B.

As seen in Lewy Body Dementia patients^86^, we observed that α-syn aggregation reduced CHMP2B levels and several other ESCRT-III subunits (Figures 1B and S4A). This decrease depended on α-syn inclusion production and was not stimulated by α-syn expression or by PFF exposure without an intracellular α-syn pool for templating into aggregates (Figure 1B). Inhibition of ubiquitination or the proteasome suppressed the decrease, as did mutation of the α-syn binding site on CHMP2B (Figures 4C, 4D, S4C, S4E, and S4F). Furthermore, excess collateral degradation substrates saturated the pathway (Figures S4E and S4F), either by competing for endogenous CHMP2B for α-syn binding or for recognition by E3 ligases. Thus, the loss of CHMP2B levels (and by extension other ESCRT-III proteins) results from ubiquitination after co-aggregation with α-syn.

Collateral degradation is then perhaps the result of a ubiquitin-dependent clearance mechanism targeting α-syn aggregates. We posit that E3 ligases recognize aggregates and inadvertently ubiquitinate CHMP2B bound to fibrillar α-syn, alongside α-syn itself. As aggregates constantly form and become ubiquitinated, they therefore serve as a sink for ESCRT-III proteins, leading to functional ESCRT-III depletion (Figure 7 Panel 4). Inadvertent ubiquitination has been described in the context of E3 ligases ubiquitinating collaborating E1 and E2 ligases^142^, but to our knowledge has not previously been identified on sequestered proteins in aggregates. Future studies are needed to decipher mechanistic details, including the responsible E3 ligase and the α-syn species where ubiquitination takes place (e.g. small aggregate nuclei or mature fibrils). Comparative studies will determine whether collateral degradation is a common feature of disease-associated protein aggregation, perhaps representing an undesired side effect of aggregate clearance pathways.

For optimal activity, ESCRT-III subunits must assemble in a choreographed sequence^36–39,41,43, 46^. Disturbing this coordination with imbalanced subunit ratios impairs ESCRT-III function^116^. Even partial reduction of the available pool of several ESCRT-III species could thereby exert an outsized impact on ESCRT-III oligomerization. If ESCRT function were disrupted by imbalance between the many different ESCRT-III subunits, we would not anticipate that raising the levels of any individual subunit could rescue function. Attempts at ameliorating the effect of α-syn aggregation on ESCRT function might be better invested in developing strategies to raise the levels of all ESCRT-III subunits in concert or by fortifying endolysosome homeostasis through ESCRT-independent means.

### ESCRT disruption as a driver of endolysosomal dysfunction in PD

Mutations in several genes associated with endolysosomal function are linked to PD. Although the gene products perform diverse functions, they all converge on the endolysosome^143–146^. This convergence highlights a central role for endolysosomal health in PD and suggests that wide-ranging endolysosomal stressors can promote disease pathogenesis. Clinical evidence of ESCRT-III sequestration^86,87,89^ and our mechanistic work here position ESCRT disruption as a driver of α-syn-related endolysosomal dysfunction. Furthermore, feedback between endolysosomes, ESCRT, and α-syn aggregation (Figure 7) may amplify other sources of endolysosomal stress, augmenting their contribution to PD.

An emergent property of the feedback loop described here is that cells carrying α-syn inclusions tend to leak endocytosed fibrils into the cytoplasm (Figure 7 Panel 4). It is easy to imagine how ESCRT dysfunction resulting from α-syn aggregation could lead to toxicity. Yet the consequences of increased endolysosome leakiness are less apparent. Leakiness may usher in a “second wave” of α-syn seeding events and thereby accelerate the pace of aggregation and endolysosomal proteostasis decline (Figure 7 Panel 4), eventually overwhelming aggregate clearance and causing proteostasis collapse. Moreover, damaged endolysosomes could spill other endocytosed proteins into the cytoplasm that are not intended for cytoplasmic translocation, such as other fibrillar species like tau. PD and other synucleinopathies commonly feature tau co-pathology^147–150^. In addition to accelerating α-syn spreading, leakage of tau fibrils into the cytoplasm of α-syn inclusion-bearing cells could accelerate the propagation of tau aggregates. Understanding the pathophysiological relevance of enhanced endolysosomal leakiness caused by α-syn aggregation will require further investigation in animal models.

### Limitations of the study

Although we detected secondary entry of fibrils into the cytoplasm after prior α-syn aggregation (Figure 6G), we did not possess the tools to specifically ablate this arm of the feedback loop. As a result, we are unable to confirm whether this leakage of fibrils produces secondary seeding events to exacerbate aggregate burden. We anticipate that development of tools to specifically block secondary fibril leakage will clarify the functional contribution of each feedback loop component (Figure 7) and therefore identify the most promising therapeutic target.

We isolated the α-syn binding region of CHMP2B (Figures S2C, 2C, and 2D), yet the fibril-specific binding mechanism remains unknown. Answering this question will require structural insights into full-length CHMP2B or α2 bound to fibrils. Previous structural studies have revealed non-proteinaceous densities near amyloid fibrils^151,152^. However, to our knowledge only one cryogenic electron tomography reconstruction has captured a protein bound to amyloid fibrils, but the resolution was insufficient to reveal the structural basis for the interaction^153^. The dearth of such structures may stem from the need for an ideal amyloid binder with a uniform binding pattern or for updated helical reconstruction methods that can more sensitively detect fibril-adjacent densities. Given its pathomechanistic relevance, the CHMP2B-α-syn complex offers a compelling test case for such a method.

## Supporting information

Key Resources Table

Suppplementary Figures and Tables

## RESOURCE AVAILABILITY

### Lead contact

Further information and requests for resources and reagents should be directed to and will be fulfilled by the lead contact, F. Ulrich Hartl.

### Materials Availability

Plasmids generated in this study will be available on Addgene at the time of publication. Cell lines from this study are available upon request to the lead contact.

### Data and code availability

Code used in this study has been deposited at Github. Raw and processed data from this study, as well as an expanded Key Resource Table, will be available at Zenodo at the time of publication. The locations for all of these resources can be found in the Key Resource Table.

## ACKNOWLEDGMENTS

We thank Martin Spitaler, Giovanni Cardone, and Markus Oster at MPIB for assistance with flow cytometry and confocal microscopy; Nadine Wischnewski, Silvia Gaertner, and Romy Lange for protein purification efforts; the Core for Imaging Technology & Education (CITE) at Harvard Medical School for imaging assistance; Stephan Uebel and Stefan Pettera at MPIB for synthesis of the biotinylated CHMP2B_α2_ peptide; Martin Mueller for helpful comments on the manuscript. This work was supported by the Deutsche Forschungsgemeinschaft (DFG, German Research Foundation) under Germany’s Excellence Strategy within the framework of the Munich Cluster for Systems Neurology (EXC 2145 SyNergy—ID 390857198) (to F.U.H.) and by the joint efforts of The Michael J. Fox Foundation for Parkinson’s Research (MJFF) and the Aligning Science Across Parkinson’s (ASAP) initiative (to F.U.H and J.W.H.). MJFF administers the grants ASAP-000282 and ASAP-024268 on behalf of ASAP and itself. C.S.S. was supported by an EMBO Long Term Fellowship (ALTF 360-2020).

## AUTHOR CONTRIBUTIONS

C.S.S. planned and performed experiments and analyzed the data along with V.A.T., A.G., M.G., P.Y.C., and U.E.D. C.S.S., V.A.T., and F.U.H. conceived the project. D.F. and J.Z. generated critical cell lines – primary mouse neurons and CHMP2B-edited iPSCs, respectively – with supervision from J.W.H. and I.D. C.S.S. and F.U.H. wrote the initial draft of the manuscript, which was critically edited by V.A.T., A.G., and J.W.H. F.U.H. supervised the project.

## DECLARATION OF GENERATIVE AI AND AI-ASSISTED TECHNOLOGIES IN THE WRITING PROCESS

During the preparation of this work the authors used ChatGPT for help in condensing writing and generating protocols for protocols.io using the text from the STAR methods section. After using this tool, the authors reviewed and edited the content as needed and take full responsibility for the content of the publication.

## SUPPLEMENTAL INFORMATION

Supplemental information includes Figures S1-S7 and Table S1.

## STAR★METHODS

### EXPERIMENTAL MODEL AND SUBJECT DETAILS

#### Cell culture

HEK293T cells were cultured in DMEM (Thermo Fisher Scientific, 11995073) containing 10% FBS (Thermo Fisher Scientific, 10270106) and 1X Penicillin-Streptomycin (Thermo Fisher Scientific, 15140163) at 37°C and 5% CO_2_. HEK293T TetOff-α-syn-A53T cells were cultured in 2 µg/mL Blasticidin S (Thermo Fisher Scientific, A1113903). HEK293T TetOff-α-syn-A53T EGFP-RNF152-IRES-mScarlet-I, HEK293T TetOff-α-syn-A53T mRuby3-Galectin-3 mClover3-Galectin-3, and HEK293T α-syn-A53T-mRuby3 α-syn-A53T-mClover3 cells were cultured in 2 µg/mL Blasticidin S and 225 µg/mL G418 (Thermo Fisher Scientific, 10131035). HEK293T TetOff-α-syn-A53T mNeonGreen-3K-1-10-IRES-mKate2 cells were cultured in 2 µg/mL Blasticidin S and 200 ng/mL Puromycin (Thermo Fisher Scientific, A1113803). Cells were passaged using TrypLE Express (Thermo Fisher Scientific, 12605036). Cultures were regularly checked for mycoplasma contamination.

Drug treatments were performed for the indicated amounts of time by addition of compounds dissolved in DMSO, except for LLOMe (Santa Cruz, sc-285992B) and Leupeptin (Sigma Aldrich, 11017101001), which were dissolved in PBS. Each time a drug treatment was performed, a control using an equal volume of the appropriate solvent (DMSO or PBS) was used for comparison.

HEK293T TetOff-α-syn-A53T cells were created by transduction with pTetOFF α-syn A53T virus (details about virus transduction below). The transduced cells were then selected with 10 µg/mL Blasticidin S. Monoclones were isolated by single-cell sorting with a BD FACS Aria III (BD Biosciences). HEK293T TetOff-α-syn-A53T EGFP-RNF152-IRES-mScarlet-I and HEK293T TetOff-α-syn-A53T mRuby3-Galectin-3 mClover3-Galectin-3 cell lines were produced in this genetic background by transfection with either pCMV-EGFP-RNF152-IRES-mScarlet-I or pCMV mRuby3-Galectin-3 and pCMV mClover3-Galectin-3, respectively, using Lipofectamine 3000 (Thermo Fisher Scientific, L3000008). Stably transfected polyclonal cell lines were then created by selection with 1 mg/mL G418. Monoclones of these two cell lines were then isolated by limiting dilution. The HEK293T TetOff-α-syn-A53T mNeonGreen-3K-1-10-IRES-mKate2 polyclonal cell line was created by transducing HEK293T TetOff-α-syn-A53T with pEF1a mNeonGreen-3K-B1-10-IRES-mKate2-2A-PuroR virus, and selecting with 2 µg/mL Puromycin.

HEK293T α-syn-A53T-mRuby3 α-syn-A53T-mClover3 cells were produced by transfection with pCMV α-syn-A53T-mRuby3 and pCMV α-syn-A53T-mClover3 with Lipofectamine 3000, followed by selection with 1 mg/mL G418. Monoclones were then isolated by single-cell sorting on a BD FACS Aria III.

Primary cortical neuron cultures were produced from E15.5 CD-1 wild-type mouse embryos. All mouse experiments were performed in accordance with the relevant guidelines and regulations of the Government of Upper Bavaria, Germany. Mice were maintained in a pathogen-free animal facility at 22 ± 1.5°C, 55 ± 5% humidity, with a 14/10 hr light/dark cycle. Mice were allowed food and water access ad libitum. Pregnant females were sacrificed by cervical dislocation. The uterus was then dissected from the sacrificed animal and submerged into a 10 cm sterile dish on ice filled with Dissection Medium: Hank’s balanced salt solution and 10 mM HEPES, 10 mM MgSO_4_, and 1X Penicillin-Streptomycin. Choosing sexes randomly, heads were cut from the embryos and the brains were removed and submerged into Dissection Medium on ice. Cortical hemispheres were then dissected and the meninges were removed. 6-7 embryos worth of cortical tissue was then digested in a 15 mL conical with 0.25% Trypsin-EDTA solution (Sigma Aldrich, T4049), 1 mM EDTA, and 15 μL DNAse I (Thermo Fisher Scientific, EN0521) at 37°C for 20 min. Trypsin was then quenched by washing the cortices settled at the bottom of the conical twice with Neurobasal medium (Thermo Fisher Scientific, A3582901) supplemented with 5% FBS. To generate a single-cell suspension, the digested tissue was then triturated 15 times through a 1 mL pipette tip in 2 mL Neurobasal medium. Cells were then centrifuged at 130 x g and resuspended in Primary Neuron Culture Medium: Neurobasal medium, 2% B-27 Plus (Thermo Fisher Scientific, A3582801), 1% L-Glutamine (Thermo Fisher Scientific, 25030081), and 1X Penicillin-Streptomycin. Neurons were then counted and plated either at 1 x 10^5^ neurons per well onto glass coverslips (VWR, 630-2190) in a 24-well plate or at 2 x 10^5^ neurons per well of a 12-well plate. In both cases, the coverslips or wells were pre-coated with 1 mg/mL poly-D-lysine (Sigma, A-003-E) and 1 µg/mL laminin (Thermo Fisher Scientific, 23017015). The day of plating was considered day in vitro (DIV) 0. A detailed protocol can be found at dx.doi.org/10.17504/protocols.io.ewov1ojwklr2/v1.

The KOLF2.1J AAVS1-TRE3G-NGN2 cell line was previously described^154^. During the generation of mutant cell lines in this background (described below), cells were cultured in StemFlex medium (Gibco, A3349401) at 37°C and 5% CO_2_ on plates coated with Matrigel (Corning, 354277). iPSCs were passaged using 0.5 mM Ultrapure EDTA pH 8 (Thermo Fisher Scientific, 15575020). Details about handling these cultures can be found at dx.doi.org/10.17504/protocols.io.j8nlkoq56v5r/v1.

### METHOD DETAILS

#### Recombinant protein purification and aggregation

Recombinant α-syn A53T and variants thereof were purified based on published protocol^155^. BL21(DE3) *E. Coli* were transformed with pT7-7 α-syn A53T (Addgene #105727; a gift of Hilal Lashuel^156^) or pT7-7 α-syn A53T D115A mNeonGreen-3K-B11 (this study). The culture was grown in Terrific Broth (Sigma, T0918) at 37°C until OD600 = 0.8, at which point the culture was induced with 1 mM IPTG for 4 hr. Cell pellets were harvested and lysed by sonication for 5 min in High Salt Buffer: 750 mM NaCl, 50 mM Tris, 1 mM EDTA, cOmplete EDTA-free Protease Inhibitor (Roche, 5056489001), pH 7.6. The lysate was then boiled for 15 min, cooled on ice, and clarified by centrifugation at 6,000 x g for 20 min at 4°C. The clarified lysate was then dialyzed into Size Exclusion Buffer, consisting of 10 mM Tris, 50 mM NaCl, 1 mM EDTA, pH 7.6. The dialyzed protein was then purified further by size exclusion chromatography in a Superdex 200 column (GE, 28989335). Following size exclusion, the protein was dialyzed into Anion Exchange Buffer containing 10 mM Tris, 25 mM NaCl, 1 mM EDTA, pH 7.6. The dialyzed protein was run on a MonoQ column (Amersham, 17-0506-01) and eluted with a linear gradient of 25 mM to 1 M NaCl in Anion Exchange Buffer. The anion-exchanged protein was then dialyzed into 50 mM Tris, 150 mM KCl, pH 7.5, concentrated to 5 mg/mL, frozen, and stored at -80°C until use. A detailed protocol can be found at dx.doi.org/10.17504/protocols.io.btynnpve.

α-Syn PFFs were produced based on an established protocol^95^. 5 mg/mL recombinant α-syn A53T or α-syn A53T D115A mNeonGreen-3K-B11 was centrifuged for 1 hr at 100,000 x g at 4°C. The supernatant was then added to a new 1.5 mL tube and shaken for 24 hr in a Thermomixer (Eppendorf) at 900 rpm and 37°C.

α-Syn PFFs were labeled with Alexa Fluor 647 NHS Ester (Thermo Fisher Scientific, A20006) according to the manufacturer’s instructions. 5 mg of PFFs were pelleted by 30 min of 20,000 x g centrifugation and resuspended in 0.1 M NaHCO_3_, pH 8.3. The PFFs were then reacted with 0.5 mg of Alexa Fluor Ester dissolved in DMSO (Thermo Fisher Scientific, D12345) for 1 hr at room temperature with shaking at 400 rpm, protected from light. The PFFs were then washed by 5 rounds of pelleting with a 20,000 x g centrifugation for 30 min and resuspension in 0.1 M NaHCO_3_, pH 8.3. After the final wash, the PFFs were finally resuspended in PBS, pH 7.2 (Thermo Fisher Scientific, 20012068) at a concentration of 5 mg/mL.

CHMP2B was expressed as a His-SUMO fusion protein in BL21(DE3) *E. Coli* transformed with pET28-His-SUMO-CHMP2B (this study). After outgrowth at 30°C, the culture was shifted to 18°C and induced with 0.5 mM IPTG for 16 hr. Pellets were lysed by sonication for 20 min in Lysis Buffer containing 50 mM Tris, 500 mM NaCl, 10 mM imidazole, 2 mM β-Mercaptoethanol (β-ME), 5% glycerol, cOmplete EDTA-free Protease Inhibitor, pH 8. The lysate was then clarified by centrifugation and run over 3 connected 5 mL HisTrap columns (Cytiva, 17524802), followed by elution with a gradient of 50 mM to 500 mM imidazole in a buffer of 50 mM Tris, 500 mM NaCl, 2 mM β-ME, pH 8.0. To cleave the His-SUMO tag, the eluted His-SUMO-CHMP2B was then combined with His-SenP2 protease (7500 U per L of culture, Max Planck Institute of Biochemistry Core Facility) and glycerol to 10%; the mixture was dialyzed overnight in 20 mM Tris, 150 mM NaCl, 4 mM imidazole, 2 mM β-ME, pH 7.5. To separate the cleaved CHMP2B from His-SUMO, a reverse HisTrap column was then performed as above with two alterations: the buffers were instead at pH 7 and the flow-through was collected instead of the eluate. The CHMP2B in the flow-through was then subjected to size exclusion chromatography in a Superdex 200 column using 20 mM Tris, 150 mM NaCl, 2 mM β-ME, pH 7. Glycerol was added to the eluted CHMP2B to a final concentration of 10%, then the sample was concentrated to 1.11 mg/mL. A detailed protocol can be found at dx.doi.org/10.17504/protocols.io.bp2l623r1gqe/v1.

Aβ42 was purified from inclusion bodies as in the “without urea” protocol described in ref.^157^. BL21(DE3) *E Coli* were transformed with the pET-Sac-Abeta(M1-42) plasmid (Addgene #71875; a gift of Sarah Linse^158^), grown in LB media at 37°C, and induced with 0.4 mM IPTG for 4 hr. Bacterial pellets from a 3 L culture were sonicated in 50 mL TE7.5 Buffer (10 mM Tris, 1 mM EDTA, cOmplete EDTA-free Protease Inhibitor, 2 U/mL Benzonase (Max Planck Institute of Biochemistry Core Facility), pH 7.5) until the sample appeared homogenous. To clean the pellets, the lysate went through two washing steps consisting of centrifugation at 18,000 x g for 8 min at 4°C, discarding of the supernatant, and sonication of the pellet in TE7.5 Buffer. The sample was then pelleted, resuspended in 40 mL TE9.5 Buffer (10 mM Tris, 1 mM EDTA, pH 9.5), sonicated, and centrifuged at 18,000 x g for 10 min at 4°C. The supernatant was filtered and adjusted to pH 8.5 with 1 M HCl. The pH-adjusted supernatant was then passed through a self-packed column of DEAE sepharose (GE, 17-0709-01) equilibrated with TE8.5 Buffer (10 mM Tris, 1 mM EDTA, pH 8.5). Aβ42 was then eluted with TE8.5 containing 50 mM NaCl. The eluted Aβ42 then underwent buffer exchange into TE8.5 Buffer via a desalting column (Cytiva, 17508702). The buffer-exchanged Aβ42 was then lyophilized.

To prepare Aβ42 monomers and aggregates, 500 µg of lyophilized Aβ42 was resuspended in 700 µL 6M GuHCl, 20 mM sodium phosphate, pH 8.5. The solubilized Aβ42 was then subjected to size exclusion chromatography on a Superdex 75 column (GE, GE17-5174-01) in Aβ42 Aggregation Buffer containing 20mM sodium phosphate, 0.2 mM EDTA, pH 8.5. The eluted monomeric Aβ42 was pooled and its concentration was determined by 205 nm absorbance on a Nanodrop One (Thermo Fisher Scientific). The concentration of Aβ42 monomers was adjusted to 10 µM in Aβ42 Aggregation Buffer. Part of the pool of Aβ42 monomers was snap-frozen in low-bind tubes (Eppendorf, 0030108116) and stored, while the rest was aggregated. To aggregate Aβ42, the monomers were aliquoted into a 96-well plate (Corning, 3881) at 80 µL per well, and incubated at 37°C for 24 hr. The aggregates were then snap-frozen and stored at -80°C.

To purify Cpf1, a 500 mL culture of Rosetta (DE3)pLysS *E. Coli* cells (Merck, 70956) transformed with the pDEST-his-AsCpf1-EC plasmid^159^ was grown at 37°C in LB media containing 50 μg/mL chloramphenicol and 34 μg/mL kanamycin with shaking until OD600 reached 0.6-1. The culture was cooled to <20°C in an ice bath before induction with 0.5 mM IPTG, followed by overnight shaking at 16–20°C. Cells were harvested by centrifugation and stored at −80°C. The cell pellet was resuspended in 25 mL lysis buffer (50 mM HEPES-NaOH, 150 mM NaCl, 5 mM MgCl2, 20 mM imidazole, 0.5 mM TCEP, pH 7.0). The resuspended cells were lysed by the addition of 2.5 mL 10X FastBreak buffer (Promega, V8571). Lysis was enhanced by 15 min Benzonase (Merck, 71205-M) treatment in 4 µL increments, as needed, to reduce viscosity. Once DNA was digested, the salt concentration was then adjusted to 500 mM by adding 5 M NaCl. Insoluble material was removed by centrifugation at 38,000 x g for 10 min at 4°C. 4 mL pre-washed Ni-NTA resin (Thermo Fisher Scientific, 88221) was equilibrated in a buffer of 50 mM HEPES-NaOH, 500 mM NaCl, 5 mM MgCl2, 20 mM imidazole, pH 7.0. Clarified lysate was bound to this resin at 4°C for 1 hr with gentle rotation. The slurry was loaded into a disposable column (Bio-Rad, 7321010) and washed three times with 20 mL high-salt buffer (50 mM HEPES-NaOH, 2 M NaCl, 20 mM imidazole, 0.5 mM TCEP, pH 7.0) and twice eluted with 5 mL elution buffer (50 mM HEPES-NaOH, 5 mM MgCl2, 400 mM imidazole, 0.5 mM TCEP, pH 7.0). Eluates were diluted 4-fold with PBS and loaded onto a Hitrap Heparin HP Column (Sigma, 17-0407-03) pre-equilibrated with PBS pH 7.4. The heparin column was washed with 20 mL of 0.1 M NaCl and eluted sequentially with 10 mL of PBS pH 7.4 containing increasing NaCl concentrations (0.3 M, 0.5 M, 0.7 M, 1.0 M, 2.0 M). Fractions judged by SDS-PAGE to contain purified Cpf1 were pooled and concentrated to 3 mL using a 50 kDa MWCO Centriprep YM-50 column (Merck, 4311). The sample was twice brought to 15 mL in PBS with 20% Glycerol and 2 mM TCEP, then concentrated to 3 mL with the Centriprep column. The final preparation was briefly centrifuged, sterilized using a syringe filter, and quantified by 280 nm absorbance on a NanoDrop. Purified protein was stored at 4 mg/mL at −80°C.

#### iPSC gene editing

Mutations were made by gene editing, performed as described in^160^ using a detailed protocol that can be found at: dx.doi.org/10.17504/protocols.io.q26g71q68gwz/v1. To target the appropriate region of the CHMP2B gene, the SpCas9 sgRNA was generated by using the GeneArt Precision gRNA synthesis kit (Thermo Fisher Scientific) and the Cas12a (Cpf1) sgRNAs were ordered from Integrated DNA Technologies (IDT). The sequences of these sgRNAs (5’ to 3’) are as follows: GCCAAACAACTTGTGCATCTACGG [CHMP2B^-/-^, CRISPR-Cas12a (Cpf1)], ATCAAGAACTTGATTCACAATATC [CHMP2B^Q165X/+^, CRISPR-Cas12a (Cpf1)], and AGAGTTACGAGGTACACAGA (CHMP2B^I29V/I29V^, CRISPR-SpCas9). Ultramer ssDNA oligos (IDT) were also designed to act as a homology-directed repair template in generating the point mutants, possessing the following sequences (bases that diverge from the genomic sequence capitalized): ccaactaagaaaagatgatgttcatacctttccagaaatttcaattccaatCtcatcaagaacttAattcacaatatcctggctttcttcttcgtcatcagaaccgtcaaagatgtc (CHMP2B^Q165X/+^), and ctcctagatgtaataaaggaacagaatcgagagttacgaggtacacagagAgctataGtcagagatcgagcagctttagagaaacaagaaaaacagctggtaagtag (CHMP2B^I29V/I29V^). Information about oligonucleotides used in gene editing can be found in Table S1.

Gene edits were introduced by first incubating 0.6 μg sgRNA with 3 μg SpCas9-NLS protein (UC Berkeley QB3 Macrolab) or 80 pmol sgRNA with 63 pmol Cas12a (Cpf1) protein for 10 min at room temperature, then electroporating the mixture into 2 x 10^5^ KOLF2.1J AAVS1-TRE3G-NGN2 cells using a Neon Transfection System (Thermo Fisher Scientific). For the CRISPR-Cas12a (Cpf1) system, the cells were co-transfected with 39 pmol Alt-R Cpf1 Electroporation Enhancer (IDT, 1076301). For the point mutants, 80 pmol of the appropriate Ultramer ssDNA oligo was added to the electroporated CRISPR/Cas protein mixture. Cells then recovered for 1 day before single cells were sorted with a SH800S Cell Sorter (Sony Biotechnology).

Clones were then grown for 7-12 days before validation by DNA sequencing on a MiSeq (Illumina) as well as Sanger sequencing of PCR-amplified regions. The primer pairs used for amplification were: AAGAAAATGGCCAAGATTGGTA & CCATCTTCATTTGGGAATTCAT (CHMP2B^-/-^), TTGATGACATCTTTGACGGTTC & GAAATAAAAACCATGCACCTCC (CHMP2B^Q165X/+^), and GGTTTCTTTTGTGATTCTCCTAG & CATGTGCCTTCTTCCTAGTTAGC (CHMP2B^I29V/I29V^). Information about these genotyping oligonucleotides appear in Table S1. CHMP2B protein expression in sequencing-validated clones was then assessed by immunoblot.

#### iNeuron Differentiation

KOLF2.1J AAVS1-TRE3G-NGN2 background iPSCs were differentiated into iNeurons using the forced NGN2 expression protocol^134^. Prior to differentiation, iPSC lines were cultured in mTeSR Plus (Stem Cell Technologies, 100-0276) on Matrigel-coated plates at 37°C and 5% CO_2_. The cells were passaged with ReLeSR (Stem Cell Technologies, 05873). For each differentiation, a single cell suspension of the iPSCs was generated by Accutase (Stem Cell Technologies, 07920) treatment and 2.5 x 10^5^ cells were plated per well in a Matrigel-coated 6-well plate into ND1 medium: DMEM/F12, 1X NEAA (Gibco, 11140-068), 1X N2 (Thermo Fisher Scientific, 17502048), 10 ng/mL NT3 (Peprotech, AF-450-03-10), 10 ng/mL BDNF (Peprotech, 450-02), 200 ng/mL laminin (Bio-techne, 3446-005-01), and 2 μg/mL doxycycline (Clontech, 631311), supplemented with 10 μM Y-27632 (Biozol, S1049). The day of this first plating was considered DIV 0. On DIV 1, the media was exchanged for ND1 without Y-27632. On DIV 2, the media was exchanged for ND2 medium: Neurobasal, 1X Glutamax (Thermo Fisher Scientific, 35050061), 2% B-27 Plus, 10 ng/mL NT3, 10 ng/mL BDNF, and 2 μg/mL doxycycline. Half of the media was exchanged with fresh ND2 on DIV 4. On DIV 6, cells were split using Accutase and plated at 5 x 10^4^ cells per well onto coverslips coated with 0.01% poly-L-ornithine (Sigma-Aldrich, P3655-50MG) and 10 μg/mL laminin in a 24-well plate. Half of the media was exchanged every 2 days thereafter with fresh ND2 media, with the media lacking doxycycline from DIV10 onwards. After addition of PFFs, the media was additionally supplemented with 1X Penicillin-Streptomycin. A detailed protocol for this differentiation can be found at dx.doi.org/10.17504/protocols.io.x54v9p8b4g3e/v1.

#### Transfections, transductions, and knockdowns

For experiments involving expression of CHMP2B variants with their related controls or Cas9 with an sgRNA, plasmids were transfected using Fugene 6 (Promega, E2692). Cells at 70-80% confluence were transfected with a 4:1 (μL:μg) ratio of Fugene 6 to plasmid. Cells were replated the following day to avoid overgrowth of the culture.

For the strong CHMP2A and CHMP2B knockdown, the *CHMP2B* locus was first disrupted by transfection with an all-in-one plasmid encoding both Cas9 and an sgRNA with the sequence AAUUCCCAAAUGAAGAUGGC, pSPcas9(BB)-2A-Puro V2.0 sgCHMP2B-2. As a non-targeting control, cells were transfected with the parental pSPcas9(BB)-2A-Puro V2.0 plasmid (Addgene # 62988; a gift of Feng Zhang^161^). The cells were then selected with 2 μg/mL puromycin for 2 days. CHMP2A was then knocked down by reverse transfection; 1.5 x 10^5^ cells were plated into the wells of a 12-well plate pre-loaded with reverse transfection mix: 12.5 pmol siPOOL siRNA against CHMP2A (siTOOLs Biotech, si-G020-27243) or a non-targeting negative control (siTOOLs Biotech, si-C002), 0.94 μL Lipofectamine 3000, and 36.9 μL Opti-MEM (Thermo Fisher Scientific, 31985-062). The media was exchanged the following day and measurements were taken after 72 hr of knockdown.

For the mild CHMP2A and CHMP2B knockdown, the *CHMP2B* locus was disrupted as above except for the puromycin selection, which was instead performed for two days. CHMP2A was then knocked down by transfecting a 60-70% confluent well of a 12-well plate with 100 pmol of ON-TARGETplus Human CHMP2A SMARTPool siRNA (Horizon Discovery, L-020247-01-0005) or ON-TARGETplus Non-targeting Pool siRNA (Horizon Discovery, D-001810-10-05) and 7.5 μL Lipofectamine 3000. Cells were then replated and measured 96 hr after knockdown.

To produce lentivirus, 2.5 x 10^6^ Lenti-X 293T cells (Takara, 632180) were plated onto a 10 cm plate. The following day, cells were transfected with 0.65 pmol psPAX2 (Addgene #12260; a gift of Didier Trono), 0.36 pmol pMD2.G (Addgene #12259; a gift of Didier Trono), and 0.82 pmol of the desired lentiviral transfer vector using X-tremeGENE HP DNA Transfection Reagent (Sigma Aldrich, 6366244001) at a ratio of 4 μL per μg of transfer vector. The transfer vectors used were either pLJC5-Tmem192-3xHA (Addgene #102930; a gift of David Sabatini^125^), pTetOFF α-syn A53T (this study), or pEF1a mNeonGreen-3K-B1-10-IRES-mKate2-2A-PuroR (this study). The following day, the media was exchanged with 9 mL of fresh DMEM. 72 hr after transfection, the media was collected, filtered, and mixed with 3 mL Lenti-X concentrator (Takara, 631231), and incubated at 4°C overnight. The viral precipitate was then concentrated by pelleting at 1500 x g for 45 min at 4°C and resuspending in 120 μL ice-cold PBS. This concentrated virus was then frozen and stored at -80°C. A detailed protocol can be found at: dx.doi.org/10.17504/protocols.io.n92ldmdyol5b/v1.

To create stable HEK293T cell lines (detailed above in “Cell culture”), cells were plated in 24-well plates and treated with 10 μL of concentrated virus and 0.5 μg/mL polybrene (Sigma Aldrich, TR-1003-G).

#### PFF treatment and induction of α-syn aggregation

For Lipofectamine 3000-mediated α-syn aggregation induction, HEK293T cells were first plated at 250 cells/mm^2^ (e.g. 10^5^ cells per well of a 12-well plate) for flow cytometry, immunoblot, or immunoprecipitation experiments or 125 cells/mm^2^ (e.g. 2.5 x 10^4^ cells per well of a 24-well plate) for microscopy experiments. On the following day, a 5 mg/mL aliquot of PFFs was thawed, diluted 1:20 in PBS, and sonicated in a BioRuptor Plus (Diagenode, B01020001) for 25 cycles of 5 seconds on and 5 seconds off. Opti-MEM and Lipofectamine 3000 were then mixed at a ratio of 50:3 and incubated for 5 min. In no Lipofectamine 3000 controls, an equal volume of Opti-MEM served as a substitute. The sonicated PFFs were then added to this pre-incubated mixture at a ratio of 50:3:20 (Opti-MEM:Lipofectamine 3000:diluted PFFs) to form the “seeding mixture.” In no PFF controls, the PFFs were substituted by an equal volume of PBS. The seeding mixture was incubated for 10 min at room temperature, then added dropwise to wells. The volume of seeding mixture used was adjusted such that 1 μL of sonicated PFFs were added per 10 mm^2^ of well surface area (e.g. 73 μL of seeding mixture used in 24-well plates and 146 μL in 12-well plates). The following day, the media was exchanged with fresh DMEM to minimize off-target effects of Lipofectamine 3000. Measurements were then taken 2 days after PFF treatment. A detailed protocol can be found at: dx.doi.org/10.17504/protocols.io.eq2lyjbbplx9/v1.

Extended PFF treatment in HEK293T cells without Lipofectamine 3000 was performed exactly as described above with 4 alterations: 1) cells were instead plated at 32 cells/mm^2^, 2) Lipofectamine 3000 was not used, 3) rather than a media exchange, half the volume of medium was spiked in after 3 days, and 4) measurements were taken 6 days after PFF treatment. PFF treatment in primary mouse neurons and iNeurons occurred as described above, although the plating density was as described in the “Cell culture” and “iNeuron differentiation” sections, Opti-MEM was replaced with the appropriate growth medium, Lipofectamine 3000 was not used, and there was no full media exchange after PFF treatment. For iNeurons, instead of exchanging the media in the first media change after PFF treatment, half the well volume of media was spiked in and none was removed. In primary neuron experiments, PFFs were added on different days such that all measurements were taken on DIV 18 (e.g. PFFs added on DIV 4 for 14 day treatment and on DIV 11 for 7 day treatment). iNeurons were treated with PFFs on DIV 16.

#### Immunofluorescence Microscopy

For immunofluorescence microscopy, HEK293T cells were grown on poly-L-lysine-coated coverslips (Neuvitro, GG-12-1.5-PLL), iNeurons were grown on poly-L-ornithine- and laminin-coated coverslips (described above), and primary neurons were grown on poly-D-lysine- and laminin-coated coverslips (described above). Cells growing on coverslips were fixed by 4% PFA (Thermo Fisher Scientific, 28908) for 10 min at room temperature. Permeabilization was then carried out in 0.1% Triton X-100 in PBS for 5 min at room temperature. The cells were blocked for 1 hr in a Blocking Solution consisting of PBS, 5% milk powder (Sucofin), 0.1% Triton X-100, pH 7.4. Primary antibodies were diluted in Blocking Solution and incubated on the coverslips overnight at 4°C. All primary antibodies were used at 1:500 dilution, except for anti-MAP2 (Merck Millipore, AB5543) at 1:1500 and anti-CHMP3 (Santa Cruz, sc-166361) at 1:100. Coverslips were then washed three times in PBS before incubation with secondary antibodies diluted in Blocking Solution for 3 hr at room temperature protected from light. Secondary antibodies (listed in the STAR Resources table) were diluted 1:1000. Coverslips were then washed with PBS, then nuclei were stained with NucBlue (Thermo Fisher Scientific, R37606) for 5 min, followed by two more PBS washes. Dako Fluorescence Mounting Medium (Agilent, S3023) was used to mount coverslips on Epredia glass slides (Thermo Fisher Scientific, 17294884). A detailed protocol can be found at: dx.doi.org/10.17504/protocols.io.e6nvwd7n7lmk/v1.

Images were acquired either with a FEI CorrSight using a 63X oil objective running FEI MAPS 2.1 or 3.8 (FEI), a Leica SP8 Falcon using a 63X oil objective running Leica Applications Suite X 3.5.7.23225 (Leica; Max Planck Institute of Biochemistry Imaging Facility), a Zeiss LSM800 using a 40X Plan-Apochromat objective in Airyscan super resolution mode running Zen 2.6 (Zeiss), or a W1 spinning disk confocal (Yokogawa) on an Eclipse Ti-2 (Nikon) with a 100X oil Plan-Apochromat objective (Harvard Medical School Core for Imaging Technology & Education) running NIS-Elements 5.21.03.

Prior to further analysis, Airyscan images acquired on the LSM980 were first processed in ZEN software (Zeiss). For images generated on the CorrSight, LSM800, and Eclipse microscopes, figure panels of micrographs were prepared in Fiji^162^. Images acquired on the SP8 falcon were processed using a Matlab script, deposited at https://github.com/csitron/Microscopy_lif_Adjust_MATLAB. Quantification of pSer129 α-syn area compared to nuclear (DAPI) area was done in Matlab using the script deposited at https://github.com/csitron/Microscopy_lif_Channel_Quant_MATLAB.

#### Immunoblotting

For harsh lysis, HEK293T cells were harvested by trypsinization and lysed in RIPA Buffer (Thermo Fisher Scientific, 89900) supplemented with cOmplete Mini EDTA-free Protease Inhibitor Cocktail (Roche, 04693159001) and 7.5 U/mL Benzonase on ice. The lysate was then sonicated in a BioRuptor Plus for 5 cycles of 30 seconds on and 30 seconds off. The concentration was then determined using the Pierce Rapid Gold BCA Protein Assay Kit (Thermo Fisher Scientific, A53225) and equalized between samples. The lysate was then denatured by addition of one third of the sample volume of 4X NuPAGE LDS Sample Buffer (Thermo Fisher Scientific, NP0007) supplemented with 5% β-ME and boiling for 5 min at 95°C.

Primary neurons were lysed in the growth vessel by first washing with ice-cold PBS, then direct addition of the RIPA Buffer described above. The lysate was then sonicated and the concentrations were then equalized as described above. Total protein was precipitated from the lysate with 10% TCA. The protein pellets were then washed twice with ice-cold acetone and dried. The dried pellets were rehydrated with 300 mM Tris, pH 8.3, then treated with an equal volume of 4X NuPAGE LDS Sample Buffer with 5% β-ME and heated for 10 min at 70°C.

Samples were then run on NuPAGE 1.5 mm 4-12% Bis-Tris gels (Thermo Fisher Scientific, NP0323BOX) along with the PageRuler Prestained Protein Ladder (Thermo Fisher Scientific, 26617) and transferred to a PVDF membrane (Sigma Aldrich, 3010040001) for 1 hr at 100 V in Towbin Buffer (25 mM Tris, 192 mM glycine, pH 8.3) containing 20% methanol. For membranes stained for untagged α-syn, the membrane was fixed in 4% PFA for 30 min at room temperature after transfer.

To perform filter trap assays, cells were collected by trypsinization and lysed in 50 mM Tris, 150 mM NaCl, 1% Triton X-100, pH 7.5 supplemented with cOmplete Mini EDTA-free Protease Inhibitor Cocktail and 7.5 U/mL Benzonase. The lysate was then sonicated in a BioRuptor Plus for 3 cycles of 30 seconds on and 30 seconds off. The sonicated lysate underwent gentle clarification by centrifugation at 500 x g for 5 min at 4°C. After performing a BCA Gold assay to determine the protein concentration, the concentrations were then equalized and loaded onto a 0.2 µm pore size cellulose acetate membrane (GE, 10404131) in a slot blot vacuum manifold (Hoefer, PR648). A small sample of the lysate was then denatured with 4X NuPAGE LDS Sample Buffer with 5% β-ME and processed for SDS-PAGE to run as a loading control. More details can be found at: dx.doi.org/10.17504/protocols.io.x54v92re4l3e/v1.

PVDF and cellulose acetate membranes were blocked for 1 hr at room temperature in Blocking Buffer: 5% milk powder in TBS-T (50 mM Tris, 150 mM NaCl, 0.1% Tween-20, pH 7.4). Membranes were then probed with primary antibodies diluted in Blocking Buffer overnight at 4°C. Concentrations of primary antibodies were as follows: 1:1000 rabbit anti-CHMP2A (Proteintech, 10477-1-AP), 1:2000 rabbit anti-CHMP2B (Abcam, ab33174), 1:2000 mouse anti-α-synuclein MJFR1 (Abcam, ab138501), 1:2000 mouse anti-α-synuclein LB509 (Abcam, ab27766), 1:2000 rabbit anti-α-synuclein phosphoSer129 EP1536Y (Abcam, ab51253), 1:500 mouse anti-β-Amyloid (Biolegend, 803001), 1:100 mouse anti-LAMP1 (Developmental Studies Hybridoma Bank, H3A3), 1:2000 mouse anti-Ubiquitin (Santa Cruz, sc-8017), 1:2000 rat anti-GFP (Proteintech, 3h9-150), 1:5000 mouse anti-β-actin (Abcam, ab6276), 1:2000 mouse anti-GAPDH (Merck Millipore, MAB374), 1:1000 mouse anti-Hsp60 (Abcam, ab59458), 1:1000 rabbit anti-Calreticulin (Cell Signaling Technologies, 12238S), 1:1000 mouse anti-Calnexin (Santa Cruz, sc-46669), 1:1000 mouse anti-α-tubulin (Sigma Aldrich, T6199). After three washes in TBS-T, the membrane was then stained with either 1:2000 anti-mouse HRP (Sigma Aldrich, A4416), 1:10000 anti-rabbit HRP (Sigma Aldrich, A9169), 1:2000 anti-rat HRP (Sigma Aldrich, A9037), or 1:2000 Veriblot (Abcam, ab131366) diluted in Blocking Buffer.

Stained membranes were then washed, developed with either Immobilon Forte or Classico Western HRP Substrate (Merck Millipore, WBLUF0500 or WBLUC0500), and imaged on an Amersham ImageQuant 800 biomolecular imager (Cytiva). Before re-probing previously-stained membranes, the membranes were first stripped using Restore Western Blot Stripping Buffer (Thermo Fisher Scientific, 21059) and then blocked and stained as described above.

Immunoblots were quantified using a custom Matlab script that has been deposited at https://github.com/csitron/Western-Blot-Quantification-in-MATLAB.

#### Immunoprecipitations and Pulldowns

For immunoprecipitation of CHMP2B from lysate, cells were collected by trypsinization and lysed in PBS, 0.1% Triton X-100, 0.02% Tween-20, pH 7.2 with cOmplete Mini EDTA-free Protease Inhibitor Cocktail and 0.75 U/mL Benzonase on ice. The affinity matrix was prepared by binding 2.67 µL of rabbit anti-CHMP2B to 30 µL Protein G Dynabeads (Invitrogen, 10007D), then blocking the antibody-bound beads with PBS, 0.1% Triton X-100, 0.02% Tween-20, 3% BSA, pH 7.2 with cOmplete Mini EDTA-free Protease Inhibitor Cocktail. These beads were then incubated with 200 µg of lysate with an additional 113 mM NaCl (for a total of 250 mM NaCl) for 1 hr at 4°C with rotation. The beads were then washed three times in PBS, 0.1% Triton X-100, 0.02% Tween-20, 113 mM additional NaCl, pH 7.2. The bound protein was eluted by boiling with 2X NuPAGE LDS Sample Buffer with 2.5% β-ME at 95°C for 5 min. The eluate was denatured again at 95°C for 5 min after addition of an equal volume of 10 mM DTT in the lysis buffer. To resolve CHMP2B and a nonspecific band in the eluate, we found it critical to run the input and elution fractions on 12% Bis-Tris NuPAGE gels (Thermo Fisher Scientific, NP0342BOX). A detailed protocol has been uploaded to dx.doi.org/10.17504/protocols.io.eq2lywe9pvx9/v1.

To immunoprecipitate ubiquitinated proteins, cell pellets were lysed on ice in 20 mM Tris, 150 mM NaCl, 1% Triton X-100, pH 8 supplemented with cOmplete Mini EDTA-free Protease Inhibitor Cocktail, freshly-added 20 mM N-ethylmaleimide (Sigma Aldrich, E3876-25G), 50 µM PR-619 (Sigma Aldrich, 662141-25MG), and 7.5 U/mL Benzonase. Lysates were sonicated in a BioRuptor Plus for 5 cycles of 30 seconds on and 30 seconds off and clarified at 18000 x g for 10 min at 4°C. 1 mg of lysate was incubated with 50 µL of pre-equilibrated Ubiquitin pan-selector resin (NanoTag Biotechnologies, N2510) for 1 hr at 4°C with rotation. The resin was washed 3 times in 20 mM Tris, 150 mM NaCl, 1 mM EDTA, 1% Triton X-100, 0.1% SDS, pH 8 with an additional 50 µM PR-619. Elution was then carried out by boiling for 5 min at 95°C in 100 µL 2X NuPAGE LDS Sample Buffer with 2.5% β-ME. A detailed protocol appears at: dx.doi.org/10.17504/protocols.io.dm6gp36k1vzp/v1.

Lysis prior to LysoIP^125^ was carried out by subjecting cell pellets from a 10 cm dish to 30 strokes of a Dounce Homogenizer in 1 mL of ice-cold KPBS (10 mM KH_2_PO_4_, 136 mM KCl, pH 7.25 with cOmplete Mini EDTA-free Protease Inhibitor Cocktail). The lysate was then clarified by centrifugation at 1000 x g for 4 min at 4°C. 100 µg of the clarified lysate was bound to 75 µL of pre-equilibrated anti-HA magnetic beads (Thermo Fisher Scientific, 88836) at 4°C for 20 min. After 3 washes with KPBS, the bound proteins were eluted by addition of 70 µL RIPA with cOmplete Mini EDTA-free Protease Inhibitor Cocktail at 4°C for 30 min. An in-depth protocol is described at: dx.doi.org/10.17504/protocols.io.yxmvmep25g3p/v1.

For the in vitro co-immunoprecipitation between CHMP2B and α-syn, 5 µM CHMP2B was mixed with either 5 µM α-syn PFFs or 5 µM α-syn monomers in PBS-T: PBS, 0.02% Tween-20, pH 7.2. Prior to use, both CHMP2B and α-syn monomers were centrifuged at 16,000 x g for 10 min at 4°C to remove any aggregated material. 250 µL of these protein mixtures and control mixtures thereof (i.e. without CHMP2B) were added to 20 µL of Protein G Dynabeads that had been previously complexed with 2 µL of rabbit anti-CHMP2B antibody (Abcam, ab33174). The protein mixtures were allowed to bind the beads for 45 min at room temperature with rotation. The beads were then washed 5 times with PBS-T and eluted with 50 µL 2X NuPAGE LDS Sample Buffer with 2.5% β-ME at 70°C for 10 min. More details can be found at: dx.doi.org/10.17504/protocols.io.8epv5rb66g1b/v1.

The biotinylated CHMP2B_55-96_ peptide pulldown was carried out by first conjugating 3.2 pmol of a peptide with the sequence Biotin-GSKEACKVLAKQLVHLRKQKTRTFAVSSKVTSMSTQTKVMNSQM-NH_2_ (Max Planck Institute of Biochemistry Bioorganic Chemistry & Biophysics Core Facility) dissolved in 45% ethanol, 10% acetic acid, and 1 mM DTT to 25 µL Pierce Magnetic Streptavidin Beads (Thermo Fisher Scientific, 88816). These beads were then blocked in PBS, 1% Triton X-100, additional 113 mM NaCl (final concentration 250 mM), 3% BSA, pH 7.2. To eliminate aggregates from the non-aggregated species, monomeric Aβ42 and α-syn were centrifuged at 20,000 x g for 30 min at 4°C prior to use. The blocked peptide-conjugated beads (or peptide-free control) were incubated with of 2.5 µM Aβ42 monomers, Aβ42 aggregates, α-syn monomers, or α-syn PFFs for 1 hr in a buffer consisting of PBS, 1% Triton X-100, additional 113 mM NaCl, 0.1% SDS, 1 mM DTT, pH 7.2. In the case of the Aβ42 monomers, incubation was carried out at 4°C to reduce aggregation; incubations otherwise occurred at room temperature. α-syn samples were eluted by boiling at 95°C for 10 min in 50 µL 1X NuPAGE LDS Sample Buffer with 1.25% β-ME. Aβ42 samples were eluted with 100 µL formic acid at 19°C for 40 min.

The formic acid was then evaporated in a SpeedVac at 45°C for 1 hr. The dried protein pellet was re-solubilized by incubation in 50 µL HU Buffer (8 M urea, 5% SDS, 200 mM Tris, 1 mM EDTA, 0.01% bromophenol blue, pH 6.8) with 5% β-ME for 10 min at 60°C. A detailed protocol can be found at: dx.doi.org/10.17504/protocols.io.261ge5wrwg47/v1.

#### Flow Cytometry

The cells measured for flow cytometry were grown in 12-well plates. Two experiments required unique replating and washing conditions. First, for the assay measuring disappearance of Gal3 FRET signal after LLOMe treatment in cells with or without aggregates (Figure 5D), α-syn aggregation was first induced in the Gal3 FRET cell line as previously described. On the following day, the cells were trypsinized and replated at a density of 1.5 x 10^5^ cells per well in 12-well plates that had been coated with 0.05 mg/mL poly-D-lysine (Thermo Fisher Scientific, A3890401). The day after replating, the media was replaced with media containing 300 µM LLOMe for 2 hr. The cells were then washed once with DMEM lacking FBS, then fresh medium was added. After the indicated amounts of time, the cells were fixed and prepared for flow cytometry as described below. For the quantification of PFF leakage in cells with or without aggregates (Figure 6G), aggregation was first induced in HEK293T TetOff-α-syn-A53T mNeonGreen-3K-1-10-IRES-mKate2 cells. The cells were then trypsinized and replated on the following day at a density of 10^5^ cells per well in 0.05 mg/mL poly-D-lysine-coated 12-well plates. Half of the wells containing replated cells were then treated with 100 µL Opti-MEM and either 40 µL of 1:20 diluted, sonicated α-syn PFFs carrying the D115A mutation and C-terminally tagged with mNeonGreen_11_. The other half of the wells were treated with 100 µL Opti-MEM and 40 µL PBS. The mNeonGreen_11_-tagged PFFs carried the D115A mutation to limit C-terminal cleavage^163^, and thereby ensure that the mNeonGreen_11_ tag remained attached to the PFFs. Two days after addition of these PFFs, the cells were then processed for flow cytometry. A detailed protocol is described at: dx.doi.org/10.17504/protocols.io.q26g71dr8gwz/v1.

Cells were collected for flow cytometry by trypsinization, pelleted by centrifugation at 1,500 x g for 3 min at 4°C, washed once in ice-cold PBS, then fixed in 4% PFA for 10 min at 37°C. The fixed samples were then resuspended in PBS and stored at 4°C before measurement. At least two technical replicates were collected within each experiment. Experiments conducted on different days were considered biological replicates.

Flow cytometry analysis was performed on an Attune NxT Flow Cytometer (Thermo Fisher Scientific) running Attune Cytometric Software 5.1.2111.1. mTagBFP2 fluorescence was measured by excitation with a 405 nm laser and recorded with a 440/50 filter (corresponding to the VL1-A channel). EGFP, mClover3, and split mNeonGreen-3K used a 488 nm laser and 530/30 filter (BL1-A), while mScarlet-I used a 561 nm laser and a 620/15 filter (YL2-A). FRET between mClover3 and mRuby3 was measured by via 488 nm excitation and emission through a 590/40 filter (BL2-A).

Data were processed using custom Matlab scripts, deposited at https://github.com/csitron/MATLAB-Programs-for-Flow-Cytometry. In all cases, cells were first gated based on forward scatter (FSC-H) and side scatter (SSC-A) to eliminate cellular debris, as shown in Figure S3A. Cells transfected with CHMP2B variants were selected in the analysis by gating for VL1-A fluorescence above a non-transfected control, as these CHMP2B plasmids additionally encoded an mTagBFP2 transfection marker.

For ratiometric measurements of the ESCRT reporter, the ratio of EGFP to mScarlet-I fluorescence was assessed and a median value was calculated for each technical replicate. The values obtained from technical replicates were then averaged and normalized to the average value from a negative control within each biological replicate, such that all values presented in the figures represent a fold-change relative to this negative control (i.e. NT knockdown control in Figure 3C; untreated in Figures 3D, S3E, and S3F; PBS in Figure S3B; Lpf/NT KD or Lpf/DMSO in Figure 3E).

To measure FRET between mClover3 and mRuby3, a reference no FRET control histogram was first constructed by averaging distributions of BL2-A:BL1-A fluorescence ratios for a no FRET control (e.g. no LLOMe). This histogram was then compared to an experimental condition (e.g. 2 mM LLOMe), and the percentage of the experimental condition histogram that did not overlap with the no FRET control was computed for each technical replicate, as shown in Figure S5A. The technical replicates were then averaged for each biological replicate.

For measurement of mNeonGreen-3K-positive cells in the PFF leakage assay, BL1-A vs BL2-A fluorescence was plotted for each sample. A polygonal gate was then drawn above the xy scatter of a sample known to be mNeonGreen negative (i.e. a sample not treated with PFF-mNeonGreen_11_). The percentage of cells lying in this gate was then calculated for each sample, as shown in Figure S7A.

#### Bioinformatic Analysis of ESCRT-III proteins

Using a human proteome accessed from Uniprot, ESCRT-III protein sequences were aligned with Clustal Omega^164^. Regions homologous to CHMP2B K55-M96 (α2) were then selected for further analysis. The amino acid frequencies in α2 sequences were then computed and compared to the proteome at large. The Kyte Doolittle hydrophobicity was additionally calculated for each position in the α2 sequences and averaged. The code used for this analysis was deposited at https://github.com/csitron/Amino_acid_composition_and_hydrophobicity_analysis.

#### Statistical Analysis

Statistical analysis and plotting were performed in Graphpad Prism 8 (Dotmatics). Comparisons were made between biological replicates, which we define as experiments performed on different days. Multiple comparisons were analyzed by one-way or two-way ANOVA with a post-hoc Tukey HSD test. P-values below 0.05 were considered significant.

