## Supplementary material for "α-Synuclein aggregates inhibit ESCRT-III through sequestration and collateral degradation": Key Resources Table

| REAGENT or RESOURCE | SOURCE | IDENTIFIER |
| --- | --- | --- |
| Antibodies | | |
| CHMP2A | Proteintech | Cat#10477-1-AP;RRID: AB_2079470 |
| CHMP2B | Abcam | Cat#ab33174;RRID: AB_2079471 |
| CHMP2B | Proteintech | Cat#12527-1-AP;RRID: AB_10603358 |
| CHMP3 | Santa Cruz | Cat#sc-166361;RRID: AB_2217111 |
| CHMP4B | Proteintech | Cat#13683-1-AP;RRID: AB_2877971 |
| CHMP6 | Proteintech | Cat#16278-1-AP;RRID: AB_2079498 |
| α-synuclein (MJFR1) | Abcam | Cat#ab138501;RRID: AB_2537217 |
| α-synuclein (LB509) | Abcam | Cat#ab27766;RRID: AB_727020 |
| α-synuclein phosphoS129 (81A) | Abcam | Cat#ab184674;RRID: AB_2819037 |
| α-synuclein phosphoS129 (EP1536Y) | Abcam | Cat#ab51253;RRID: AB_869973 |
| MAP2 | Merck Millipore | Cat#AB5543;RRID: AB_571049 |
| HA | Santa Cruz | Cat#sc-7392;RRID: AB_627809 |
| β-Amyloid | Biolegend | Cat#803001;RRID: AB_2564652 |
| p62 | Abcam | Cat#ab56416;RRID: AB_945626 |
| K48 ubiquitin | Merck Millipore | Cat#05-1307;RRID: AB_11213655 |
| EEA1 | Abcam | Cat#ab2900;RRID: AB_2262056 |
| LAMP2 | Santa Cruz | Cat#sc-18822;RRID: AB_626858 |
| LAMP1 | Developmental Studies Hybridoma Bank (DSHB) | Cat#H4A3;RRID: AB_2296838 |
| Ubiquitin | Santa Cruz | Cat#sc-8017;RRID: AB_628423 |
| GFP | Proteintech | Cat#3h9-150;RRID: AB_10773374 |
| β-actin | Abcam | Cat#ab6276;RRID: AB_2223210 |
| GAPDH | Merck Millipore | Cat#MAB374;RRID: AB_2107445 |
| HSP60 | Abcam | Cat#ab59458;RRID: AB_942025 |
| Calreticulin | Cell Signaling Technologies | Cat#12238S;RRID: AB_2688013 |
| Calnexin | Santa Cruz | Cat#sc-46669;RRID: AB_626784 |
| α-tubulin | Sigma Aldrich | Cat#T6199;RRID: AB_477583 |
| Anti-rabbit IgG (H+L), F(ab')2 Fragment Alexa Fluor 488 | Cell Signaling Technologies | Cat#4412S;RRID: AB_1904025 |
| Goat anti-Mouse IgG (H+L) Cross-Adsorbed Secondary Antibody, Alexa Fluor 488 | Thermo Fisher Scientific | Cat#A-11001;RRID: AB_2534069 |
| F(ab')2-Goat anti-Mouse IgG (H+L) Cross-Adsorbed Secondary Antibody, Alexa Fluor 568 | Thermo Fisher Scientific | Cat#A-11019;RRID: AB_143162 |
| Goat anti-Chicken IgY (H+L) Secondary Antibody, Alexa Fluor 647 | Thermo Fisher Scientific | Cat#A-21449;RRID: AB_2535866 |
| Goat anti-Mouse IgG (H+L) Highly Cross-Adsorbed Secondary Antibody, Alexa Fluor Plus 647 | Thermo Fisher Scientific | Cat#A-32728;RRID: AB_2633277 |
| Goat anti-Rabbit IgG (H+L) Highly Cross-Adsorbed Secondary Antibody, Alexa Fluor Plus 647 | Thermo Fisher Scientific | Cat#A-32733;RRID: AB_2633282 |
| Goat anti-Rabbit IgG (H+L) Secondary Antibody, Alexa Fluor 405 | Thermo Fisher Scientific | Cat#A-31556;RRID: AB_221605 |
| anti-Mouse HRP | Sigma Aldrich | Cat#A4416;RRID: AB_258167 |
| anti-Rabbit HRP | Sigma Aldrich | Cat#A9169;RRID: AB_258434 |
| anti-Rat HRP | Sigma Aldrich | Cat#A9037;RRID: AB_258429 |
| Veriblot | Abcam | Cat#ab131366;RRID: AB_2892718 |
| Bacterial and virus strains | | |
| BL21(DE3) E Coli | N/A | N/A |
| Rosetta (DE3)pLysS E Coli cells | Merck | Cat#70956 |
| pTetOFF a-syn A53T | This study | pCS1; RRID:Addgene_215372 |
| pEF1a mNeonGreen-3K-B1-10-IRES-mKate2-2A-PuroR | This study | pCS7; RRID:Addgene_ 215377 |
| pLJC5-Tmem192-3xHA | Abu-Remaileh et al.^125^ | RRID:Addgene_102930 |
| Chemicals, peptides, and recombinant proteins | | |
| cOmplete EDTA-free Protease Inhibitor | Roche | Cat#5056489001 |
| cOmplete Mini EDTA-free Protease Inhibitor Cocktail | Roche | Cat#4693159001 |
| Alexa Fluor 647 NHS Ester | Thermo Fisher Scientific | Cat#A20006 |
| LLOMe | Santa Cruz | Cat#sc-285992B |
| Leupeptin | Sigma Aldrich | Cat#11017101001 |
| MLN7243 | Selleckchem | Cat#S8341 |
| Bafilomycin A1 | Invivogen | Cat#tlrl-baf1 |
| Doxycycline | Clontech | Cat#631311 |
| Y-27632 | Biozol | Cat#S1049 |
| PR-619 | Sigma Aldrich | Cat#662141-25MG |
| Bortezomib | LC Laboratories | Cat#B-1408 |
| N-ethylmaleimide | Sigma Aldrich | Cat#E3876-25G |
| Blasticidin S | Thermo Fisher Scientific | Cat#A1113903 |
| G418 | Thermo Fisher Scientific | Cat#10131035 |
| Puromycin | Thermo Fisher Scientific | Cat#A1113803 |
| Terrific Broth | Sigma Aldrich | Cat#T0918 |
| Lipofectamine 3000 | Thermo Fisher Scientific | Cat#L3000008 |
| Fugene 6 | Promega | Cat#E2692 |
| X-tremeGENE HP DNA Transfection Reagent | Sigma Aldrich | Cat#6366244001 |
| B-27 Plus | Thermo Fisher Scientific | Cat#A3582801 |
| Polybrene | Sigma Aldrich | Cat#TR-1003-G |
| Lenti-X concentrator | Takara | Cat#631231 |
| poly-D-lysine | Sigma Adrich | Cat#A-003-E |
| Laminin | Thermo Fisher Scientific | Cat#23017015 |
| Laminin | Bio-techne | Cat#3446-005-01 |
| Matrigel | Corning | Cat#354277 |
| Accutase | StemCell | Cat#7920 |
| N2 | Thermo Fisher Scientific | Cat#17502048 |
| NT3 | Peprotech | Cat#AF-450-03-10 |
| BDNF | Peprotech | Cat#450-02 |
| MEM NEAA | Thermo Fisher Scientific | Cat#11140050 |
| GlutaMAX | Thermo Fisher Scientific | Cat#35050061 |
| 1% L-Glutamine | Thermo Fisher Scientific | Cat#25030081 |
| mTeSR Plus | Stem Cell Technologies | Cat#100-0276 |
| Neurobasal | Thermo Fisher Scientific | Cat#21103049 |
| StemFlex Medium | Gibco | Cat#A3349401 |
| UltraPure 0.5M EDTA, pH 8.0 | Thermo Fisher Scientific | Cat#15575020 |
| ReLeSR | Stem Cell Technologies | Cat#5873 |
| DMEM/F12 | Thermo Fisher Scientific | Cat#11995073 |
| FBS (fetal bovine serum) | Thermo Fisher Scientific | Cat#1027010 |
| Penicillin-Streptomycin | Thermo Fisher Scientific | Cat#15140163 |
| TrypLE Express | Thermo Fisher Scientific | Cat#12605036 |
| NucBlue | Thermo Fisher Scientific | Cat#R37606 |
| Paraformaldehyde (PFA) | Thermo Fisher Scientific | Cat#28908 |
| Dako Fluorescence Mounting Medium | Agilent | Cat#S3023 |
| 4X NuPAGE LDS Sample Buffer | Thermo Fisher Scientific | Cat#NP0007 |
| Immobilon Forte Western HRP Substrate | Merck Millipore | Cat#WBLUF0500 |
| Immobilon Classico Western HRP Substrate | Merck Millipore | Cat#WBLUC0500 |
| Restore Western Blot Stripping Buffer | Thermo Fisher Scientific | Cat#21059 |
| Protein G Dynabeads | Invitrogen | Cat#10007D |
| Ubiquitin pan-selector resin | NanoTag Biotechnologies | Cat#N2510 |
| anti-HA magnetic beads | Thermo Fisher Scientific | Cat#88836 |
| Pierce Magnetic Streptavidin Beads | Thermo Fisher Scientific | Cat#88816 |
| Biotinylated CHMP2B 55-96 α2 peptide | Max Planck Institute of Biochemistry Bioorganic Chemistry & Biophysics Core Facility | Biotin-GSKEACKVLAKQLVHLRKQKTRTFAVSSKVTSMSTQTKVMNSQM-NH_2_ |
| DNAse I | Thermo Fisher Scientific | Cat#EN0521 |
| Cas9-NLS | QB3 MacroLab; UC Berkeley | N/A |
| Cpf1 (recombinant protein) | in-house | N/A |
| α-Synuclein | in-house | N/A |
| CHMP2B | in-house | This paper |
| Benzonase | Max Planck Institute of Biochemistry Core Facility | N/A |
| Benzonase | Merck | Cat#71205-M |
| His-SenP2 protease | Max Planck Institute of Biochemistry Core Facility | N/A |
| DMSO | Thermo Fisher Scientific | Cat#D12345 |
| DEAE sepharose | GE | Cat#17-0709-01 |
| Ni-NTA Resin | Thermo Fisher Scientific | Cat#88221 |
| Lenti-X concentrator | Takara | Cat#631231 |
| PageRuler Prestained Protein Ladder | Thermo Fisher Scientific | Cat#26617 |
| RIPA Buffer | Thermo Fisher Scientific | Cat#89900 |
| Critical commercial assays | | |
| GeneArt Precision gRNA Synthesis Kit | Thermo Fisher Scientific | Cat#A29377 |
| Pierce Rapid Gold BCA Protein Assay Kit | Thermo Fisher Scientific | Cat#A53225 |
| Deposited data | | |
| Flow Cytometry Data | Zenodo | 10.5281/zenodo.14506782 |
| Microscopy Data | Zenodo | 10.5281/zenodo.14506782 |
| Immunoblot Data | Zenodo | 10.5281/zenodo.14506782 |
| Tabular Data | Zenodo | 10.5281/zenodo.14506782 |
| UniProt (accessed 2024-07-30) | https:www.uniprot.org | RRID:SCR_002380 |
| Experimental models: Cell lines | | |
| HEK293T | ATCC | RRID: CVCL_0063 |
| HEK293T a-syn-A53T-mRuby3 a-syn-A53T-mClover3 | This study | RRID: CVCL_D4BV |
| HEK293T TetOff-a-syn-A53T | This study | RRID: CVCL_D4BW |
| HEK293T TetOff-a-syn-A53T mRuby3-Galectin-3 mClover3-Galectin-3 | This study | RRID: CVCL_D4BX |
| HEK293T TetOff-a-syn-A53T EGFP-RNF152-IRES-mScarlet-I | This study | RRID: CVCL_D4BY |
| HEK293T TetOff-a-syn-A53T mNeonGreen-3K-1-10-IRES-mKate2 | This study | RRID: CVCL_D4BZ |
| HEK293T TetOff-a-syn-A53T EGFP-RNF152-IRES-mScarlet-I TMEM192-3xHA | This study | RRID: CVCL_E4JW |
| Lenti-X 293T cells | Takara | Cat#632180; No RRID |
| KOLF2.1J AAVS1-TREG3-NGN2 | Hoyer et al.^129^ | RRID: CVCL_D1J6 |
| KOLF2.1J AAVS1-TRE3G-NGN2 CHMP2B-/- cells | This study | RRID: CVCL_E3Y0 |
| KOLF2.1J AAVS1-TRE3G-NGN2 CHMP2B Q165X/+ cells | This study | RRID: CVCL_E3Y1 |
| KOLF2.1J AAVS1-TRE3G-NGN2 CHMP2B I29V/I29V cells | This study | RRID: CVCL_E3Y2 |
| Experimental models: Organisms/strains | | |
| CD-1 wild-type mouse | N/A | N/A |
| Oligonucleotides | | |
| Negative Control siPOOL | siTOOLs Biotech | Cat#si-C002 |
| CHMP2A siPOOL | siTOOLs Biotech | Cat#si-G020-27243 |
| ON-TARGETplus Non-targeting Pool siRNA | Horizon Discovery | Cat#D-001810-10-05 |
| ON-TARGETplus Human CHMP2A SMARTPool siRNA | Horizon Discovery | Cat#L-020247-01-0005 |
| Ultramer repair templates, gRNAs, and primers for gene editing validation are described in Table S1 | N/A | N/A |
| Recombinant DNA | | |
| pTetOFF a-syn A53T | This study | pCS1; RRID:Addgene_215372 |
| pCMV-EGFP-RNF152-IRES-mScarlet-I | This study | pCS2; RRID:Addgene_215373 |
| pCMV mRuby3-Galectin-3 | This study | pCS3; RRID:Addgene_ 215375 |
| pCMV mClover3-Galectin-3 | This study | pCS4; RRID:Addgene_ 215376 |
| pCMV a-syn-A53T-mRuby3 | This study | pCS5; RRID:Addgene_ 215379 |
| pCMV a-syn-A53T-mClover3 | This study | pCS6; RRID:Addgene_ 215378 |
| pEF1a mNeonGreen-3K-B1-10-IRES-mKate2-2A-PuroR | This study | pCS7; RRID:Addgene_ 215377 |
| pSPcas9(BB)-2A-Puro V2.0 | Addgene | RRID:Addgene_62988 |
| pSPcas9(BB)-2A-Puro V2.0 sgCHMP2B-2 aauucccaaaugaagauggc | This study | pCS8; RRID:Addgene_ 231998 |
| pCMV mTagBFP2 | This study | pCS9; RRID:Addgene_ 231999 |
| pCMV mTagBFP2-2A-CHMP2B | This study | pCS10; RRID:Addgene_ 232000 |
| pCMV mTagBFP2-2A-CHMP2B-Q165X | This study | pCS11; RRID:Addgene_ 232001 |
| pCMV CHMP2B-3XHA | This study | pCS12; RRID:Addgene_ 232002 |
| pCMV CHMP2B-L4D,F5D-3XHA | This study | pCS13; RRID:Addgene_ 232003 |
| pCMV CHMP2B-L4D F5D-d10-52-3xHA (Δα1) | This study | pCS14; RRID:Addgene_ 232004 |
| pCMV CHMP2B-L4D F5D-d55-96-3xHA (Δα2) | This study | pCS15; RRID:Addgene_ 232005 |
| pCMV CHMP2B-L4D F5D-d97-106-3xHA (Δα2/3 loop) | This study | pCS16; RRID:Addgene_ 232006 |
| pCMV CHMP2B-L4D F5D-d106-113-3xHA (Δα3) | This study | pCS17; RRID:Addgene_ 232007 |
| pCMV CHMP2B-L4D F5D-d118-138-3xHA (Δα4) | This study | pCS18; RRID:Addgene_ 232008 |
| pCMV CHMP2B-L4D F5D-d55-138-3xHA (Δα2-4) | This study | pCS19; RRID:Addgene_ 232009 |
| pCMV CHMP2B-L4D F5D-d159-174-3xHA (Δα5) | This study | pCS20; RRID:Addgene_ 232010 |
| pCMV CHMP2B-L4D F5D-d201-213-HA (ΔVPS4 binding) | This study | pCS21; RRID:Addgene_ 232011 |
| pEF1a FLAG-CHMP2B | This study | pCS22; RRID:Addgene_ 232012 |
| pEF1a FLAG-CHMP2B-d55-96 | This study | pCS23; RRID:Addgene_ 232013 |
| pCMV sfGFP-CHMP2B-55-96 | This study | pCS24; RRID:Addgene_ 232014 |
| pCMV sfGFP-CHMP2B-3x-55-96 | This study | pCS25; RRID:Addgene_ 232015 |
| pCMV sfGFP-CHMP2B-201-213 | This study | pCS26; RRID:Addgene_ 232016 |
| pT7-7 a-syn A53T | Rospigliosi et al.^156^ | RRID:Addgene_105727 |
| pT7-7 a-syn A53T D115A mNeonGreen-3K-B11 | This study | pCS27; RRID:Addgene_ 232017 |
| pET28 His-SUMO-CHMP2B | This study | pCS28; RRID:Addgene_ 232018 |
| pMD2.G | Addgene; gift of Dider Trono | RRID:Addgene_12259 |
| psPAX2 | Addgene; gift of Dider Trono | RRID:Addgene_12260 |
| pET-Sac-Abeta(M1-42) | Linse^157^ | RRID:Addgene_71875 |
| pLJC5-Tmem192-3xHA | Abu-Remaileh et al.^125^ | RRID:Addgene_102930 |
| pDEST HisAsCpf1 | Kraus et al.^159^ | N/A |
| Software and algorithms | | |
| Fiji 1.54F | https://fiji.sc | RRID:SCR_002285 |
| Matlab R2020b | <http://www.mathworks.com/products/matlab/> | RRID:SCR_001622 |
| Graphpad Prism 8 | <http://www.graphpad.com/> | RRID:SCR_002798 |
| ZEN 2.6 | https://www.zeiss.com/microscopy/en/products/software/zeiss-zen.html | RRID:SCR_013672 |
| MATLAB Flow Cytometry Analysis | https://github.com/csitron/MATLAB-Programs-for-Flow-Cytometry | RRID:SCR_026125; 10.5281/zenodo.14411900 |
| MATLAB Immunoblot Quantification | https://github.com/csitron/Western-Blot-Quantification-in-MATLAB | RRID:SCR_026124; 10.5281/zenodo.14411929 |
| Microscopy lif Channel Quant MATLAB | https://github.com/csitron/Microscopy_lif_Channel_Quant_MATLAB | RRID:SCR_026188; 10.5281/zenodo.14411941 |
| Microscopy lif Adjust MATLAB | https://github.com/csitron/Microscopy_lif_Adjust_MATLAB | [RRID:SCR_026187; 10.5281/zenodo.14411935](https://doi.org/10.5281/zenodo.14411935) |
| Amino acid composition and hydrophobicity analysis | https://github.com/csitron/Amino_acid_composition_and_hydrophobicity_analysis | Awaiting RRID; 10.5281/zenodo.14637057 |
| Leica Applications Suite X 3.5.7.23225 | https://www.leica-microsystems.com/products/microscope-software/p/leica-las-x-ls/ | RRID:SCR_013673 |
| FEI MAPS 2.1 and 3.8 | <http://www.fei.co.jp/_documents/CorrSightDatasheet.pdf> | RRID:SCR_018738 |
| NIS-Elements 5.21.03 | <https://www.nikoninstruments.com/Products/Software> | RRID:SCR_014329 |
| Attune^TM^ Cytometric Software 5.1.2111.1 | https://www.thermofisher.com/de/en/home/life-science/cell-analysis/flow-cytometry/flow-cytometers/attune-nxt-flow-cytometer/software.html | N/A |
| Cytiva Amersham ImageQuant 800 Control Software | https://www.cytivalifesciences.com/en/us/shop/protein-analysis/molecular-imaging-for-proteins/imaging-systems/amersham-imagequant-800-systems-p-11546?srsltid=AfmBOoqGOKRP31R6BqnSdqFpfeJZnjfSKA6yKOY43t3byI4xX30yImGn | N/A |
| Clustal omega | <https://www.ebi.ac.uk/jdispatcher/msa/clustalo>; Madeira et al.^164^ | RRID:SCR_001591 |
| Other | | |
| Superdex 200 Column | GE | Cat#28989335 |
| MonoQ HR 16/10 20 mL Column | Amersham | Cat#17-0506-01 |
| HisTrap Columns | Cytiva | Cat#17524802 |
| Superdex 75 Column | GE | Cat#GE17-5174-01 |
| Hitrap Heparin HP Column | Sigma-Aldrich | Cat#17-0407-03 |
| 50 kDa MWCO Centriprep YM-50 column | Merck | Cat#4311 |
| Nanodrop One | Thermo Fisher Scientific | [https://www.thermofisher.com/eg/en/home/industrial/spectroscopy-elemental-isotope-analysis/molecular-spectroscopy/uv-vis-spectrophotometry/instruments/nanodrop/instruments/nanodro-one.html; RRID:SCR_021242](https://www.thermofisher.com/eg/en/home/industrial/spectroscopy-elemental-isotope-analysis/molecular-spectroscopy/uv-vis-spectrophotometry/instruments/nanodrop/instruments/nanodro-one.html;%20RRID:SCR_021242) |
| BioRuptor Plus | Diagenode | Cat#B01020001; RRID:SCR_023470 |
| FEI CorrSight | FEI | https://www.maastrichtuniversity.nl/sites/default/files/2023-03/corrsight_-_product_flyer.pdf |
| Leica SP8 Falcon | Leica | [https://www.leica-microsystems.com/products/confocal-microscopes/p/leica-tcs-sp8/; RRID:SCR_018169](https://www.leica-microsystems.com/products/confocal-microscopes/p/leica-tcs-sp8/;%20RRID:SCR_018169) |
| Zeiss LSM800 with Airyscan | Zeiss | [https://engineering.unl.edu/nercf/zeiss-lsm-800-airyscan/; RRID:SCR_015963](https://engineering.unl.edu/nercf/zeiss-lsm-800-airyscan/;%20RRID:SCR_015963) |
| Eclipse Ti-2 with Yokogawa W1 spinning disk confocal | Nikon | [https://www.microscope.healthcare.nikon.com/products/confocal-microscopes/csu-series/csu-w1; RRID:SCR_021242](https://www.microscope.healthcare.nikon.com/products/confocal-microscopes/csu-series/csu-w1;%20RRID:SCR_021242) |
| Amersham ImageQuant 800 biomolecular imager | Cytiva | https://www.cytivalifesciences.com/en/us/shop/protein-analysis/molecular-imaging-for-proteins/imaging-systems/amersham-imagequant-800-systems-p-11546?s_kwcid=AL!14612!3!669373576240!b!!g!!imagequant%20800&dtid=semp_google_20450083429_152697004776&ps_kw=imagequant%20800&extcmp=ppc-se-paid-DM-Research-WB-800-EMEA-CY22045-Global-WB-Imagers-Overall-EMEA-RSA-800-Imagers&gad_source=1&gclid=CjwKCAiA34S7BhAtEiwACZzv4cGl7uIx9LckUE_pBY7BvDVKlfZ-D8KOwO5Quld63or85rKx-EK56xoCc_8QAvD_BwE |
| Attune NxT Flow Cytometer | Thermo Fisher Scientific | [https://www.thermofisher.com/us/en/home/life-science/cell-analysis/flow-cytometry/flow-cytometers/attune-acoustic-focusing-flow-cytometer.html; RRID:SCR_019590](https://www.thermofisher.com/us/en/home/life-science/cell-analysis/flow-cytometry/flow-cytometers/attune-acoustic-focusing-flow-cytometer.html;%20RRID:SCR_019590) |
| Desalting Column | Cytiva | Cat#17508702 |
| Low bind 1.5 mL centrifuge tubes | Eppendorf | Cat#30108116 |
| Econo-Pac Disposable Chromatography Columns | Bio-Rad | Cat#7321010 |
| Poly-L-lysine-coated coverslips | Neuvitro | Cat#GG-12-1.5-PLL |
| Epredia glass slides | Thermo Fisher Scientific | Cat#17294884 |
| NuPAGE 1.5 mm 4-12% Bis-Tris gels | Thermo Fisher Scientific | Cat#NP0323BOX |
| NuPAGE 1 mm 12% Bis-Tris gels | Thermo Fisher Scientific | Cat#NP0342BOX |
| PVDF membrane | Sigma-Adlrich | Cat#3010040001 |
| 0.2 µm pore size cellulose acetate membrane | GE | Cat#10404131 |
| Slot blot vacuum manifold | Hoefer | Cat#PR648 |
