## Supplementary material for "α-Synuclein aggregates inhibit ESCRT-III through sequestration and collateral degradation": Suppplementary Figures and Tables

### SUPPLEMENTAL INFORMATION

#### Supplementary Figures

Sitron, et al. Figure S1

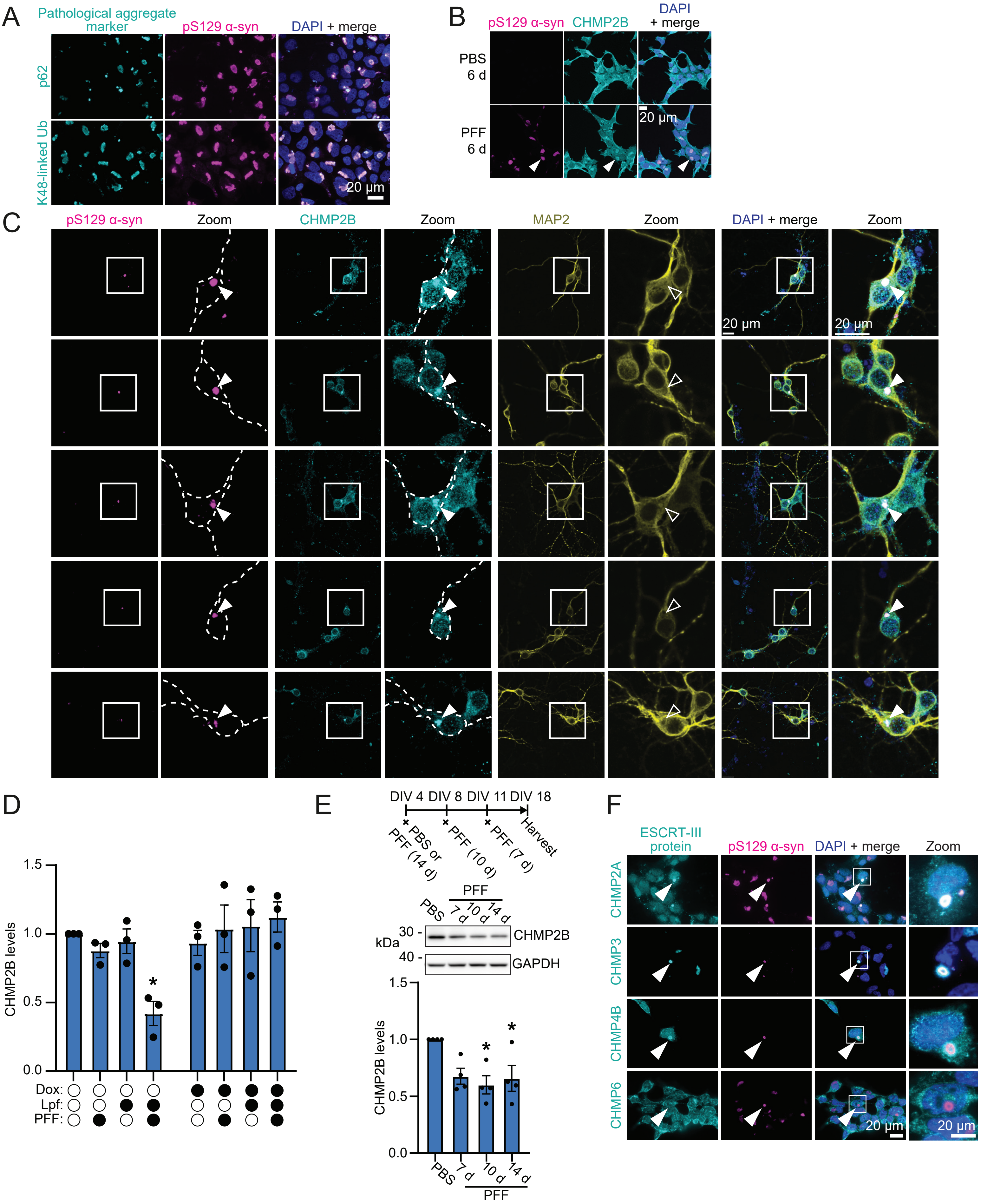

**Figure S1. Co-localization of ESCRT-III proteins with α-syn aggregates, related to Figure 1**

(A) Representative immunofluorescence micrographs of HEK α-syn expressing cells after induction of aggregation using PFFs and Lpf for two days, immunostained for markers of pathologic α-syn aggregation: Ser129-phosphorylated (pS129) α-syn, p62, and K48-linked ubiquitin (Ub; scale bar, 20 μm).

(B) Representative immunofluorescence microscopy of HEK α-syn expressing cells treated with PFFs for 6 days in the absence of Lpf. Cells were stained with anti-CHMP2B and anti-pS129 α-syn antibodies (Scale bar, 20 μm). Arrowheads denote the location of α-syn inclusions.

(C) Additional representative micrographs of CHMP2B-positive pS129 α-syn inclusions in primary neurons treated for 14 days with PFFs, supporting Figure 1D. Cells were stained with antibodies against CHMP2B, pS129 α-syn, and neuronal cytoskeletal marker MAP2 (Scale bar, 20 μm). Arrowheads as in (B).

(D) Densiometric quantification of β-actin-normalized CHMP2B levels for the immunoblot in Figure 1B. Values were additionally normalized to the untreated control. Error bars represent mean ± SEM (n=3). *$p<0.05$ by two-way ANOVA, highlighted comparison between Lpf/PFF condition and untreated control.

(E) Representative anti-CHMP2B and anti-GAPDH immunoblots of lysates from primary neurons, treated with PFFs as described in the schematic above. A densiometric quantification thereof appears below, with CHMP2B levels normalized to the GAPDH loading control and further normalized to the PBS control condition. Error bars signify mean ± SEM (n = 4). *$p<0.05$ by one-way ANOVA, relative to PBS control.

(F) Representative immunofluorescence micrographs of HEK α-syn expressing cells after induction of aggregation, stained with antibodies against pS129 α-syn as well as the ESCRT-III proteins CHMP2A, CHMP3, CHMP4B, or CHMP6 (Scale bar, 20 μm). Arrowheads as in (B).

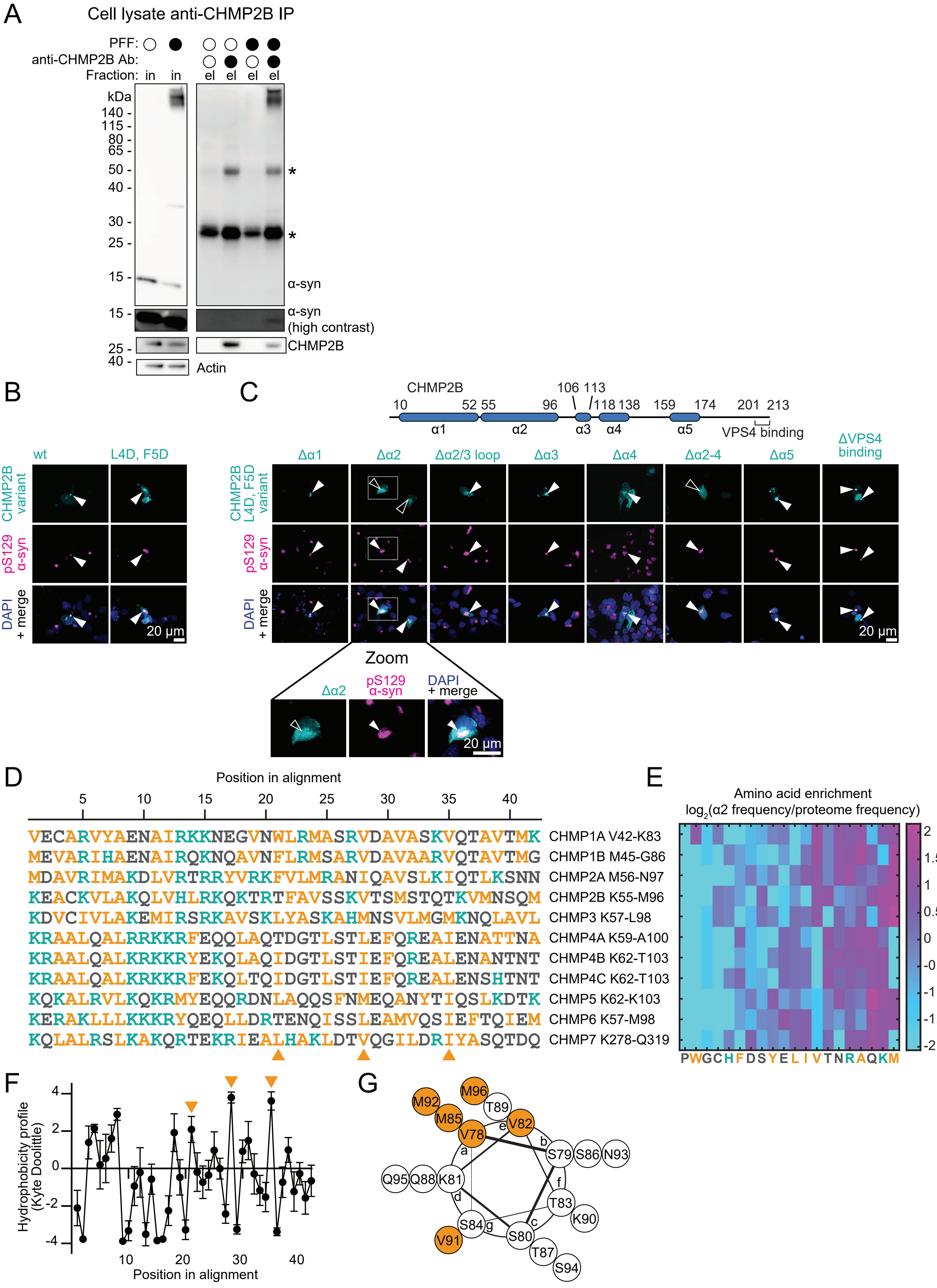
A
Cell lysate anti-CHMP2B IP
PFF:
anti-CHMP2B Ab:
Fraction: in in el el el el
kDa
140 - 115 - 80 - 65 - 50 - 40 - 30 - 25 - 15 -
α-syn
α-syn (high contrast)
CHMP2B
Actin
B
wt
L4D, F5D
CHMP2B variant
pS129 α-syn
DAPI + merge
20 µm
C
CHMP2B
10 52 55 96 106 113 118 138 159 174 201 213
α1 α2 α3 α4 α5 VPS4 binding
Δα1 Δα2 Δα2/3 loop Δα3 Δα4 Δα2-4 Δα5 ΔVPS4 binding
CHMP2B L4D, F5D variant
pS129 α-syn
DAPI + merge
20 µm
Zoom
Δα2
pS129 α-syn
DAPI + merge
20 µm
D
Position in alignment
5 10 15 20 25 30 35 40
VECARVYAENAIRKKNEGVNWLRMASRVDAVASKVQTAVTMK CHMP1A V42-K83
MEVARIHAENAIRQKNQAVNFLRMSARVDAVAARVQTAVTMG CHMP1B M45-G86
MDAVRIMAKDLVRTRRYVRKFVLMRANIQAVSLKIQTLKSNN CHMP2A M56-N97
KEACKVLAKQLVHLRKQKTRTFAVSSKVTSMSTQTKVMNSQM CHMP2B K55-M96
KDVCIVLAKEMIRSRKAVSKLYASKAHMNSVLMGMKNQLAVL CHMP3 K57-L98
KRAALQALRRKKRFEQQLAQTDGTLSTLEFQREAIENATTNA CHMP4A K59-A100
KRAALQALKRKKRYEKQLAQIDGTLSTIEFQREALENANTNT CHMP4B K62-T103
KRAALQALKRKKRFEKQLTQIDGTLSTIEFQREALENSHTNT CHMP4C K62-T103
KQKALRVLKQKRMYEQQRDNLAQQSFNMEQANYTIQSLKDTK CHMP5 K62-K103
KERAKLLLKKKRYQEQLLDRTENQISSLEAMVQSIEFTQIEM CHMP6 K57-M98
KQLALRSLKAKQRTEKRIEALHAKLDTVQGILDRIYASQTDQ CHMP7 K278-Q319
E
Amino acid enrichment
log2(α2 frequency/proteome frequency)
2 1.5 1 0.5 0 -0.5 -1 -1.5 -2
PWGCHFDSYELIVTNRAQKM
F
Hydrophobicity profile (Kyte Doolittle)
4 2 0 -2 -4
10 20 30 40
Position in alignment
G
M96 M92 T89 M85 V82 V78 e b a S79 S86 N93 Q95 Q88 K81 d f T83 K90 S84 g c S80 V91 T87 S94

**Figure S2. Characterization of the interaction between the ESCRT-III protein CHMP2B and aggregated α-syn, related to Figure 2.**

(A) Representative immunoblot of an anti-CHMP2B immunoprecipitation from lysates of HEK α-syn expressing cells after extended treatment with PFFs for 6 days in the absence of Lpf (n=2). The immunoblot was stained with anti-CHMP2B, anti-α-syn, and anti-β-actin (loading control) antibodies. Asterisks denote detection of non-specific bands that elute from the beads used in the immunoprecipitation. The denoted bands arise from Protein G and the anti-CHMP2B antibody. In, input. El, eluate.

(B-C) Analysis of the structural features required for CHMP2B to co-localize with induced α-syn aggregates. HEK α-syn expressing cells were transfected with plasmids encoding indicated CHMP2B mutants tagged with a C-terminal HA-tag and treated to induce α-syn aggregation. Cells were stained with antibodies against pS129 α-syn as well as HA. A schematic of relevant structural features and domain boundaries in CHMP2B is shown above the images in (C) (Scale bar, 20 μm). Arrowheads highlight α-syn aggregates.

(D) Alignment of the predicted α2 region (based on homology to CHMP2B) of human ESCRT-III proteins. The boundaries of the aligned regions are labeled to the right of the alignment. Note that IST1 was excluded due to its divergent sequence. Hydrophobic residues are colored in orange and basic residues are colored in cyan. Orange triangles below the alignment indicate the position of strong, conserved hydrophobic residues placed 7 amino acids apart.

(E) Heatmap of $log_2$-transformed amino acid enrichment, defined as enrichment within the α2 region, displayed and labeled as in (D), divided by the enrichment of that amino acid in the entire human proteome. The color bar adjacent to the plot indicates the values corresponding to the colors in the heatmap.

(F) Kyte Doolittle hydrophobicity profile of the aligned α2 sequences in (D). Data are displayed as mean ± SEM (n=11). Orange triangles are the same as in (D).

(G) Helical wheel diagram of the C-terminal half of $CHMP2B_{\alpha 2}$, emphasizing the presence of a proposed hydrophobic face. Hydrophobic residues are highlighted in orange. Lowercase letters inside the wheel indicate shared radial positions along the alpha-helix.

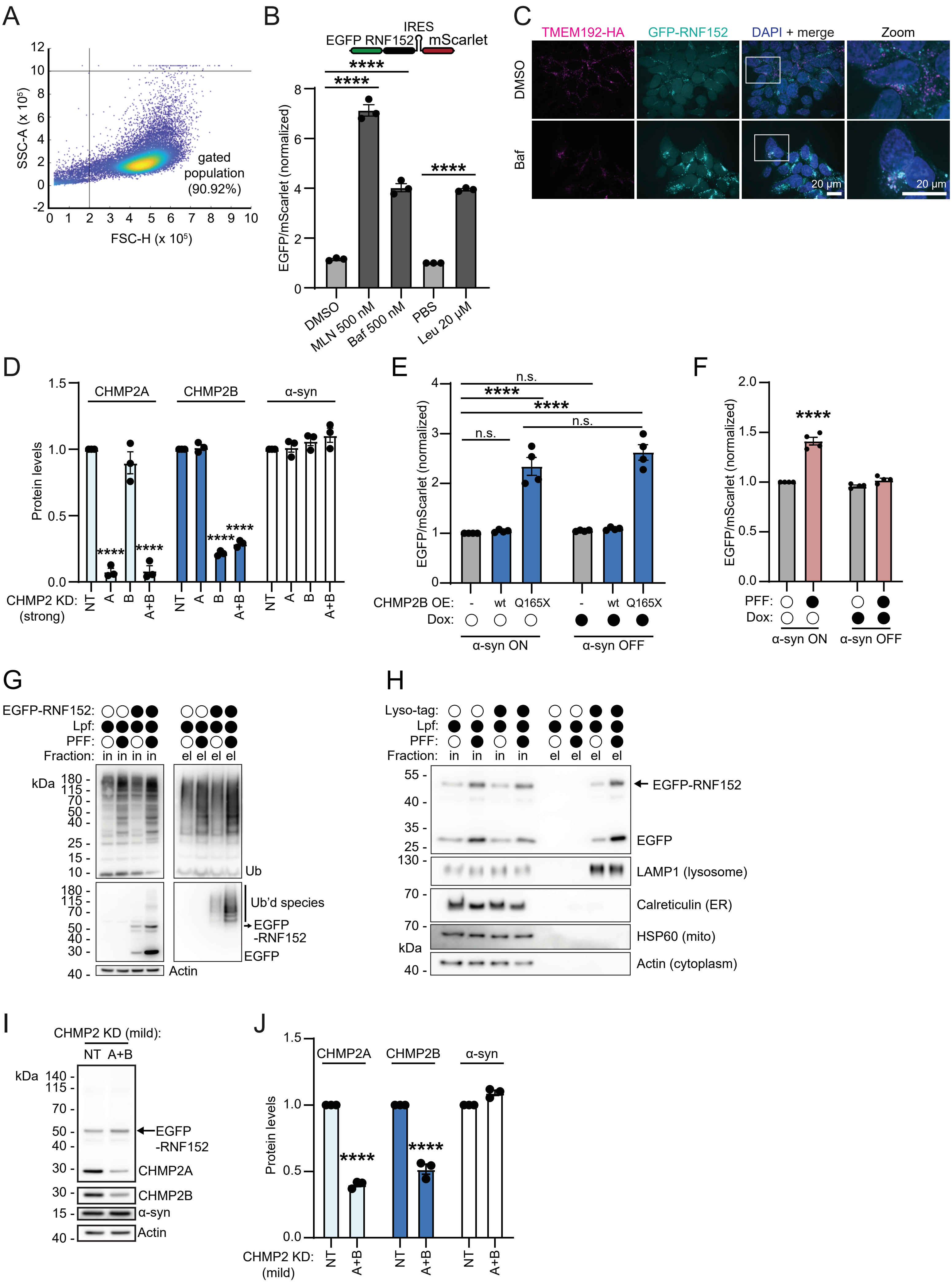

A
SSC-A (x 10⁵)
FSC-H (x 10⁵)
gated population (90.92%)
B
IRES
EGFP RNF152 mScarlet
EGFP/mScarlet (normalized)
DMSO
MLN 500 nM
Baf 500 nM
PBS
Leu 20 μM
C
TMEM192-HA
GFP-RNF152
DAPI + merge
Zoom
DMSO
Baf
20 μm
D
CHMP2A
CHMP2B
α-syn
Protein levels
CHMP2 KD: (strong)
NT A B A+B
E
n.s.
EGFP/mScarlet (normalized)
CHMP2B OE: - wt Q165X
Dox:
α-syn ON
α-syn OFF
F
EGFP/mScarlet (normalized)
PFF:
Dox:
α-syn ON
α-syn OFF
G
EGFP-RNF152:
Lpf:
PFF:
Fraction: in in in in el el el el
kDa
Ub
Ub'd species
EGFP -RNF152
EGFP
Actin
H
Lyso-tag:
Lpf:
PFF:
Fraction: in in in in el el el el
EGFP-RNF152
EGFP
LAMP1 (lysosome)
Calreticulin (ER)
HSP60 (mito)
Actin (cytoplasm)
kDa
I
CHMP2 KD (mild):
NT A+B
kDa
EGFP -RNF152
CHMP2A
CHMP2B
α-syn
Actin
J
CHMP2A
CHMP2B
α-syn
Protein levels
CHMP2 KD: (mild)
NT A+B

**Figure S3. Characterization and perturbation of the RNF152-based ESCRT reporter, related to Figure 3.**

(A) Representative side scatter and forward scatter profile from flow cytometric analysis of the ESCRT reporter cell line. The forward scatter minimum and side scatter maximum values used to gate cells are indicated with vertical and horizontal lines, respectively.

(B) Flow cytometry of ESCRT reporter (schematic above) stability after pharmacological inhibition with the E1 ubiquitin ligase inhibitor MLN7243 (MLN), lysosomal v-ATPase inhibitor bafilomycin A1 (Baf), or lysosomal protease inhibitor leupeptin (Leu). For statistical tests, MLN and Baf are compared to their solvent-only control DMSO while Leu is compared to its solvent control PBS.  DMSO, MLN, and Baf treatment were performed for 6 hr, while PBS and Leu treatment occurred for 18 hr. All values are normalized to the PBS condition. Error bars signify mean ± SEM (n = 3). ****$p<0.0001$ by one-way ANOVA.

(C) Representative immunofluorescence micrographs showing localization of the RNF152 ESCRT reporter relative to lysosomes with and without stabilizing 500 nM bafilomycin A1 treatment for 8 hr. Lysosomes are marked by staining the co-expressed lysosomal marker TMEM192-HA with an anti-HA antibody (Scale bar, 20 μm). A zoomed image of the boxed area appears on the right.

(D) Densiometric quantification of the CHMP2A, CHMP2B, and α-syn signals in the immunoblot in Figure 3B. Values were normalized to the loading control β-actin, then to NT (non-targeting control knockdown). Error bars represent mean ± SEM (n=3). ****$p<0.0001$ by one-way ANOVA, comparing the indicated condition to NT control.

(E) Flow cytometric stability measurements of the ESCRT reporter after transfection of plasmids expressing indicated CHMP2B variants as well as Dox treatment. Values are normalized to the untreated condition. Error bars signify mean ± SEM (n=4). ****$p<0.0001$; n.s. $p>0.05$ by two-way ANOVA. OE, overexpression.

(F) ESCRT reporter flow cytometry after triggering α-syn aggregation without Lpf, using an extended 6-day PFF treatment. Dox treatment is used to silence α-syn expression and thereby control for the effect of PFFs, separate from production of intracellular α-syn aggregates. Normalization as in (E). Error bars represent mean ± SEM (n=4). ****$p<0.0001$ by two-way ANOVA.

(G) Representative immunoblots of anti-ubiquitin (Ub) pulldowns from EGFP-RNF152 ESCRT reporter-expressing lysates with and without aggregation induction. Lysates from cells not expressing the reporter are included as a control to ensure that any immunoprecipitated material recognized by an anti-GFP antibody comes from the reporter. Immunoblots were stained with anti-Ub, anti-GFP, and anti-β-actin. β-actin served as a loading control (n=2).

(H) Representative immunoblots of lyso-IP from EGFP-RNF152 ESCRT reporter-expressing lysates with and without aggregation induction. As a control for nonspecific bead binding, lyso-IP was also performed on lysates lacking expression of the TMEM192-HA "lyso-tag." The reporter was detected with anti-GFP while anti-LAMP1, anti-Calreticulin, anti-HSP60, and anti-β-actin detected the indicated cellular compartments (n=3).

(I) Representative immunoblots to quantify the mild CHMP2A/B knockdown protocol used in Figure 3E to accomplish a weak CHMP2 paralog knockdown in ESCRT reporter HEK cells. Immunoblots were stained with anti-GFP, anti-CHMP2A, anti-CHMP2B, anti-α-syn, and anti-β-actin, with β-actin serving as a loading control (n=3). The anti-CHMP2A immunoblot was re-probed to acquire the anti-GFP signal.

(J) Densiometric quantification of CHMP2A, CHMP2B, and α-syn from the immunoblot in (I). Each value was normalized to β-actin, the loading control, then further normalized to NT. Error bars represent mean ± SEM (n=3). ****$p<0.0001$ by two-way ANOVA, comparisons shown between CHMP2A/B knockdown condition and NT control.

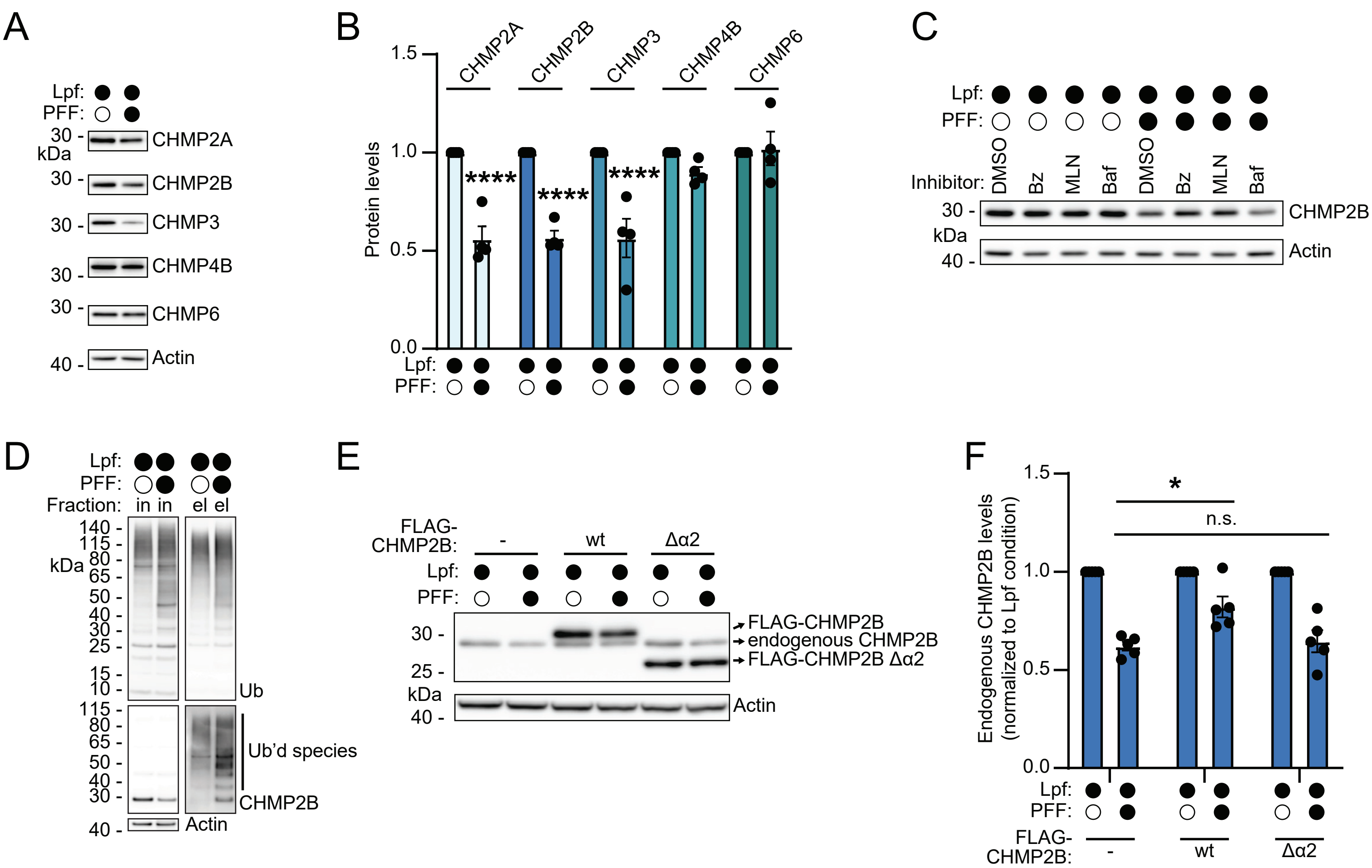
A
Lpf:
PFF:
CHMP2A
CHMP2B
CHMP3
CHMP4B
CHMP6
Actin
kDa
B
CHMP2A
CHMP2B
CHMP3
CHMP4B
CHMP6
Protein levels
****
****
****
Lpf:
PFF:
C
Lpf:
PFF:
Inhibitor:
DMSO
Bz
MLN
Baf
DMSO
Bz
MLN
Baf
CHMP2B
Actin
kDa
D
Lpf:
PFF:
Fraction:
in in
el el
kDa
Ub
Ub'd species
CHMP2B
Actin
E
FLAG-CHMP2B:
-
wt
Δα2
Lpf:
PFF:
FLAG-CHMP2B
endogenous CHMP2B
FLAG-CHMP2B Δα2
Actin
kDa
F
*
n.s.
Endogenous CHMP2B levels (normalized to Lpf condition)
Lpf:
PFF:
FLAG-CHMP2B:
-
wt
Δα2

**Figure S4. Dependence of collateral ESCRT-III degradation on the ubiquitin-proteasome system and α2, related to Figure 4.**

(A) Representative immunoblots of indicated ESCRT-III proteins after induction of aggregation in HEK α-syn cells. The immunoblots were stained with anti-CHMP2A, anti-CHMP2B, anti-CHMP3, anti-CHMP4B, anti-CHMP6, and anti-β-actin antibodies, with the latter serving as a loading control (n=4).

(B) Densiometric quantification of indicated ESCRT-III proteins, normalized to β-actin, from the immunoblots in (A). Signals are additionally normalized for each ESCRT-III protein to the value obtained in the Lpf condition. Error bars represent mean ± SEM (n=4). ****$p<0.0001$ by two-way ANOVA, with comparisons shown between Lpf and Lpf/PFF conditions.

(C) Representative immunoblots of cell lysates from HEK α-syn cells, with or without aggregation induction for two days then 8 hr treatment with either DMSO, 500 nM Bortezomib (Bz, a proteasome inhibitor), 500 nM MLN7243 (MLN, an E1 ubiquitin ligase inhibitor), or 250 nM bafilomycin A1 (Baf, an inhibitor of lysosome acidification). Immunoblots were stained with anti-CHMP2B and anti-β-actin, with the latter detecting the loading control (n=3). A quantification appears in Figure 4C.

(D) Representative immunoblot of anti-ubiquitin immunoprecipitations from HEK α-syn cells with and without aggregation induction. Immunoblots were stained with antibodies against ubiquitin (Ub), CHMP2B, and β-actin (loading control; n=2). Species consistent with ubiquitinated CHMP2B are highlighted with the label “Ub’d species.” In, input. El, eluate.

(E) Representative immunoblots of HEK α-syn expressing cell lysates, which were first transfected with indicated N-terminally FLAG-tagged CHMP2B constructs and then treated to induce α-syn aggregation. Note that cells express both transfected and endogenous CHMP2B. Antibodies against CHMP2B and β-actin (loading control) stained the immunoblots (n=5). A quantification of FLAG-CHMP2B signal in this immunoblot appears in Figure 4D.

(F) Densiometric quantification of endogenous CHMP2B signal from the immunoblot in (E). CHMP2B signal was normalized to β-actin and then internally within each transfection to the Lpf control condition. Error bars signify mean ± SEM (n=5). *$p<0.05$ by two-way ANOVA.

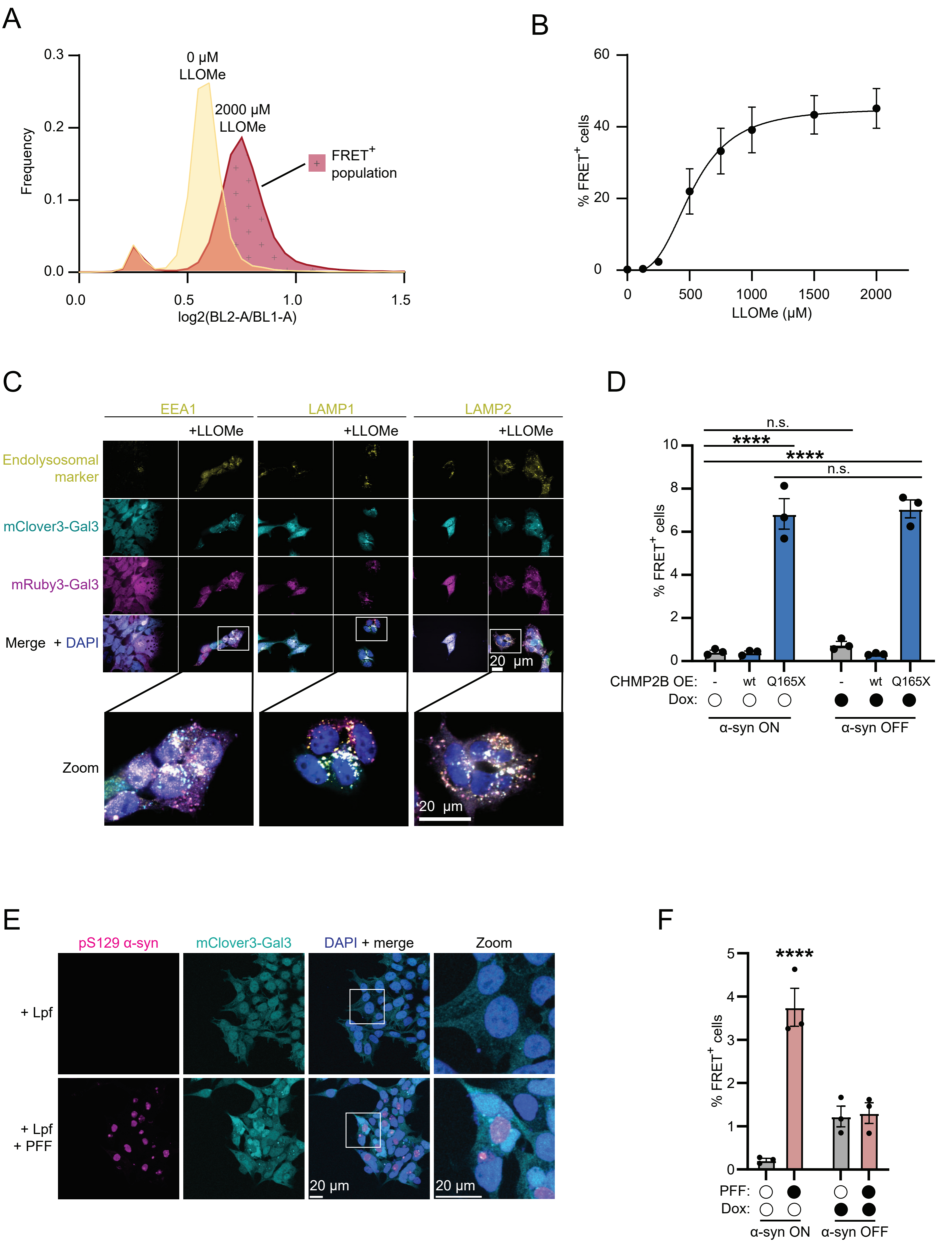
A
0.3
0.2
0.1
0.0
Frequency
0 μM LLOMe
2000 μM LLOMe
FRET+ population
0.0
0.5
1.0
1.5
log2(BL2-A/BL1-A)
B
60
40
20
0
% FRET+ cells
0
500
1000
1500
2000
LLOMe (μM)
C
EEA1
LAMP1
LAMP2
+LLOMe
Endolysosomal marker
mClover3-Gal3
mRuby3-Gal3
Merge + DAPI
Zoom
20 μm
D
n.s.
****
****
n.s.
10
8
6
4
2
0
% FRET+ cells
CHMP2B OE: - wt Q165X - wt Q165X
Dox:
α-syn ON
α-syn OFF
E
pS129 α-syn
mClover3-Gal3
DAPI + merge
Zoom
+ Lpf
+ Lpf + PFF
20 μm
F
****
5
4
3
2
1
0
% FRET+ cells
PFF:
Dox:
α-syn ON
α-syn OFF

**Figure S5. A Gal3 FRET assay measures damage to endolysosomes, related to Figure 5.**

(A) Representative flow cytometry histograms of the ratio between FRET acceptor fluorescence (BL2-A channel) over FRET donor fluorescence (BL1-A) in Gal3 FRET cells with and without 2000 μM LLOMe treatment. The non-overlapping portion of the right-shifted histogram from the LLOMe-treated sample is defined as the FRET-positive population.

(B) Validation of the Gal3 FRET system by flow cytometric analysis of Gal3 FRET cells exposed to increasing concentrations of the lysosome-damaging drug L-Leucyl-L-Leucine methyl ester (LLOMe). Error bars signify mean ± SEM (n=3).

(C) Representative immunofluorescence microscopy of Gal3 FRET cells treated with LLOMe. Cells were stained with antibodies against the early endosome marker EEA1 and the late endosome/lysosome markers LAMP1 and LAMP2 to compare the position of Gal3-fluorescent puncta with endolysosomes (Scale bar, 20 μm). A zoomed image of the boxed area in the +LLOMe merged images is presented below.

(D) Gal3 FRET flow cytometry of cells transfected with plasmids expressing indicated CHMP2B variants as well as treatment with Dox to silence intracellular α-syn expression. Error bars represent mean ± SEM (n=3). ****$p<0.0001$; n.s. $p>0.05$ by two-way ANOVA.

(E) Representative immunofluorescence micrographs of Gal3 FRET cells with and without α-syn aggregate induction. Cells were additionally stained using anti-pS129 α-syn to indicate α-syn aggregates.

(F) Flow cytometry of GAL3 FRET cells treated with PFFs (without the use of Lpf) for 6 days. Cells were additionally grown in Dox to shut off α-syn expression and analyze the effect of PFF treatment alone, in the absence of α-syn aggregate production. Error bars signify mean ± SEM (n=3). ****$p<0.0001$ relative to untreated control by two-way ANOVA. Only significant comparisons shown.

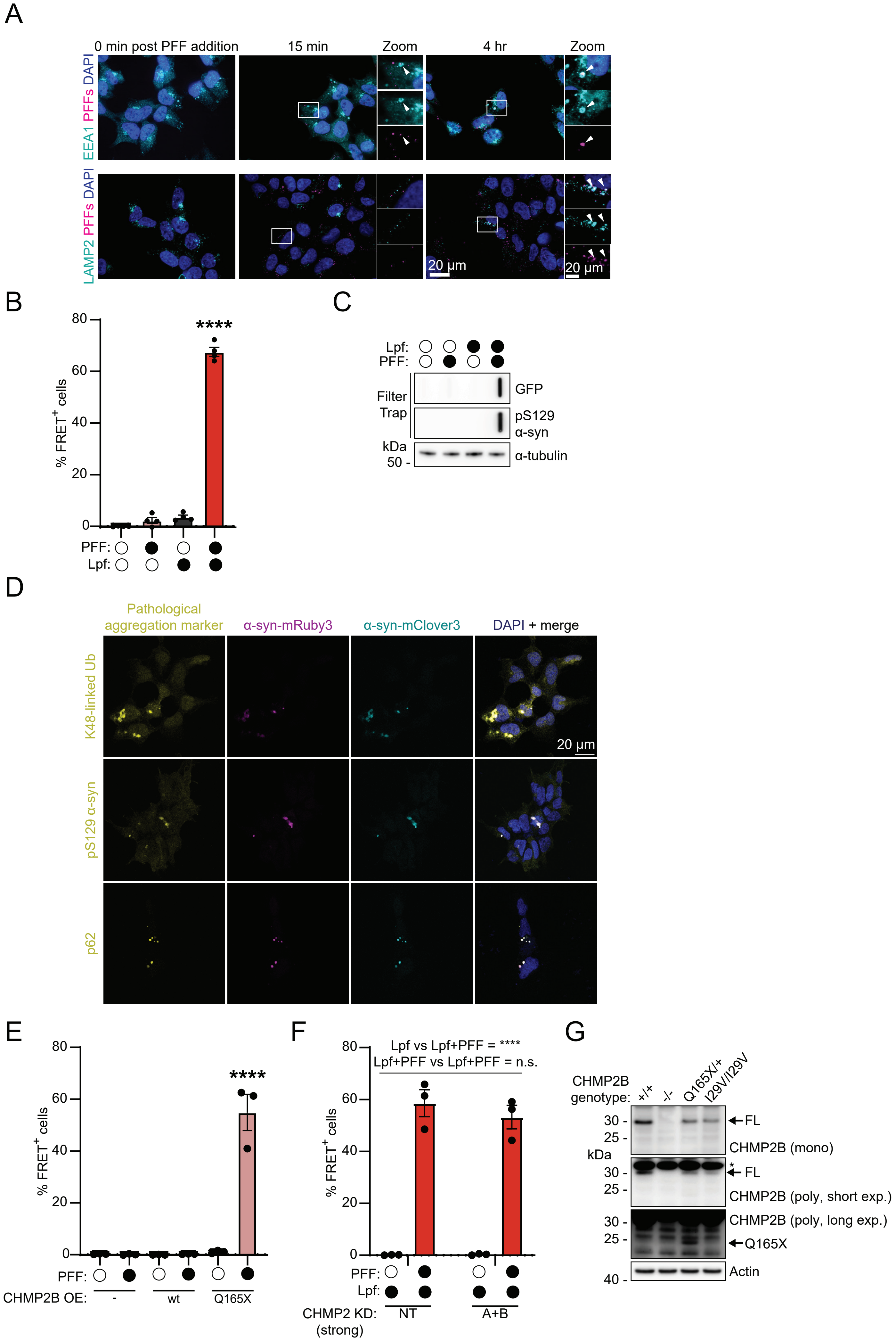
A
0 min post PFF addition
15 min
Zoom
4 hr
Zoom
EEA1 PFFs DAPI
LAMP2 PFFs DAPI
20 μm
20 μm
B
% FRET+ cells
80
60
40
20
0
****
PFF:
Lpf:
C
Lpf:
PFF:
Filter Trap
GFP
pS129 α-syn
kDa
50 -
α-tubulin
D
Pathological aggregation marker
α-syn-mRuby3
α-syn-mClover3
DAPI + merge
K48-linked Ub
pS129 α-syn
p62
20 μm
E
% FRET+ cells
****
PFF:
CHMP2B OE:
-
wt
Q165X
F
Lpf vs Lpf+PFF = ****
Lpf+PFF vs Lpf+PFF = n.s.
% FRET+ cells
PFF:
Lpf:
CHMP2 KD: (strong)
NT
A+B
G
CHMP2B genotype:
+/+
-/-
Q165X/+
I29V/I29V
kDa
30 -
25 -
40 -
FL
CHMP2B (mono)
*
CHMP2B (poly, short exp.)
CHMP2B (poly, long exp.)
Q165X
Actin

**Figure S6. Measurement of seeded α-syn aggregation, related to Figure 6.**

(A) Representative immunofluorescence microscopy of HEK cells treated with Alexa 647 dye-labeled PFFs for indicated durations. Cells were stained with anti-EEA1 and anti-LAMP2 antibodies to highlight early endosomes and late endosomes/lysosomes, respectively (Scale bar, 20 μm). Arrowheads indicate Alexa 647 PFF-positive puncta.

(B) Validation of the HEK α-syn FRET cell line by flow cytometry after treatment with Lpf and PFFs. Error bars represent mean ± SEM (n=3). ****$p<0.0001$ compared to untreated control by one-way ANOVA.

(C) Representative filter trap and immunoblot of lysates from α-syn FRET biosensor cells expressing α-syn-mClover3 and α-syn-mRuby3. The filter was stained with anti-pS129 α-syn and anti-GFP, with anti-GFP detecting α-syn-mClover3. The immunoblot membrane was stained with an anti-α-tubulin antibody to serve as a loading control (n=3).

(D) Representative immunofluorescence microscopy of α-syn FRET biosensor cells after induction of aggregation with Lpf and PFFs. Cells were stained with antibodies against p62, pS129 α-syn, and K48-linked ubiquitin, classical markers of pathologic α-syn aggregation (Scale bar, 20 μm).

(E) α-Syn FRET flow cytometry measuring α-syn seeding in cells transfected with the indicated CHMP2B variants. Error bars signify mean ± SEM (n=3). ****$p<0.0001$ compared to control transfected, untreated condition by two-way ANOVA.

(F) Flow cytometry to highlight the lack of effect of strong double CHMP2A/B knockdown (as in Figures 3B, 3C, and 5B) on PFF-templated α-syn seeding when Lpf is additionally used in α-syn FRET cells. This experiment contrasts with that in Figure 6C, where Lpf was not used. Error bars signify mean ± SEM (n=3). ****$p<0.0001$; n.s. $p>0.05$ by two-way ANOVA.

(G) Representative immunoblot of iPSCs containing the indicated CHMP2B mutations. The membrane was stained with two anti-CHMP2B antibodies and an anti-β-actin antibody, with the latter serving as a loading control (n=2). A polyclonal anti-CHMP2B antibody detected a nonspecific band close to the molecular weight of CHMP2B (denoted with an asterisk), but was capable of detecting CHMP2B Q165X, which the monoclonal antibody did not detect likely due to truncation of its epitope. Mono, monoclonal. Poly, polyclonal.

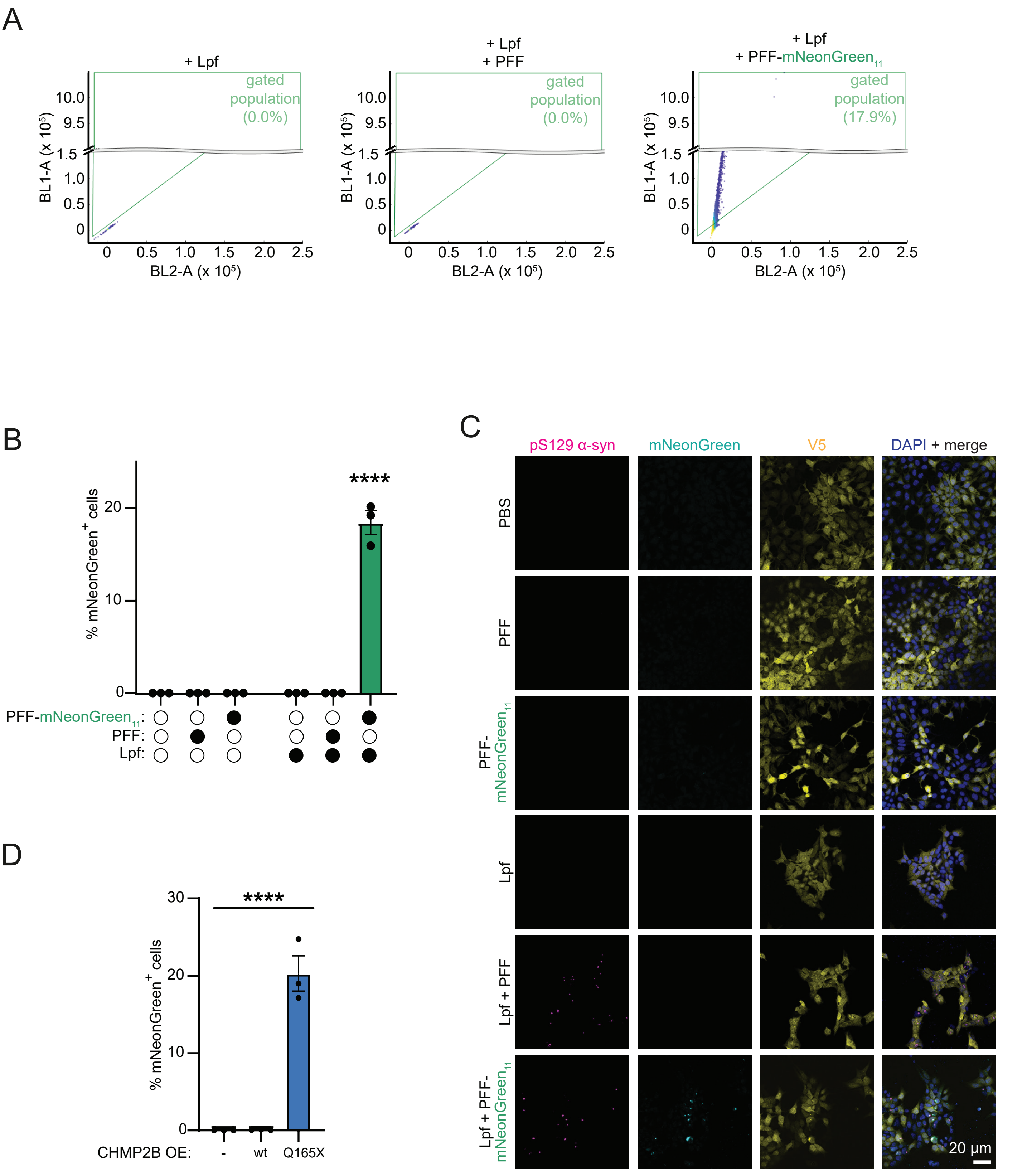
A
+ Lpf
+ Lpf
+ PFF
+ Lpf
+ PFF-mNeonGreen11
gated population (0.0%)
gated population (0.0%)
gated population (17.9%)
BL1-A (x 10^5)
BL2-A (x 10^5)
B
****
% mNeonGreen+ cells
PFF-mNeonGreen11:
PFF:
Lpf:
C
pS129 α-syn
mNeonGreen
V5
DAPI + merge
PBS
PFF
PFF-mNeonGreen11
Lpf
Lpf + PFF
Lpf + PFF-mNeonGreen11
20 μm
D
****
% mNeonGreen+ cells
CHMP2B OE:
-
wt
Q165X

**Figure S7. A split mNeonGreen assay to measure leakage of PFFs into the cytoplasm, related to Figure 6.**

(A) Representative flow cytometric scatter plots demonstrating the gating strategy used to quantify the mNeonGreen-fluorescent population in the split mNeonGreen PFF leakage assay. Cells were treated with Lpf along with either untagged PFFs or PFF-mNeonGreen$_{11}$. Cells were analyzed for their mNeonGreen fluorescence in the BL1-A channel and for their autofluorescence in a neighboring channel, BL2-A. A triangular gate was drawn above the Lpf-alone population to define the mNeonGreen-fluorescent region.

(B) Validation of the split mNeonGreen PFF leakage assay by treatment with Lpf, untagged PFFs, and PFF-mNeonGreen$_{11}$ as in (A). The plot quantifies the percentage of mNeonGreen fluorescent cells measured in flow cytometry. Error bars represent mean ± SEM (n=3). ****$p<0.0001$ compared to untreated control by two-way ANOVA.

(C) Representative immunofluorescence micrographs of the split mNeonGreen PFF leakage assay validation under the same conditions as (B). mNeonGreen signal represents regions of complemented PFF-mNeonGreen$_{11}$ and Cyto-mNeonGreen$_{1-10}$. Cells were stained with anti-pS129 α-syn to mark α-syn aggregates and anti-V5 to label total Cyto-mNeonGreen$_{1-10}$ (Scale bar, 20 μm).

(D) Flow cytometry of the split mNeonGreen PFF leakage assay in cells transfected with plasmids encoding the indicated CHMP2B variants and exposed to PFF-mNeonGreen$_{11}$. Note that this experiment does not feature Lpf treatment. Error bars signify mean ± SEM (n=3). ****$p<0.0001$ relative to control transfected condition by one-way ANOVA.

#### Supplementary Tables

##### Table S1: Information about oligonucleotides used in the creation of CHMP2B mutant iPSC cell lines, related to STAR Methods.

| Oligonucleotide | Source | Sequence | Additional Notes |
|---|---|---|---|
| gRNA for creation of CHMP2B-/- | IDT | GCCAAACAACTTGTGCATCTACGG | For use with Cpf1 |
| gRNA for creation of CHMP2B Q165X/+ | IDT | ATCAAGAACTTGATTCACAATATC | For use with Cpf1 |
| gRNA for creation of CHMP2B I29V/I29V | IDT | AGAGTTACGAGGTACACAGA | For use with Cas9 |
| Ultramer homology arms for CHMP2B Q165X | IDT | ccaactaagaaaagatgatgttcatacctttccagaaatttcaattccaatCtcatcaagaacttAattcacaatatcctggctttcttcttcgtcatcagaaccgtcaaagatgtc | |
| Ultramer homology arms for CHMP2B I29V | IDT | ctcctagatgtaataaaggaacagaatcgagagttacgaggtacacagagAgctataGtcagagatcgagcagctttagagaaacaagaaaaacagctggtaagtag | |
| CHMP2B-/- genotyping forward PCR primer | IDT | 5'-AAGAAAATGGCCAAGATTGGTA | |
| CHMP2B-/- genotyping reverse PCR primer | IDT | 5'-CCATCTTCATTTGGGAATTCAT | |
| CHMP2B Q165X genotyping forward PCR primer | IDT | 5'-TTGATGACATCTTTGACGGTTC | |
| CHMP2B Q165X genotyping reverse PCR primer | IDT | 5'-GAAATAAAAACCATGCACCTCC | |
| CHMP2B I29V genotyping forward PCR primer | IDT | 5'-GGTTTCTTTTGTGATTCTCCTAG | |
| CHMP2B I29V genotyping reverse PCR primer | IDT | 5'-CATGTGCCTTCTTCCTAGTTAGC | |
